# Lepidoptera genomics based on 88 chromosomal reference sequences informs population genetic parameters for conservation

**DOI:** 10.1101/2023.04.14.536868

**Authors:** Chiara Bortoluzzi, Charlotte J. Wright, Sangjin Lee, Trevor Cousins, Thiago A. L. Genez, David Thybert, Fergal J. Martin, Leanne Haggerty, The Darwin Tree of Life Project Consortium, Mark Blaxter, Richard Durbin

## Abstract

Butterflies and moths (Lepidoptera) are one of the most ecologically diverse and speciose insect orders, with more than 157,000 described species. However, the abundance and diversity of Lepidoptera are declining worldwide at an alarming rate. As few Lepidoptera are explicitly recognised as at risk globally, the need for conservation is neither mandated nor well-evidenced. Large-scale biodiversity genomics projects that take advantage of the latest developments in long-read sequencing technologies offer a valuable source of information. We here present a comprehensive, reference-free, whole-genome, multiple sequence alignment of 88 species of Lepidoptera. We show that the accuracy and quality of the alignment is influenced by the contiguity of the reference genomes analysed. We explored genomic signatures that might indicate conservation concern in these species. In our dataset, which is largely from Britain, many species, in particular moths, display low heterozygosity and a high level of inbreeding, reflected in medium (0.1 - 1 Mb) and long (> 1 Mb) runs of homozygosity. Many species with low inbreeding display a higher masked load, estimated from the sum of rejected substitution scores at heterozygous sites. Our study shows that the analysis of a single diploid genome in a comparative phylogenetic context can provide relevant genetic information to prioritise species for future conservation investigation, particularly for those with an unknown conservation status.

## Introduction

Biodiversity loss is one of the most urgent problems the world is facing today. Actions for biodiversity conservation and policy change are often informed and catalysed by the International Union for Conservation of Nature’s (IUCN) Red List. The aim of this list is to provide “the most objective, scientifically-based information on the current status of globally threatened biodiversity” ^1^ and the assignment of species to various extinction risk categories is based on a number of objective criteria connected to population size and trend, and geographic range ^2^. While the IUCN Red list is an invaluable tool for the monitoring of species at risk of extinction, it has a taxonomic bias that excludes species with small body sizes, narrow distribution ranges, and low dispersal abilities ^3,4,5^. For instance, although insects comprise over half of known eukaryotic species, the number of threatened and extinct insects on the IUCN Red List is minimal ^6, 7^. This effectively excludes these critical taxa from conservation efforts, which are still concentrated on a few emblematic mega-biota such as mammals and birds ^3, 8^.

Butterflies and moths (Lepidoptera) belong to one of the most speciose insect orders, with more than 157,000 described species belonging to 43 superfamilies and 133 families ^9^. Lepidoptera are the largest single radiation of plant-feeding insects, making them one of the most ecologically diverse insect orders ^9, 10^. They are a vital component of biodiversity and play important roles in ecosystem functioning, for example as pollinators or as sources of food for other organisms ^11, 12^. They also serve as model organisms for research in ecology, physiology, and agriculture, making them one of the most well-studied groups of insects ^9, 12^. Because of these ecological roles, Lepidoptera are often used as sensitive indicators of the health and functioning of ecosystems. Decline in lepidopteran assemblages and populations can thus potentially signal the risks for a collapse of the ecosystems ^6, 12,13,14^. The abundance and diversity of Lepidoptera is declining at an alarming pace, as a result of factors including habitat loss, degradation, and fragmentation, spread of invasive species, and use of harmful substances in intensive crop production ^6, 8^. Long-term monitoring data on butterflies and moths in the United Kingdom highlighted that some criteria used in IUCN Red List assessment are not suitable for all taxa ^7^. Specifically, the use of population trend sizes over the last 10 years (Criterion A of the IUCN Red List) is not sufficiently reliable to enable the accurate classification of insect taxa, because of their short generation times, high temporal variability, and the boom and bust nature of their population dynamics.

Understanding the status and trends in distribution and abundance of species is challenging, especially when species are rare and/or lack long-term monitoring data. Parameters estimated from genome sequence data can contribute to filling this knowledge gap, providing unbiased information for a better assessment of the extinction risk and potential recovery of species ^15^, even when risks are difficult to evaluate from ecological and demographic data alone ^2^. Genomic sequence data can inform conservation concerns because they retain information related to species’ demographic history, and individuals’ genome-wide genetic diversity, level of inbreeding, and total number of deleterious mutations (or genetic load). This can inform decision-making of conservation practitioners ^15,16,17^. Since the release of the domesticated silk moth (*Bombyx mor*i) genome ^18,19,20^, progress in sequencing and assembly technologies has permitted rapid generation of additional lepidopteran genome sequences. Chromosome-level, near-error free, near-gapless genomes can now be sequenced at a relatively low cost and unprecedented rate ^21^. Global and regional sequencing projects, such as the 5,000 insect genomes (i5K) initiative ^22^ and the Darwin Tree of Life Project (DToL) ^23^, are now tackling one of the most important constraints in the application of genomics to biodiversity conservation, the unbiased sampling of species across the Tree of Life. These sequencing projects are making it clear that an approach that goes beyond threatened species can drive understanding of many aspects of preserving threatened species diversity ^24, 25^. Overall, analysis of high-quality reference genomes *via* a comparative genomic approach can be a powerful tool to highlight differences in genomic patterns and parameters and thus quantify evolutionary and demographic forces - including those that reveal conservation threats - that have been acting on the genome. This approach is applicable to all taxa, whatever their recognised conservation status.

In this study, we explore the use and relevance of genome sequence data for conservation. We present a comprehensive, whole-genome multiple sequence alignment for 88 species of Lepidoptera, covering much of its taxonomic diversity. Our study shows that the analysis of single diploid genomes across a range of related species can provide relevant genetic information to prioritise species for future conservation investigation, particularly for those with an unknown conservation status.

## Results

We generated a whole-genome multiple sequence alignment of 88 lepidopteran species with chromosomal, high-contiguity reference genome sequences, covering 155 million years of evolution and 12 of the 43 superfamilies ^26^ (**Figure 1**). The alignment was generated using Cactus, a toolkit which avoids the limitations of basing the alignment on a single focal sequence. The alignment we present here is of high quality. The fraction of the genome of pairs of species that are aligned depended on the phylogenetic distance between them, varying from under 5% to over 70% (**Figure 2a**; **Figure S2**). On average 9.1% (standard deviation - SD: 6.6) of each lepidopteran genome was aligned to another genome in the dataset, although this value ranges extensively from 4.6% (*Lymantria monacha*) to 16.5% (*Autographa gamma*). While most of the 2,451- benchmarking universal single copy orthologs (BUSCO) from the *lepidoptera_odb10* set were consistently aligned by Cactus (**Figure 2b**), we identified 1,054 BUSCOs with inconsistent alignments, six of which were inconsistently aligned in 87 species. Reference genome assemblies that failed to meet all the Vertebrate Genomes Project (VGP) assembly quality metrics (see **Figure 1**) had the highest numbers of inconsistently aligned BUSCO genes, suggesting that the quality of the reference genome assembly is important in the accuracy of the Cactus alignment (**Figure 2b**). Examples of this are *Danaus plexippus* (number of incorrectly mapped BUSCO genes - *n* = 652), *Zerene cesonia* (*n* = 156), and *Spodoptera litura* (*n* = 156). Cactus consistently aligned almost all 2,501-single copy orthogroups identified by OrthoFinder ^27^ within the set of genomes with available Ensembl gene annotations (*n =* 49 species) (**Figure S3a**). In this case, we found just 31 inconsistently aligned orthogroups. We also observe that, in each of the 49 species, over 80% of the protein-coding bases are aligned to at least one other species (**Figure S3b**). We did not observe any clear correlation between the coverage at unique non-overlapping coding sequences and the position of the species in the phylogenetic tree, as variation was also observed among species belonging to the same superfamily. We also did not observe any correlation with the quality of the reference genome assembly, as all 49 species included in the coding coverage analysis were generated by Darwin Tree of Life Project (DToL), meeting all VGP assembly quality metrics.

**Figure 1.**
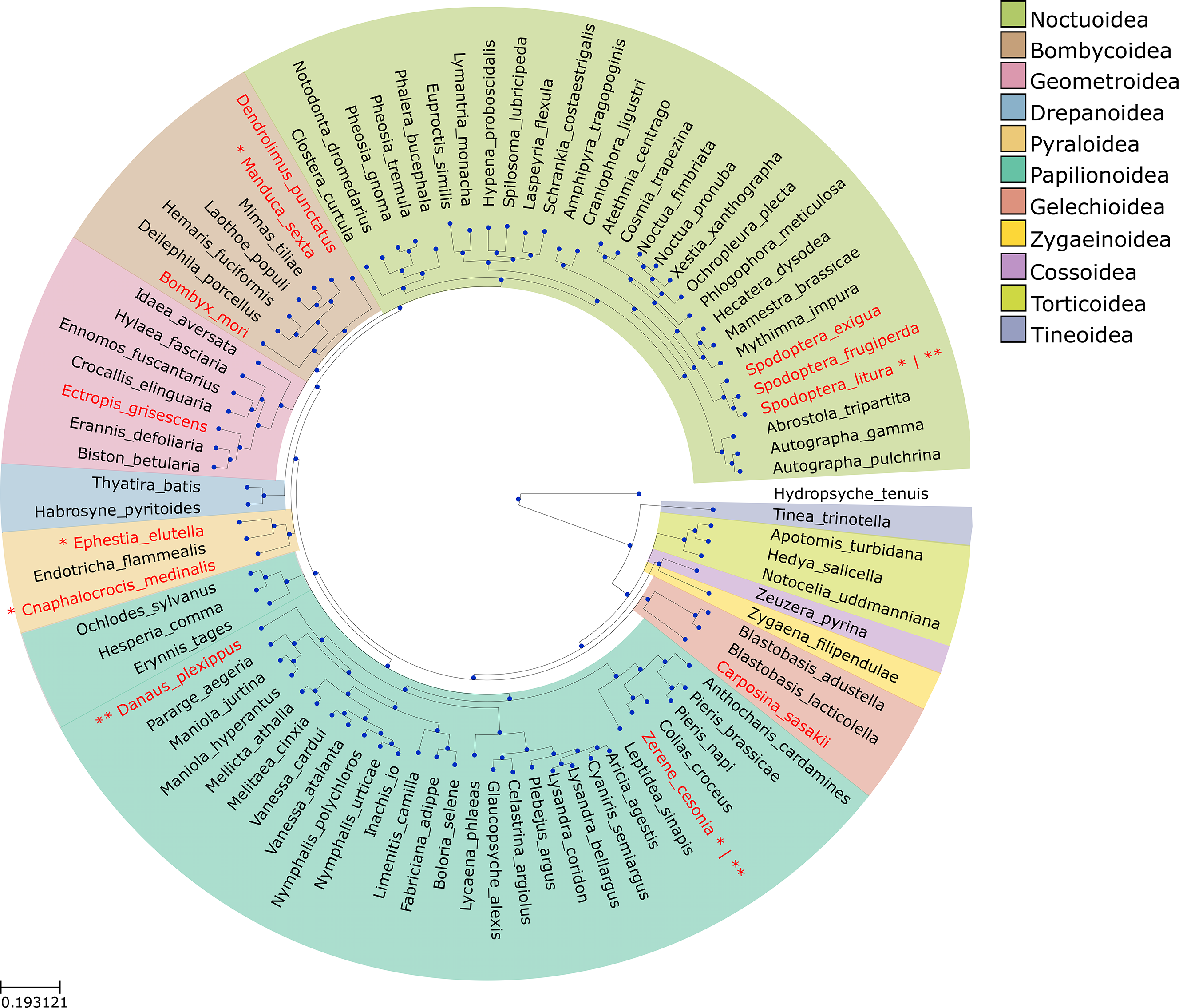
Phylogenetic tree of the 88 Lepidoptera species used as guide tree in the reference-free, whole- genome multiple sequence alignment. The phylogenetic tree includes the 88 Lepidoptera species and the trichopteran outgroup *Hydropsyche tenuis,* which is the only species left in white. Species are coloured based on the superfamily they belong to. Species whose reference genome assembly was downloaded from the International Nucleotide Sequence Database Collaboration (INSDC) have their name in red, while those whose reference genome assembly was generated by the Darwin Tree of Life (DToL) Project have their name in black. All reference genome assemblies included in the Cactus alignment are chromosome-level assemblies with a contig N50 > 1 Mb and a scaffold N50 > 10 Mb, except for five species for which the contig N50 was below 1 Mb (here identified by the one asterisk ‘*’), and three species for which the scaffold N50 was below 10 Mb (here identified by two asterisks ‘**’).

**Figure 2.**
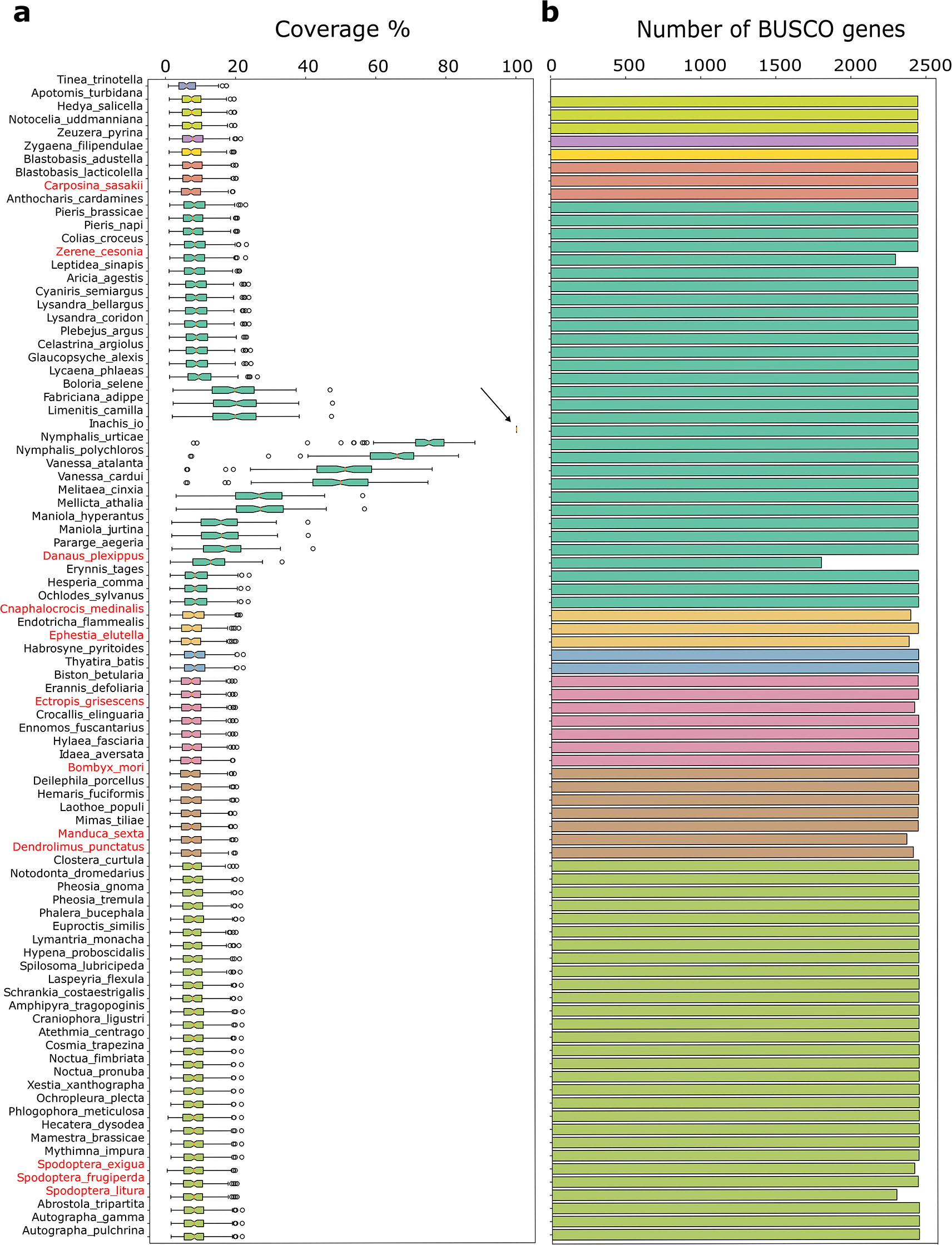
Alignment quality metrics of the 88 Lepidoptera whole-genome multiple sequence alignment. (a) Coverage calculated for each assembly on 100 intervals of 1 Mb, randomly sampled along the genome. Coverage is here expressed as the proportion of sites in species A (in this case *Inachis io*) that align to *n* or more sites in each other species. The coverage of *Inachis io* with itself (100%) is indicated by the arrow. **(b)** Number of correctly aligned (or consistent) benchmarking universal single copy orthologs (BUSCO) (*n =* 2,451). Species whose reference genome was downloaded from the INSDC have their name in red. Species are ordered according to the phylogenetic tree and are coloured based on the superfamily they belong to.

Next, we estimated two metrics of diversity, genome-wide heterozygosity and extent of inbreeding, on each of the 74 reference genomes for which PacBio HiFi data were available. We observed a clear pattern in the genome-wide heterozygosity across non-overlapping 10 kb windows. Species in the superfamily Papilionoidea (the butterflies), show, on average, a higher genome-wide heterozygosity (mean = 5.4e-03, SD = 1.9e-03 heterozygotes per site) than all other superfamilies analysed (mean = 3.9e-03, SD = 1.8e-03 heterozygotes per site) (**Figure 3a**; **Table S2**). Two genomes stand out, because they are almost entirely depleted of heterozygosity: the cabbage moth (*Mamestra brassicae*) (**Figure 3c**) and the centre-barred sallow (*Atethmia centrago*). The average genome-wide heterozygosity in *M. brassicae* is **9.8e-05** (SD: 7.5e-04) heterozygotes per site, which is a 38-fold reduction from the average superfamily (Noctuoidea) heterozygosity, whereas in *A. centrago* it is 8.8e-04 (SD: 2.0e-03) an approximately 4-fold reduction from the average superfamily (Noctuoidea) heterozygosity.

**Figure 3.**
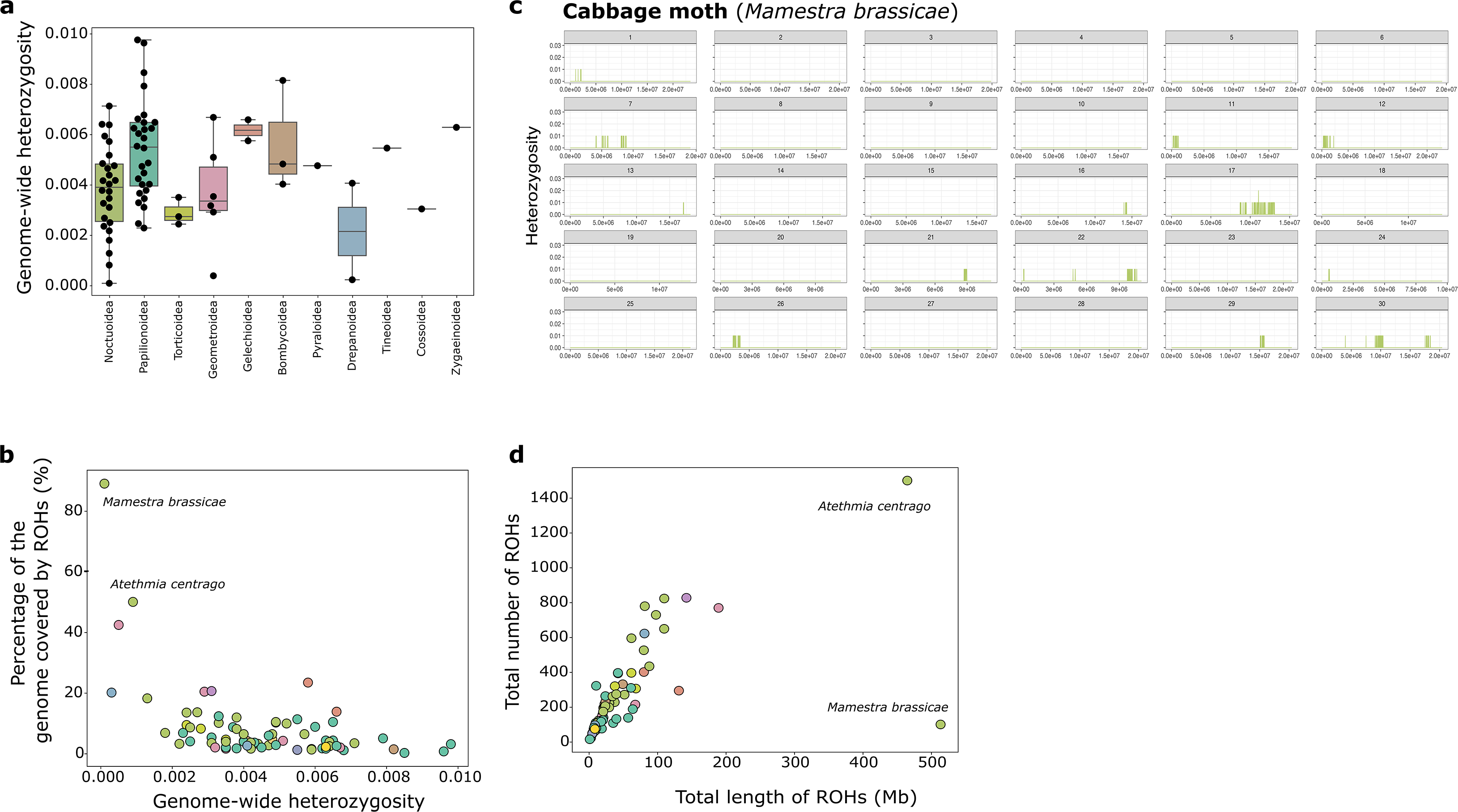
Genome-wide heterozygosity and runs of homozygosity (ROH). (a) Genome-wide heterozygosity expressed as the corrected number of heterozygous sites per base pair (sex chromosomes excluded). Species are grouped and coloured based on the superfamily they belong to. **(b)** Genome-wide heterozygosity versus the percentage of the genome in a run of homozygosity (ROH). **(c)** Genome-wide heterozygosity in each of the 30 autosomes of the cabbage moth, *Mamestra brassicae*. **(d)** Total length of all ROHs (in Mb) versus the total number of ROHs in each Lepidoptera genome.

As expected, genome-wide heterozygosity negatively correlates with the percentage of the genome in a run of homozygosity (ROH) (Pearson’s correlation: -0.51, p-value: 2.4e-06) (**Figure 3b**), as the higher the number of ROHs, the higher the number of bases found in a homozygous stretch (Pearson’s correlation: 0.61, p-value: 5.6e-09) (**Figure 3d**). To investigate whether any species sampled showed any sign of recent inbreeding, we further classified ROHs into three classes based on their total length. We found that, on average, 8.5% of a lepidopteran genome is in a run of homozygosity (**Table S2**). In our dataset, the vast majority of the identified ROHs are short (≤ 0.1 Mb) (**Table S2**; **Figure S4a**), suggesting that they trace to distant common ancestors and are thus a consequence of a historically small population size rather than of recent inbreeding. However, compared to medium (0.1 -1 Mb) and long (≥ 1.0 Mb) ROHs, short ROHs cover a small fraction of the lepidopteran genome (**Table S2**; **Figure S4b**). In two assemblies more than half of the genome is covered by ROHs: *M. brassicae*, where only 101 ROHs cover 89% of the genome (genome size = 576 Mb), and *A. centrago* where 1,501 ROHs cover 50% of the genome (genome size = 926 Mb) (**Table S2**). In *M. brassicae*, while long ROHs are the most abundant class, there are also small patches of heterozygosity, (**Figure 3c**), suggesting sustained recent inbreeding, with some selection for heterozygosity, perhaps at balanced lethal recessive alleles. However, in *A. centrago*, medium ROHs are the most abundant class, followed by long ROHs (**Figure S4**), again suggesting repeated recent inbreeding. We found that genomic diversity varied considerably also among species in different Red List categories, as defined by the butterflies and moths Red List for Great Britain ^28^ (**Table S1**). This variation was mostly observed in species without data (*No data*) and in species classified as *Least concern*. Contrary to expectations, species classified as *Vulnerable*, *Endangered*, and *Regionally extinct* had a higher heterozygosity and lower fraction of the genome covered by ROHs than expected (**Figure 4**). However, most of the samples from these three risk categories were obtained from continental European countries to reduce the impact of sampling on an already impacted population, and so do not reflect genomic diversity within the United Kingdom.

**Figure 4.**
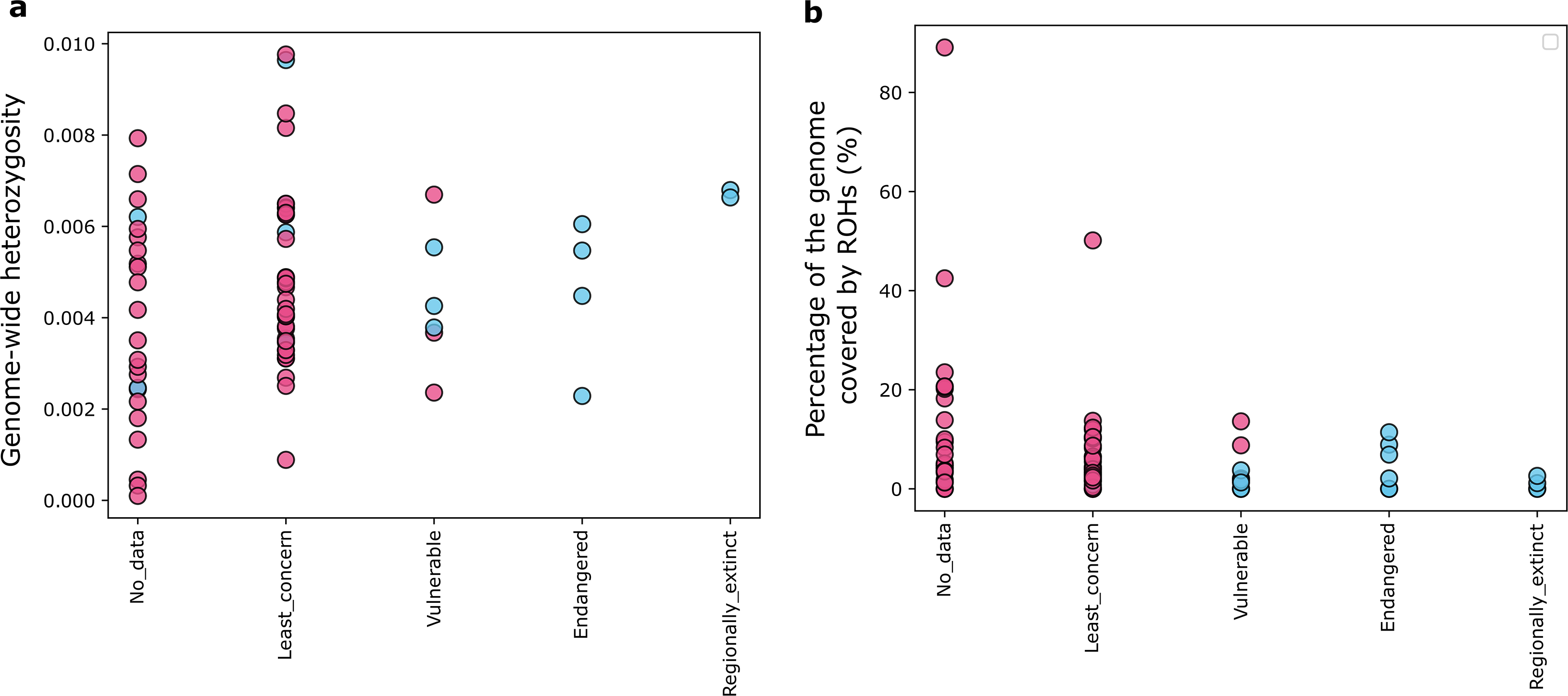
Classification of Lepidoptera in several risk categories according to the Butterfly Conservation for butterflies and moths. (a) Genome-wide heterozygosity of species grouped according to the Great Britain Red List for butterflies and moths. (b) Percentage of the genome covered by ROHs in the different species grouped according to the Great Britain Red List for butterflies and moths. Species are coloured based on their sampling origin.

We next ran a Pairwise Sequentially Markovian Coalescent (PSMC) analysis across each of the 74 reference genome assemblies for which PacBio HiFi data were available. In principle PSMC can be used to estimate long-term population size histories ^29^. The inferred histories suggest that across most species, at ∼100 Kya, the effective population size (*N_e_*) was at around 1e+06, after which it plummets to around 1e+04 by ∼10 Kya (**Figure S5a**). If accurate, the small inferred effective recent population sizes would possibly reflect high levels of recent selection, recent inbreeding, or genuinely low effective panmictic population size over the last few thousand years ^30^. However, we note that PSMC becomes increasingly unreliable when the recombination rate is higher than the mutation rate ^31^, as is the case in Lepidoptera (ratio typically greater than 20:1). Indeed, in simulations with a similar ratio of recombination to mutation as in Lepidoptera, PSMC was unable to accurately recover the simulated effective population size trajectory, suggesting that for Lepidoptera, PSMC inferences are not a reliable indicator of changes in effective population size over time (**Figure S5b**).

Sequences that display a higher than expected level of conservation across multiple and/or distantly related species are likely to be under purifying selection ^32^. Therefore, mutations at these highly conserved genomic sites are likely to have, on average, a negative effect on fitness (i.e. deleterious). Ultimately, these sites can be used to estimate and compare the total burden of deleterious alleles (i.e. genetic load) among individuals^33^. In this study, we used high rejected substitution scores estimated using GERP++ ^34^ to identify likely deleterious mutations at highly conserved sites. On average, we were able to assign a rejected substitution score (or GERP score) to 15.6%, or 74,380,594 sites, of a species genome. However, for downstream analyses we retained only sites with a GERP score ≥0, corresponding to an average of 9.6%, or 45,403,088 sites, of the lepidopteran genome. These sites represent a substitution deficit, which is indicative of sites subjected to purifying selection. We found that in most species these sites cluster at the beginning of the GERP score distribution, supporting that low scores mostly reflect nearly evolving sites (*Fabriciana adippe* is used as a representative in **Figure 5a**). We also observed another, smaller peak at the very end of the GERP score distribution indicative of the efficacy of purifying selection at removing mutations that would have had a negative effect on an individual’s fitness (**Figure 5a**). We however identified a few species for which sites with a high GERP score were much more abundant than more neutrally evolving sites (*Pieris napi* is used as a representative in **Figure 5a**). We also classified GERP scores based on their genomic context, specifically protein-coding and intronic, ignoring other non-coding regions. As expected, we found that higher GERP scores are mostly found in protein-coding sequences, which are known to be highly conserved and under purifying selection, than in intronic regions, where nearly neutrally evolving sites are mostly found (**Figure 5a**).

**Figure 5.**
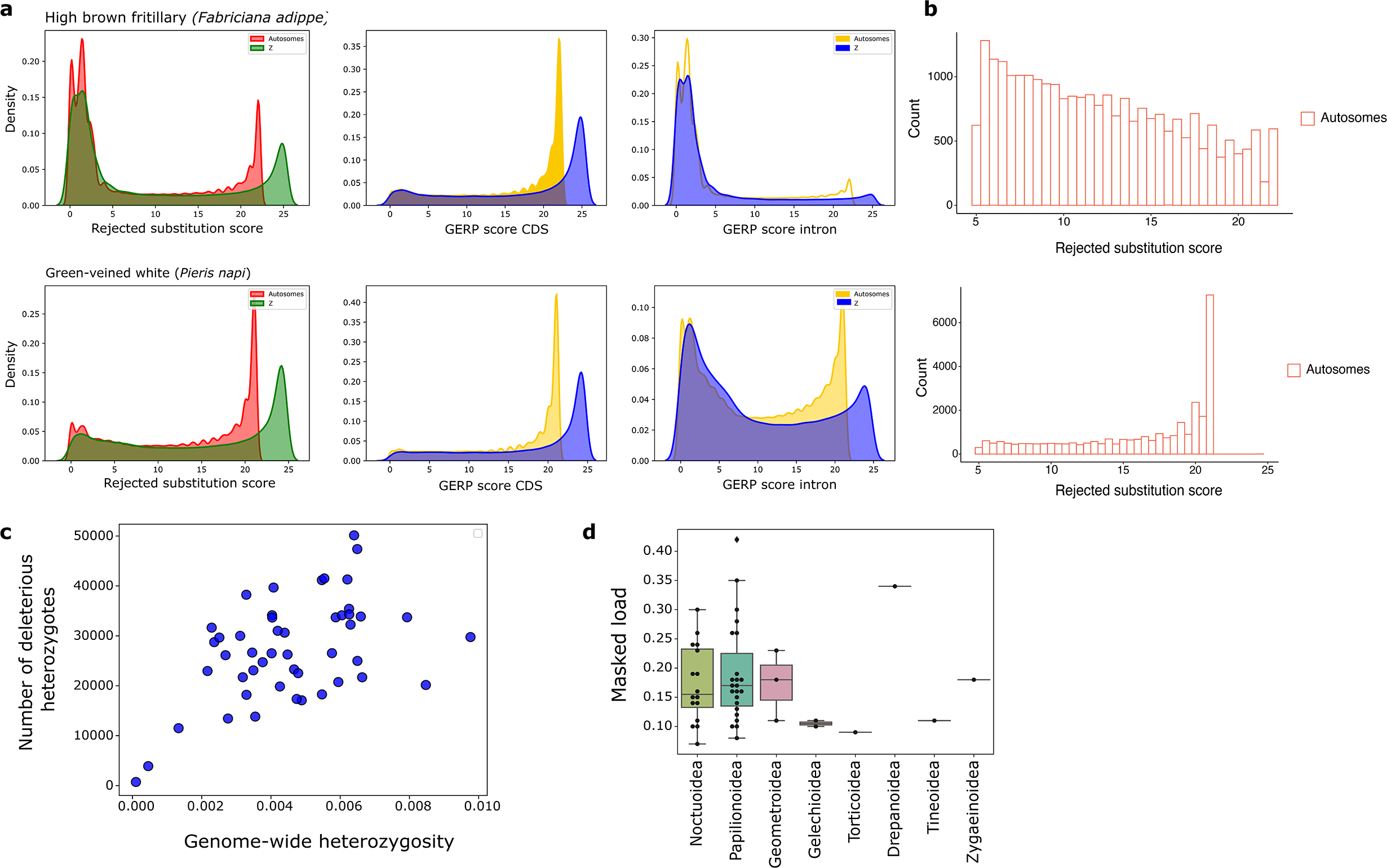
Distribution of rejected substitution scores and genetic load estimates. (a) Distribution of rejected substitution scores along the genome (left), in protein-coding sequences (centre), and in intronic regions (right) for the high brown fritillary (*Fabriciana adippe*) and green-veined white (*Pieris napi*). **(b)** Genome-wide distribution of deleterious heterozygous sites with a rejected substitution score ≥5 in the high brown fritillary (*Fabriciana adippe*) and the green-veined white (*Pieri napi*). **(c)** Genome-wide heterozygosity versus the total number of deleterious heterozygotes (GERP score ≥5) in each species genome. **(c)** Masked load estimated in each individual species from deleterious heterozygous sites (GERP score ≥5). Species are here grouped and coloured based on the superfamily they belong to.

To estimate the masked load, we focused on 48 genomes for which we had both rejected substitution scores and heterozygous calls. The distribution of the GERP score of high-confidence heterozygous sites, reveals that for most species, for example *Fabriciana adippe* in **Figure 5b**, the number of deleterious heterozygous alleles decreased with an increase in the GERP score, consistent with purifying selection being effective at removing highly deleterious variants (GERP score > 20). We found, however, a few exceptions, as shown in **Figure 5b** for *Pieris napi,* where deleterious heterozygous sites were found to be in higher numbers than neutrally evolving sites. We assessed the potential for future inbreeding depression in each species by quantifying the total number of deleterious heterozygotes per genome, which reflects alleles that could contribute to inbreeding depression when made homozygous through inbreeding, assuming a recessive model ^35^. We found that the total number of these deleterious alleles is positively correlated with the genome-wide heterozygosity (Pearson’s correlation: 0.49, p-value: 3.9e-04) (**Figure 5c**). Of all species analysed*, M. brassicae* was found to harbour the lowest number of deleterious heterozygous sites (*n* = 484 sites), whereas *Notodonta dromedarius* had the highest number (*n* = 49,898). Putatively deleterious alleles were used to estimate the individual masked load as the sum of the GERP scores at deleterious sites divided by the number of called heterozygous genotypes per individual. The masked load was, on average, higher in butterflies (superfamily Papilionoidea) (**Figure 5d**) and in our dataset ranged from 0.07 (*Phlogophora meticulosa*) to 0.42 (*Pieris napi*), confirming in the latter the GERP score distribution at heterozygous sites. Because we only had a single diploid per species, we were unable to estimate the realised load from deleterious alleles in a homozygous state. If estimated, the realised load would have provided valuable insights on the actual cost of inbreeding, particularly in highly inbred species like the cabbage moth and the centre-barred sallow ^36^.

## Discussion

Reductions in sequencing costs and rapid improvements in third-generation sequencing technologies mean that reference genome assemblies of non-model organisms are being sequenced at a rapidly increasing pace^37,38,39^. Conservation practitioners have started to appreciate the potential of sequencing data (e.g. ^40,41,42,43^). We focussed on Lepidoptera, which are both widespread and widely recorded, and for which issues of conservation concern are currently highlighted by estimates of precipitous decline in many species in Britain and Europe. We show the benefits of using sequencing data in a conservation context where traditional measures of concern are difficult to acquire. We show that the reference genome of a single diploid individual can be informative when analysed in a comparative framework, especially when combined with a high-quality whole-genome alignment. Our quality assessments show that high-quality, chromosome-level assemblies are key to improving the number of consistently aligned conserved loci in these alignments, and that the depth of the guide tree can influence the quality of the alignment. This finding is echoed in a recent whole-genome alignment of 240 mammalian species ^24^. Importantly, the Cactus toolkit we used can build on our current alignment by adding species, and thus this will form the foundation of future work. We analysed 88 species, but The Darwin Tree of Life Project is already taking the next steps to increase the coverage of British and Irish Lepidoptera by generating assemblies for many additional taxa (see https://portal.darwintreeoflife.org).

Comparing the patterns of variation in the genomes of species of different conservation concerns uncovers the evolutionary history, demography, and molecular adaptations of species ^44^. It supports the prioritisation of species for future targeted population genomics, which is particularly relevant for species that lack long-term monitoring data. By comparing the genomes of 88 Lepidoptera species, we have shown that butterflies and moths display contrasting levels of genetic diversity and inbreeding. This is particularly interesting, if we consider that a large number of butterflies and moths included in this study come from Wytham Wood, an ecosystem exceptionally rich in flora and fauna and one of the most researched pieces of temperate broadleaf woodland in the world. Butterflies and moths form an ecologically diverse and species-rich group, with species living in a wide range of habitats, ranging from forests to treeless regions to wetlands ^45^. Over the past years, trait-based analyses in these insect taxa have provided insights into the ecological drivers of their decline. For instance, the decline of butterflies specialised in dry, open, low-fertility habitats across Europe and North America has been widely demonstrated ^46, 47^ and has been attributed to direct habitat loss, air pollution, and nutrient enrichment through agricultural practices ^48, 49^. Similar patterns have been observed in moths. In the United Kingdom, such decline has been more severe in the south, due to higher rates of habitat loss through agricultural intensification ^50^. Data on British moths also suggest that declines in abundance and biomass since the late 1960s have been more severe in woodlands than in farmland ^51, 52^. This conclusion was recently confirmed by Blumgart and colleagues (2022) who modelled moth abundance, biomass, species richness and diversity from 1968 to 2016 and confirmed their significant decline in broadleaf woodlands ^53^. Our study supports these findings from a genetic perspective, as moths displayed lower levels of heterozygosity and a large number of medium size runs of homozygosity (ROHs), which reflect their long-term decline in effective population size over the past decades ^54, 55^.

Compared to butterflies (superfamily Papilionoidea), most of the moth species included in our study had an unknown risk category assigned to them by the Butterfly Conservation ^55, 56^. As expected, species without long-term monitoring data (*No data*) displayed a heterogenous level of genetic diversity, as much as species classified as of *Least concern*. However, within these two risk categories we found species whose level of heterozygosity and inbreeding would make them a good target for future population genomics studies and conservation initiatives. Examples of these are the cabbage moth, *Mamestra brassicae* (*No data*), and the centre-barred swallow, *Atethmia centrago* (*Least concern*), whose levels of genetic diversity and inbreeding are indicative of a recent and rapid population bottleneck and long-term small population size, respectively. We here suggest that species with low heterozygosity and recent inbreeding (i.e. long ROHs) should be prioritised for the sourcing and sequencing of at least 20 additional individuals across their range to confirm our results and distinguish very local processes (e.g. Wytham Wood) from species-wide ones. Secondarily, we recommend to re-sample species without any risk category assigned to them (i.e. *No data*) to assess the generality of the conclusions we have here drawn from a single diploid genome. The high level of inbreeding observed in the cabbage moth imposes another interesting reflection. This species is primarily known as a pest responsible for severe crop damage of a wide variety of plant species ^57^. The dominance of long ROHs could also reflect the consequences of strong selection by horticultural practice, such as extensive pesticide use. By the measures outlined above, this species could be considered as being of conservation concern, but equally its current ubiquity and low genetic diversity could be driven by colonisation of a new, human-constructed niche. Because our sample shows signs of intense recent inbreeding, which would most likely be local, it would be important to obtain further data from other samples across the range. For example, sampling of specimens not associated with intensive agriculture would identify whether there is a reservoir of diversity in non-agroecosystem habitats, and thus direct credible conservation interventions.

The dominance of short ROHs in most Papilionoidea (butterfly) genomes indicates a very different evolutionary history. These homozygous tracts are often a consequence of a species with historically limited local population size and/or low dispersal. Even though these short segments cover a small fraction of each genome, their abundance was surprising given the marked declines in the abundance and distribution of many species of butterflies. Over half of British butterflies have been assessed by the Butterfly Conservation as either threatened or near threatened with extinction regionally ^28^. This dire situation has once more been highlighted in the latest report of the Butterfly Conservation on the status of UK’s butterflies, according to which the distribution of 58 native species has fallen by 42%, while that of species living in specific habitats has declined by 68% ^54^. Due to the nature of their conservation status, sampling occurred outside the United Kingdom for species classified as *Regionally extinct*, *Endangered*, and (in most cases) *Vulnerable*. Although the genetic diversity and inbreeding analyses on these species does not reflect the genomic diversity within the United Kingdom, they nonetheless provide useful information for future conservation programmes of British butterflies. For instance, individuals of the same species found in continental Europe could potentially be translocated to the United Kingdom to boost local populations. Moreover, as exemplified by the case of the endangered Apollo butterfly (*Parnassius apollo*), translocations could initially take place in many sites selected based on biotic and abiotic criteria derived from existing populations of the species, to focus, on a later stage, on a smaller set of promising sites ^58^. Genetic rescue via translocations is a common species recovery practice that has been explored in many vertebrate and invertebrate species for a long time. However, as demonstrated in recent studies of mammals, a successful recovery initiative requires translocations to be carried out repeatedly over a long period of time to prevent the genomic consequences of inbreeding to quickly erase any potential gains ^59^. At the same, habitat restoration and management should be part of the translocation efforts for these to be successful, as also suggested by the failed reintroduction of the swallowtail (*Papilio machaon*) to the Wicken Fen Nature Reserve, whose population, after an initial expansion, crashed as a result of changes in the habitat and status of the butterfly’s food plant, *Peucedanum palustre,* in the nature reserve ^60^. Although improvements must be made in the context of invertebrate species on, for example, the objectives of post-release monitoring, the use of genetic screening, and habitat restoration ^61^, genetic rescue has all the premises to improve the outcomes of conservation programmes also in Lepidoptera.

Methods that aim to estimate changes in effective population size (*N_e_*) over time can deepen our understanding of species dynamics, by uncovering past population expansions and contractions. The PSMC framework ^29^ is one of the most popular methods, as confirmed by its application to a great variety of model and non-model organisms (e.g. ^62,63,64,65^). In contrast to large, long-lived taxa such as mammals and birds, we observed that in Lepidoptera PSMC struggles to reliably estimate changes in *N_e_* when the recombination rate is much larger than the mutation rate ^31^. We suggest that, rather than past *N_e_*, the PSMC curves shown in this study may reflect the landscape of variation in heterozygosity due to linked selection. Our results suggest the limited applicability of PSMC to insect genomes and highlight the need for further research on this limitation for their use on a range of applications, including the parameterization of forward-in-time evolutionary genomic models (e.g. ^15^). While we do not question the ability of the PSMC framework to reconstruct changes in *N_e_* over time in appropriate circumstances, we highlight that PSMC outputs do no always reflect past demographic changes and that the curves that are obtained should always be carefully evaluated, particularly in species with small genomes, large population sizes, and low mutation rates.

We have shown that genomic data can also provide valuable information on the total number of harmful mutations that contribute to the genetic load. Understanding the genetic load is an important step in conservation genomic studies, because it poses a considerable threat to the health and viability of endangered populations, both now and in the future ^15, 36^. Although we did not have fitness information, a common limitation in studies of non-model organisms, we were nonetheless able to estimate *in-silico* one of the two components of the genetic load - the masked load - from sequencing data alone. As extensively described in Bertorelle et al. (2022), the masked load (also called inbreeding load) consists of deleterious mutations whose effects on fitness are hidden because heterozygous ^36^. This implies that in large populations a high masked load does not necessarily represent an immediate cost for fitness, as long as the population remains large and genetic drift is negligible. Deleterious mutations represent a cost for a population when a deleterious allele is expressed, which occurs when sites are homozygous for recessive deleterious alleles (the so-called realised load). Different processes can contribute to the conversion of the masked into realised load, one of which is a population bottleneck. Predicting both types of load could provide clues on the role of genetic drift and purging on the deleterious variation landscape in each species, separately. Therefore, we highly encourage future population genomic studies on the Lepidoptera species we abovementioned to estimate both load components to confirm some of the patterns reported in this study.

## Conclusions

Lepidoptera are important umbrella species for biodiversity conservation and key indicators of environmental change and wider invertebrate declines. Therefore, it is of paramount importance to maintain up-to-date assessments of the threats and extinction risks butterfly and moth species are facing. As we have seen here, there are many species with low levels of genetic diversity and high levels of inbreeding. Sequencing data alone are not sufficient to tackle the complex issue of biodiversity loss. However, they provide a start, allowing attention to be focused on more in-depth studies. These can involve further population genomic sequencing of additional samples from the focal site and/or from additional sites, as well as more traditional conservation ecology approaches. Genomics-informed conservation programmes are becoming common for vertebrates. We believe that the comparative genomics analyses made available through this study provide a route towards their application across the great diversity of invertebrate species.

## Authors contributions

Conceptualization, C.B.; Methodology, C.B., C.J.W., S.L., T.C., T.A.P.G.; Investigation, C.B.; Resources, DToL Consortium, M.B., R.D., F.J.M., D.T., L.H.; Data curation, C.J.W., T.A.P.G., F.J.M., D.T., L.H.; Writing – Original Draft, C.B.; Writing – Review & Editing, C.B., C.J.W., S.L., T.C., R.D.; Funding Acquisition, M.B., R.D.; Supervision, R.D.

## Data and code availability

All data generated in this study have been deposited in Zenodo under 10.5281/zenodo.7755299.

Codes used to build the guide tree can be found at https://github.com/lstevens17/busco2phylo-nf and https://github.com/nylander/catfasta2phyml. Other codes used in this study can be found https://github.com/cbortoluzzi/LepCactusAlignment2023.

## Method details

### 1. Sample selection

The dataset included reference genome assemblies for a total of 88 Lepidoptera species from 12 superfamilies, plus one Trichoptera outgroup (**Figure 1)**. Of these, 76 were generated by the Darwin Tree of Life (DToL) Project ^23^, and the remaining 13 were obtained from the International Nucleotide Sequence Database Collaboration (INSDC) (**Table S1**). Sequencing and assembly details are described, for each species, in its respective primary assembly publication (**Table S1**); where references are not given for DToL species the process was the same as for ^66^. The 76 newly sequenced genomes sequenced by DToL are chromosome-level assemblies that comply with the assembly quality metrics proposed by the Vertebrate Genomes Project (VGP), which are a contig N50 > 1 Mb and a scaffold N50 > 10 Mb ^43^. For *Lysandra coridon* and *Pheosia gnoma,* scaffold N50 was < 10 Mb, because the chromosomal N50 in these species is < 10 Mb (**Table 1**). All 12 INSDC genomes met the VGP assembly quality metrics, except for *Cnaphalocrocis medinalis, Ephestia elutella, Manduca sexta, Spodoptera litura,* and *Zerene cesonia* for which contig N50 was < 1 Mb, and *Danaus plexippus, Spodoptera litura,* and *Zerene cesonia* for which scaffold N50 was < 10 Mb (**Figure 1**). All genomes generated by the DToL Project are publicly available (see https://portal.darwintreeoflife.org) and can be accessed through the European Nucleotide Archive (ENA) and other INSDC databases. Species whose reference genome assembly was generated by DToL were sampled in the United Kingdom, except for 15 that were sampled in Spain, Romania, or Hungary (**Table S1**). Finally, we also assigned to each species the latest Great Britain conservation category, as recently revised using methodology based on that used for the IUCN Red List by the Butterfly Conservation for moths^55^ and butterflies ^56^ (**Table S1)**.

Species with transcriptomic data publicly available were annotated by Ensembl with the Genebuild process using both protein homology information and RNA-seq data. For species that did not have transcriptomic data, the annotation was performed using BRAKER2 ^67^. Protein data consisted of OrthoDB (v11) data ^68^ for Lepidoptera combined with all lepidopteran proteins with protein existence levels 1 or 2 from UniProt ^69^, where level 1 or 2 represent evidence from either proteomic or transcriptomic data. Gene annotations are publicly available and were downloaded from the Ensembl Rapid Release platform (last access: June 2022) (**Table S1**) ^70, 71^.

### 2. Reference-free, whole genome multiple sequence alignment

We ran Cactus (v2.0.5) ^72^ on the Google Cloud platform using 96 CPUs and 8 GPUs. The alignment took approximately 4.5 days to complete. We chose Cactus over other widely used sequence aligners, because it does not constrain the multiple sequence alignment to only ‘single copy’ regions present in a single reference genome, thus limiting ‘reference bias’ while capturing multiple-orthology relationships.

#### 2.1 Assemblies preparation

Before running Cactus, we performed a preliminary repeat-masking step on all Lepidoptera genomes and the outgroup Trichoptera genome (see section below on Guide tree). The identification and masking of repeats was carried out using a combination of tools, including Red (v2.0) ^73^, DUST ^74^, and TRF (v4.09.1) ^75^.

#### 2.2 Guide tree

Cactus requires a phylogenetic tree with branch lengths as a guide tree. We built this guide tree from loci identified though benchmarking universal single copy orthologs (BUSCO) ^76^. The phylogenetic tree included 88 species of Lepidoptera and the outgroup *Hydropsyche tenuis* (Order: Trichoptera; Family: Hydropsychidae) (**Figure 1**). The species phylogeny was estimated using a supermatrix-based approach as implemented in the pipeline https://github.com/lstevens17/busco2phylo-nf. Briefly, BUSCO gene sets were filtered to retain the 1,542 genes that are single copy and present in all 89 species. Then, a multiple sequence alignment was constructed for each of these using MAFFT (v7.475) ^77^. Alignments were then trimmed using trimAl (v1.4) ^78^ with the options -gt 0.8 -st 0.001 -resoverlap 0.75 -seqoverlap 80. Next, the trimmed alignments were concatenated to make a supermatrix using the catfasta2phyml script available at https://github.com/nylander/catfasta2phyml. Finally, this supermatrix was used in IQ-TREE (v2.03) ^79^ to infer the species tree under the LG substitution model with gamma distributed rate variation among sites and 1000 ultrafast bootstrap replicates ^80, 81^.

### 3. Quality of the whole-genome multiple sequence alignment

We developed a set of alignment quality metrics (**Figure S1**) to evaluate the quality of our reference-free, whole-genome alignment.

#### 3.1 Calculate coverage by sampling bases

Coverage was calculated for each assembly on 100 intervals of 1 Mb, randomly sampled along the genome (**Figure S1a**). Intervals were generated using the *random* utility in Bedtools (v2.29.1) ^82^ with the options *-l 100000* and *-n 100*. For each of the 100 intervals, we obtained a whole-genome multiple sequence alignment in multiple alignment format (MAF) using the hal2maf utility (v2.1) in Cactus, retaining only orthologs to the species used each time as reference *(--*onlyOrthologs) and excluding ancestral sequences (-*-*noAncestors). Coverage was calculated using the *coverage* option in maf_stream ^25^ and is here expressed as the proportion of sites in species A (i.e. query) that aligns to *n* or more sites in species B (i.e. reference/target) (**Figure S1a**).

#### 3.2 Consistency of benchmarking universal single copy orthologs (BUSCO) and orthogroups

Next, we checked whether Cactus was able to correctly align (1) 2,451 benchmarking universal single copy orthologs (BUSCO) loci shared by all 88 Lepidoptera species and (2) 2,501 single copy orthogroups shared by the 49 Lepidoptera species for which amino acid sequences of protein-coding genes were available (Cunningham et al. 2022). The BUSCO set was obtained by running BUSCO (v5.0.0) on each reference genome assembly with default settings and the *lepidoptera_odb10* lineage (**Table S1**). Orthogroups were identified in OrthoFinder (v2.5.4) ^27^ using as input the longest transcript variant per gene.

We implemented a pairwise approach, where the most basal species in the dataset, *Tinea trinotella*, was used as reference, for a total of 87 (BUSCO) and 48 (OrthoFinder) pairwise comparisons. For each pairwise comparison, and each BUSCO/orthogroup, we obtained a whole-genome multiple sequence alignment in multiple alignment format (MAF), retaining only orthologs to *T. trinotella* and excluding ancestral sequences. We allowed for some flexibility, by taking 100 bp up- and downstream the genomic coordinates of each BUSCO/orthogroup in the species used each time as target. The BUSCO/orthogroup was classified as consistent if the genomic coordinates reported in the Cactus alignment matched the ones reported by BUSCO/OrthoFinder, otherwise it was classified as inconsistent (**Figure S1b**).

#### 3.3 Alignment depth of coding sequences

We calculated the alignment depth on unique, non-overlapping coding sequences. Alignment depth was expressed as the number of other unique genomes each base aligns to (excluding ancestral genomes) and it was calculated on the 49 Lepidopteran species for which a gene annotation was available (**Figure S1c**).

### 4. Heterozygosity

#### 4.1 Selection of assemblies for the heterozygosity analysis

Heterozygosity was estimated for all genomes for which PacBio HiFi data were available (*n* = 74). This excluded 13 INSDC genomes and one DToL genome.

#### 4.2 Variant calling and selection of high-confidence sites

We used the DeepVariant pipeline (v1.1.0) ^83^ on each of the 74 assemblies to call germline mutations and to define callable regions of the genome. DeepVariant is a deep-learning based variant caller that takes as input aligned reads in BAM or CRAM format and the assembled reference genome in FASTA format. Reads are used to produce pileup image tensors, which are subsequently classified using a convolutional neural network. Variants are reported in a standard VCF file and are classified into homozygous reference, heterozygous, and homozygous alternative based on their probabilities. As the reads and the reference genome are derived from the same individual, homozygous mutations reflect assembly or mapping errors and were thus excluded from our downstream analyses.

After assessing the distribution of the PHRED-quality score (QUAL), read depth (DP), and genotype quality (GQ), we defined a high-confidence variant any heterozygous site with a PHRED-quality score >15, a minimum read depth of 6x, a maximum read depth equal to 2 times the average genome coverage, and a genotype quality >20. The average genome coverage was calculated directly on the BAM file using the *depth* command in Samtools (v1.11) ^84^. On average, 2 070 906 heterozygous sites (or 424 SNPs/100kb) at an average distance of 437 bp were retained for further downstream analyses (**Table S2**).

#### 4.3 Inferring genome-wide heterozygosity

We estimated individual heterozygosity on a genome-wide scale by dividing the genome of each individual (excluding its sex chromosomes) into non-overlapping 10-kb windows. We corrected the total number of high-confidence heterozygous sites called within each window for the total number of sites not called because of low coverage, following Bosse et al. 2012 and Bortoluzzi et al. 2020. Insufficiently covered bins, which were here defined as 10 kb windows with less than 6 000 callable sites, were excluded from the genome-wide heterozygosity. Genome-wide heterozygosity was thus estimated on >90% of each genome, except for *Pieris napi*, for which 20% of the genome was excluded from subsequent analyses. For this analysis we discarded the W chromosome.

### 5. Runs of homozygosity

Runs of homozygosity (ROH) were identified as genomic regions showing lower heterozygosity than expected based on the genome-wide average. To identify ROHs, we considered 10 consecutive 10 kb bins . In each 100 kb, we calculated the average heterozygosity from only 10 kb bins where the number of well covered sites was ≥6,000; we then compared this bin heterozygosity to the average genome-wide heterozygosity and retained only the 10 consecutive bins where the heterozygosity level was below 25% of the average genome-wide heterozygosity. To minimise possible local assembly errors or alignment errors, we performed an additional filtering step, by allowing, within each candidate homozygous stretch, a 10 kb bin with an heterozygosity level no higher than twice the average genome-wide heterozygosity, only if, by including this peak, the heterozygosity within the final ROH did not exceed, once again, 25% of the average genome-wide heterozygosity ^85, 86^. We classified ROHs into three size classes, each of them corresponding to a specific demographic event, including past relatedness (small: ≤0.1 Mb), background relatedness (medium: 0.1 -1 Mb), and recent inbreeding or relatedness (long: ≥1.0 Mb).

### 6. Demographic history

To better understand the evolutionary processes acting on Lepidopteran genomes, we applied the pairwise sequentially Markovian coalescent (PSMC) ^29, 87^ to each of the 74 species for which we had PacBio HiFi data. Using just a diploid sequence, PSMC seeks to reconstruct the changes in the effective population size (*N_e_*) over time by estimating the inverse of the inferred coalescence rate. By estimating the time to the most recent common ancestor (TMRCA) at every position across the genome, PSMC infers a model of population size changes over time that best explain this distribution ^29^. For our PSMC analysis, we retained only SNPs (indels excluded) that had a read depth between 10 and 100, a genotype quality >30, and a PHRED-scaled genotype likelihood >25. SNPs that did not pass these criteria were masked and treated as missing data. To interpret the inference of the PSMC analysis, we used a mutation rate of 2.9e-09 per generation per base pair, as calculated on trios of *Heliconius melpomene* ^88^. The PSMC analysis was performed using 20 iterations and 32-time segments with the first 28 free and the final four fixed to the same value. Results were scaled to real time using a generation time of 3 months and the mutation rate of Keightley et al. 2015.

### 7. Genomic Evolutionary Rate Profiling

We used Genomic Evolutionary Rate Profiling (GERP++) ^34^ to calculate the number of rejected substitutions for each nucleotide in the multiple sequence alignment. Given a multiple sequence alignment and a phylogenetic tree with branch lengths representing the neutral rate between the species within the alignment, GERP++ quantifies constraint intensity at each individual position in terms of rejected substitutions, which is the difference between the neutral rate and the estimated evolutionary rate at the position ^34^. The GERP score is thus an excellent measure of the strength of selection acting on derived mutations segregating within a species ^89^.

#### 7.1 Neutral model

We estimated a neutral model on the basis of 4-fold degenerate sites obtained from unique, non-overlapping protein-coding sequences of transcripts without in-frame stop codons. As our approach requires a gene annotation file, the GERP score analysis was run on the 48 lepidopteran species annotated by Ensembl (**Table S1**). The 4-fold degenerate sites extracted using msa_view ^90^ were used to estimate a neutral (or nonconserved) phylogenetic model in phyloFit ^90^, using the topology of the guide tree and the REV substitution model. As sex chromosomes are known to evolve at a different rate to autosomes, we obtained two neutral models, one for the autosomes and one for the Z chromosome. The W chromosome was excluded from this analysis, because of its low protein-coding content and presence only in females (and thus a subset of our genome sequences).

#### 7.2 GERP++

We ran the *gerpcol* option in GERP++ ^34^ to compute the neutral rate and rejected substitution score (or GERP score) at every position in a multiple sequence alignment block with at least 6 species and no more than 3 gaps at each position. To avoid biases towards the focal genome, we used the *-j* option to project out the reference genome. Sites in the multiple sequence alignment with a rejected substitution score ≥0 were used to plot the genome-wide distribution of the GERP score in the autosomes and Z chromosome, respectively. A similar distribution was obtained from the same sites, after classifying them into protein-coding and non-coding (i.e. intronic) regions. The rationale for discarding sites with a rejected substitution score below 0 is that negative sites represent sites that are expected to evolve at a neutral rate ^34^.

### 8. Genetic load estimates

#### 8.1 Identification of deleterious alleles

The range of GERP scores obtained from a particular alignment depends on the depth of the corresponding phylogenetic tree, and thresholds for judging whether mutations should be considered deleterious or not will depend on the phylogenetic relationships among the species in the multiple sequence alignment. As a guide for setting a threshold, we compared the distribution of the GERP scores in each species, separately, and found a score of 5 to represent a compromise between excluding as many as possible neutral sites, while including as many as possible potentially deleterious sites. Therefore, in this study we classified deleterious any high-confidence heterozygous site with a GERP score ≥5.

#### 8.2 Genetic load component: the masked load

The genetic load is the occurrence of deleterious alleles in the population and can be divided into realised load (or expressed load) and masked load ^36^. In this study, we could only estimate the masked load, which is made of deleterious alleles, whose effect on an individual’s fitness is hidden by being heterozygous. Given both the GERP scores and the genotype call, we estimated the masked load as:

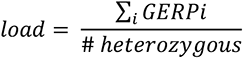

where *GERPi* is the GERP score of a deleterious allele in heterozygous state at genomic position *i*. The sum of GERP scores was divided by the number of called heterozygous genotypes per individual to account for differences in callability between individuals.

