## Supplemental Figure 1 for "Lepidoptera genomics based on 88 chromosomal reference sequences informs population genetic parameters for conservation"

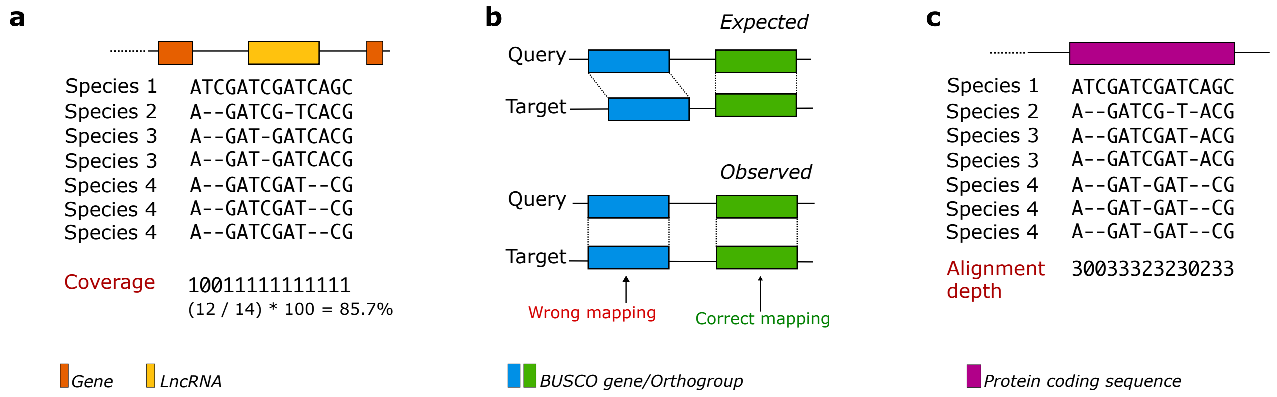


Figure S1: **Alignment quality metrics**. Set of alignment quality metrics developed to assess the quality of the cactus alignment. **a.** Coverage estimated in each species genome by randomly sample 100 regions along the genome, each 1 Mb long. Coverage was estimated independently of the genome annotation. **b.** Mapping consistency of single copy BUSCO genes and single copy orthogroups identified by OrthoFinder. The expected mapping coordinates are the ones identified in either BUSCO or OrthoFinder, whereas the observed mapping coordinates are the ones identified in the Cactus alignment. A gene/orthogroup was classified as correctly mapped (or consistent) if the expected and observed mapping coordinates matched. **c.** Alignment depth at unique, non-overlapping protein coding sequences here expressed as the number of unique genomes that map to each site in the reference/target species.
