## Supplemental Figure 2 for "Lepidoptera genomics based on 88 chromosomal reference sequences informs population genetic parameters for conservation"

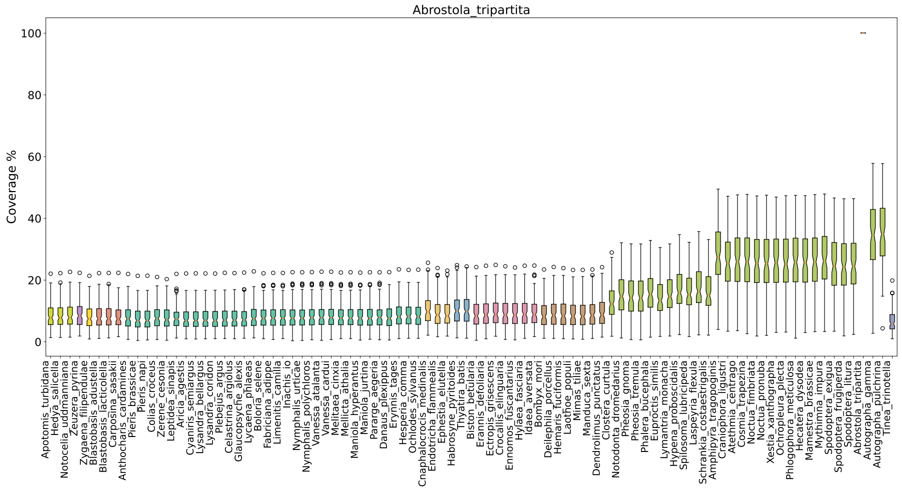
Abrostola_tripartita
Coverage %
0
20
40
60
80
100

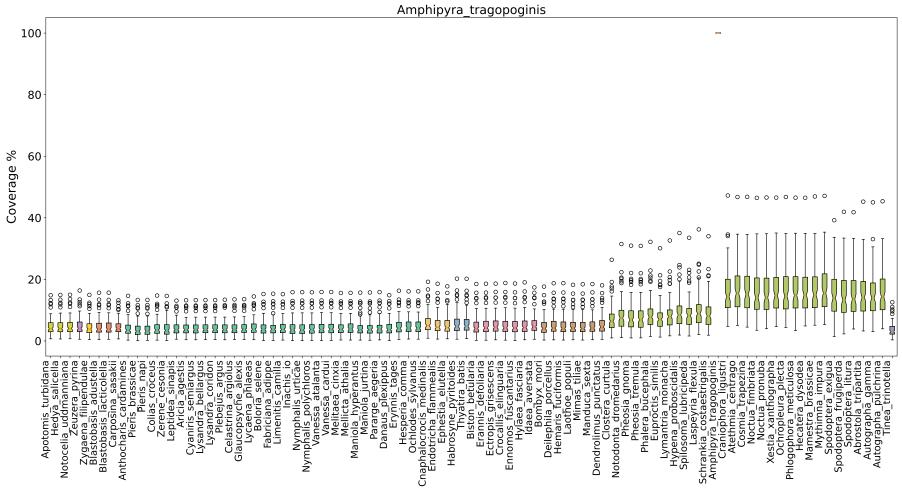
Amphipyra_tragopoginis
Coverage %
0
20
40
60
80
100

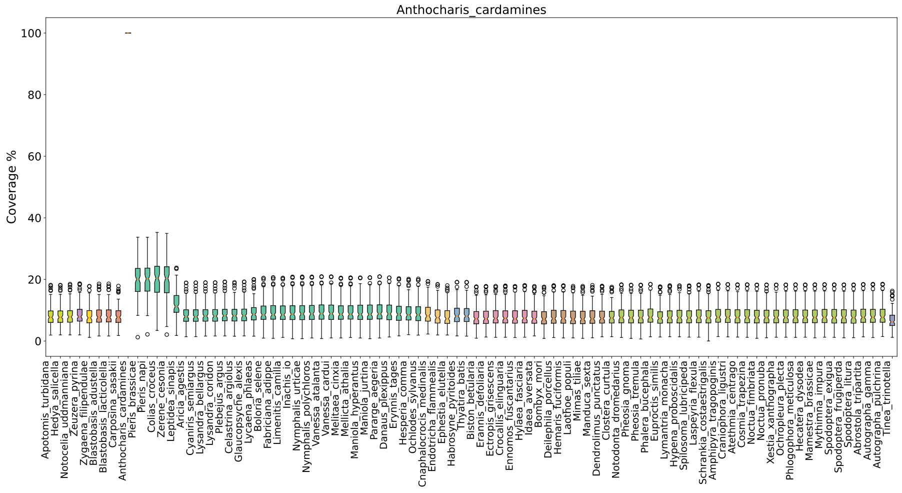
Anthocharis_cardamines
Coverage %
0
20
40
60
80
100

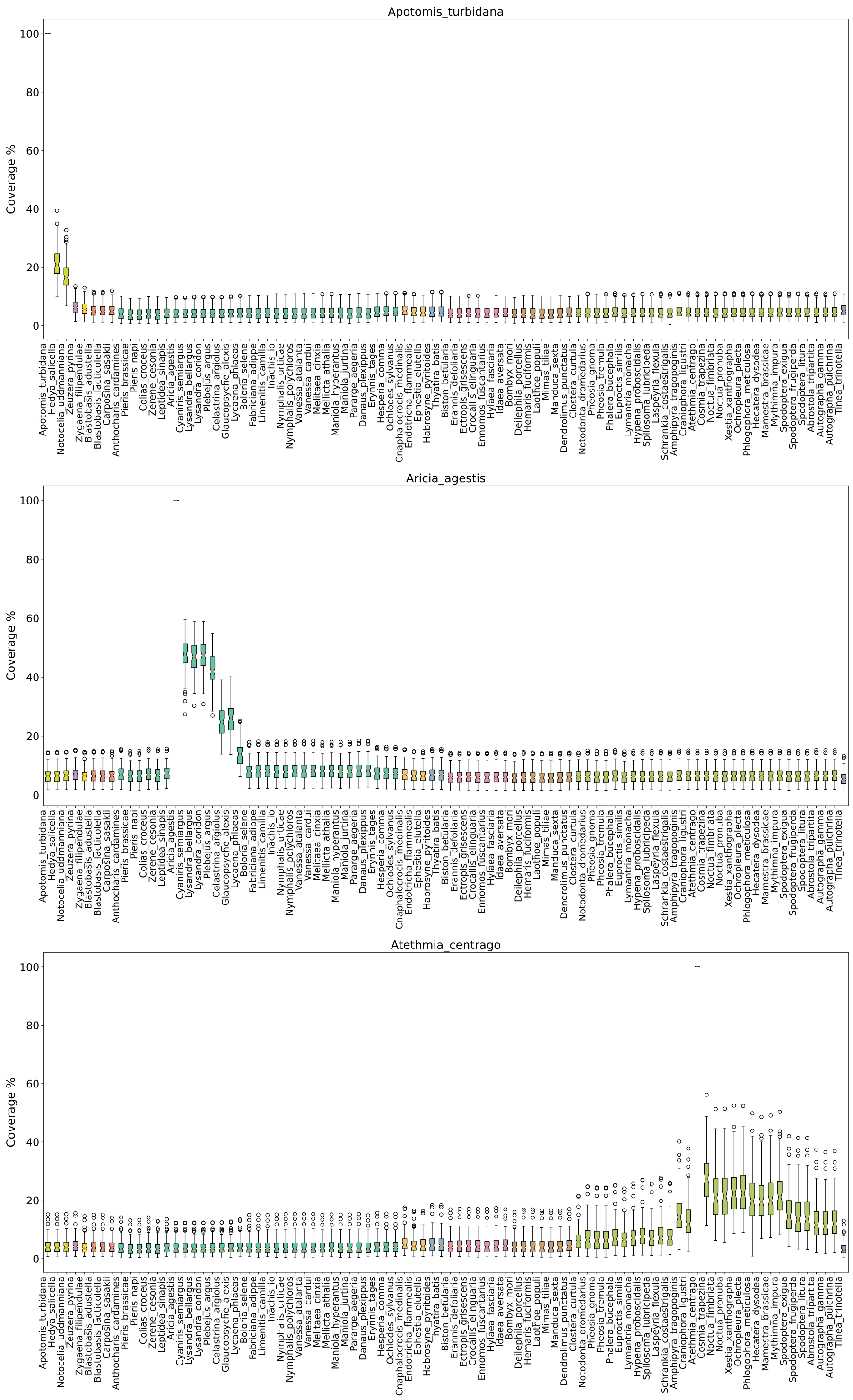
Apotomis_turbidana
Coverage %
Aricia_agestis
Coverage %
Atethmia_centrago
Coverage %

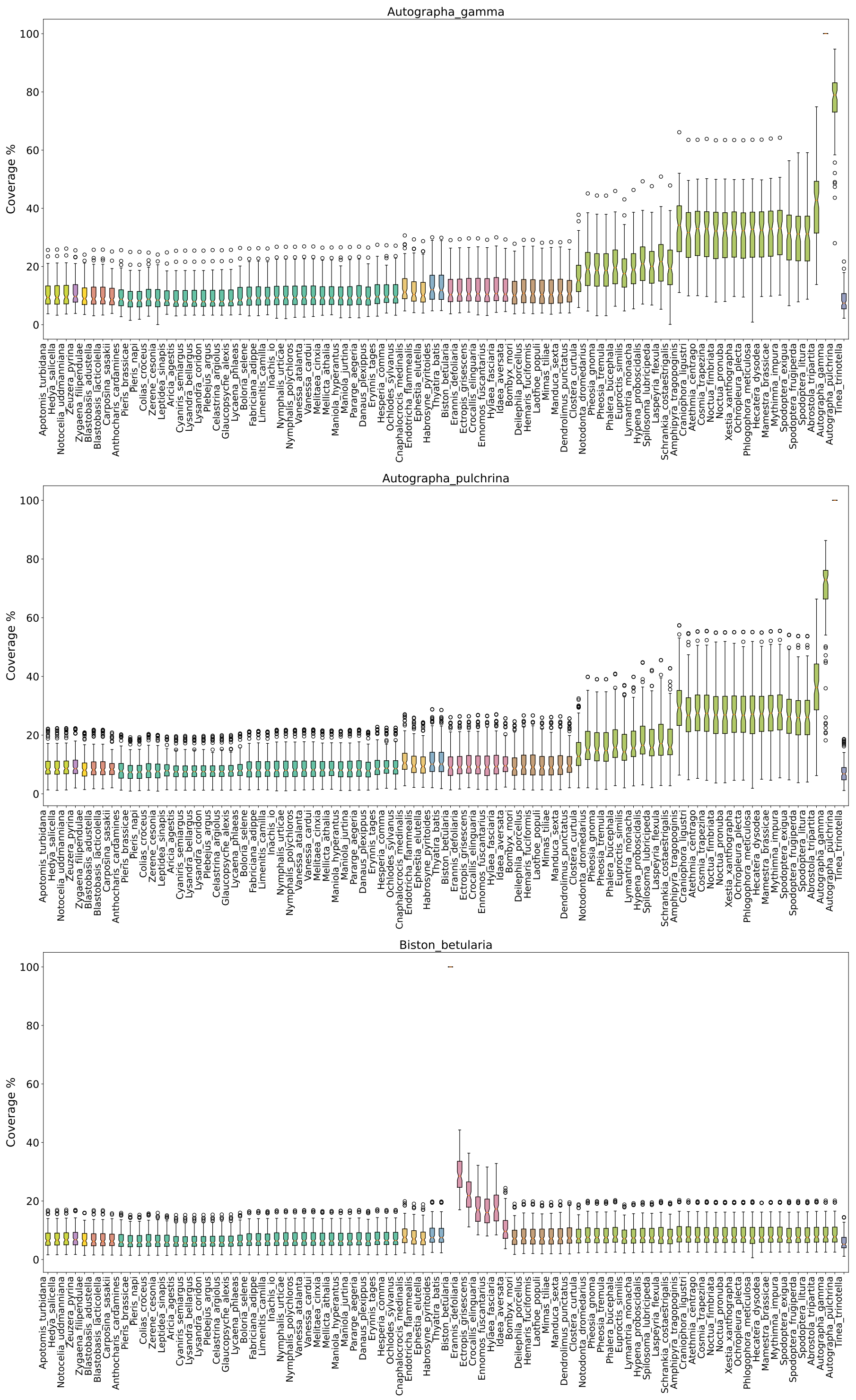
Autographa_gamma
Autographa_pulchrina
Biston_betularia
Coverage %
0
20
40
60
80
100

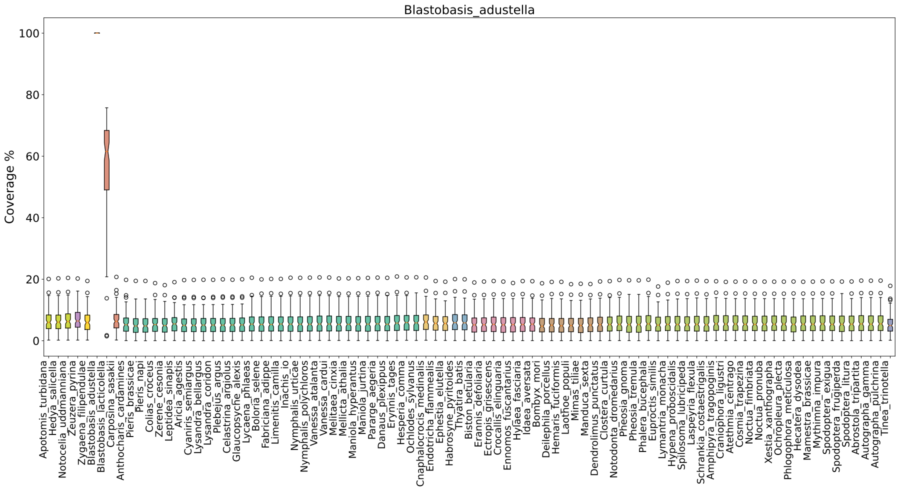
Blastobasis_adustella
Coverage %

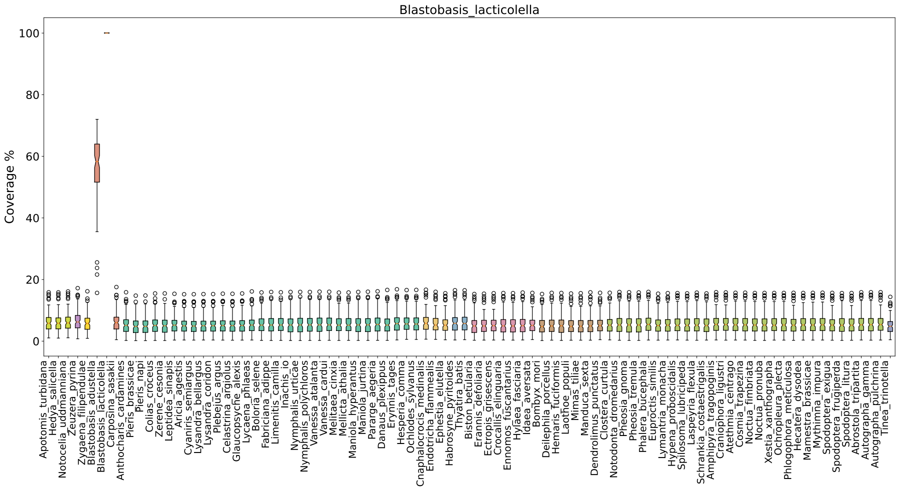
Blastobasis_lacticolella
Coverage %

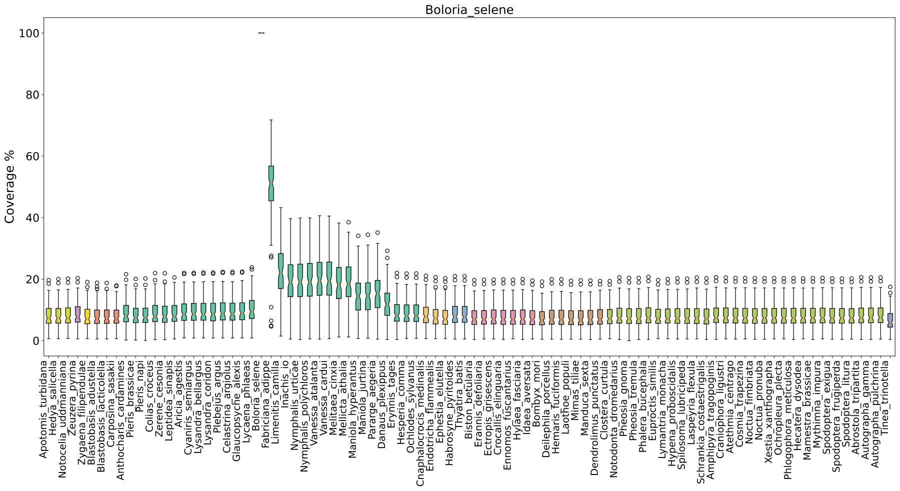
Boloria_selene
Coverage %

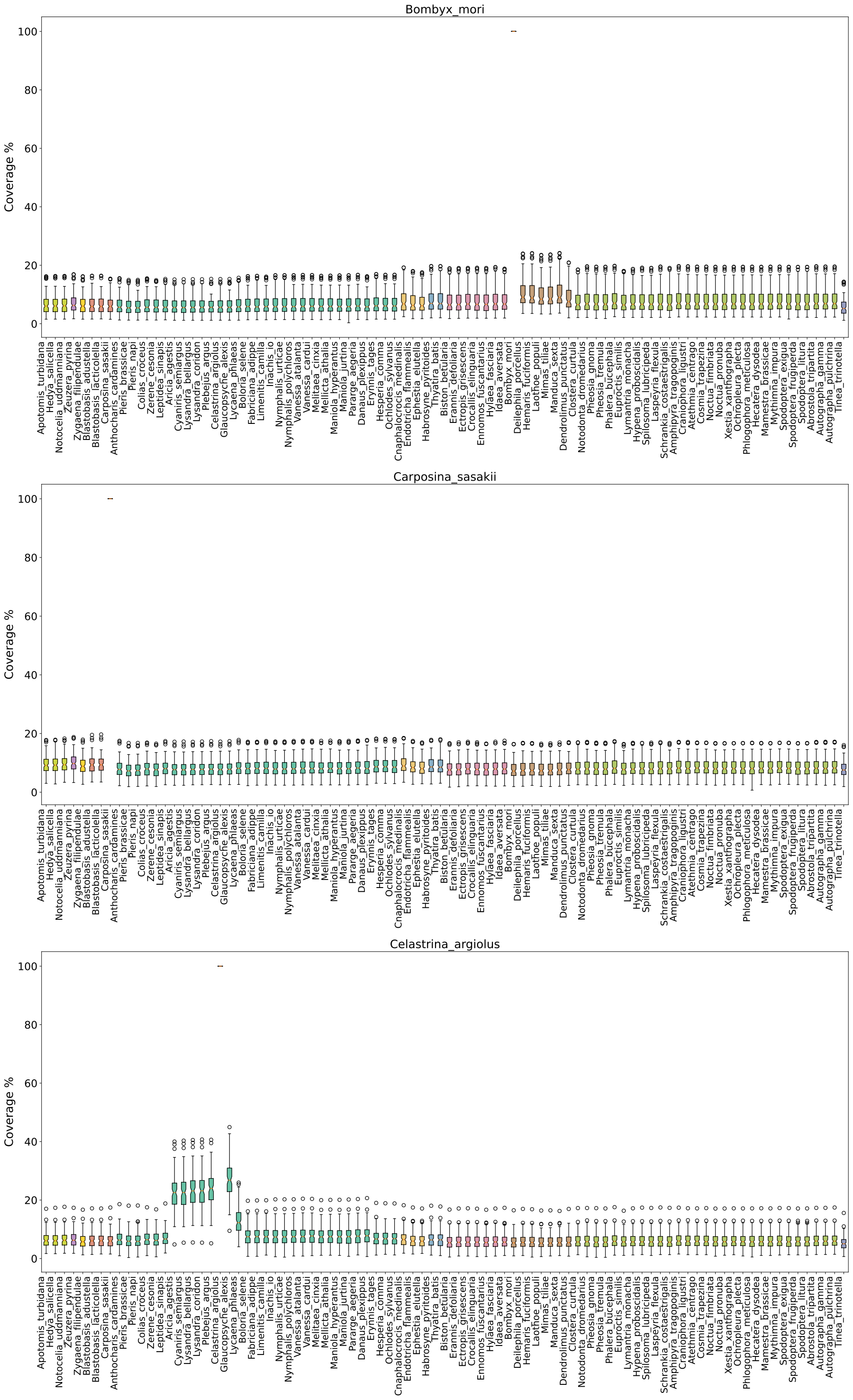
Bombyx_mori
Carposina_sasakii
Celastrina_argiolus
Coverage %
0
20
40
60
80
100

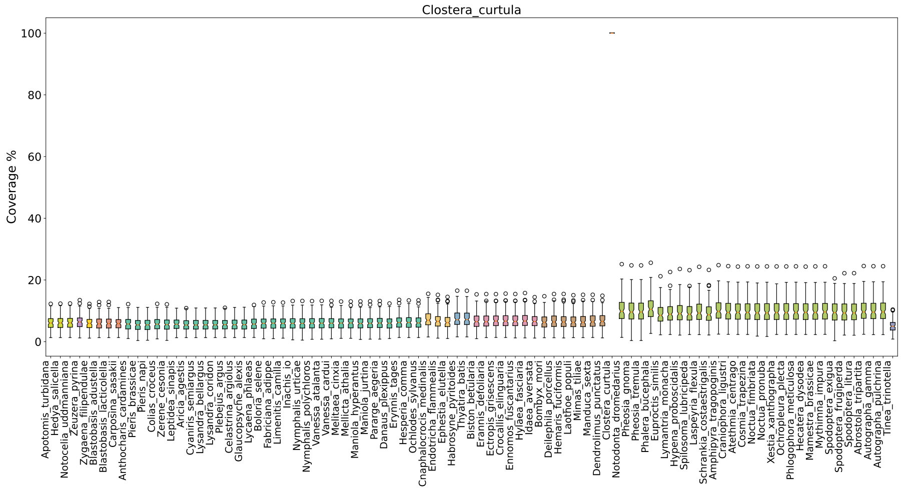
Clostera_curtula
Coverage %
100
80
60
40
20
0

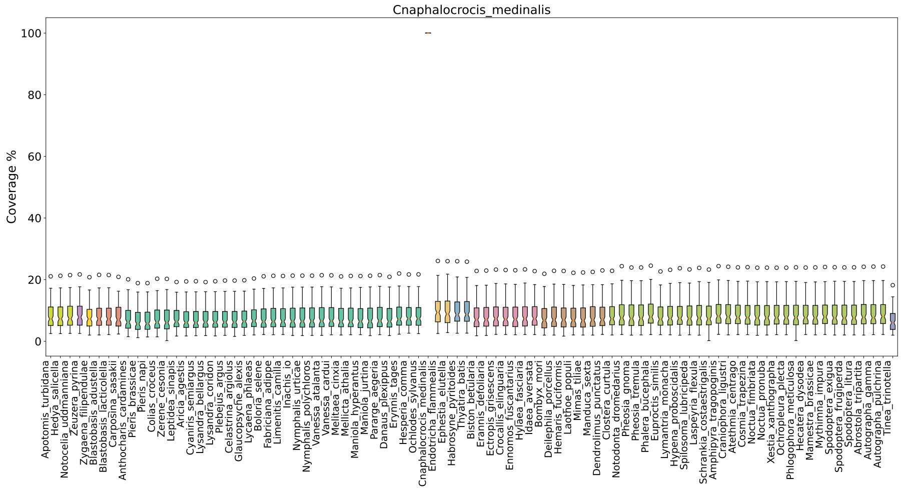
Cnaphalocrocis_medinalis
Coverage %
100
80
60
40
20
0

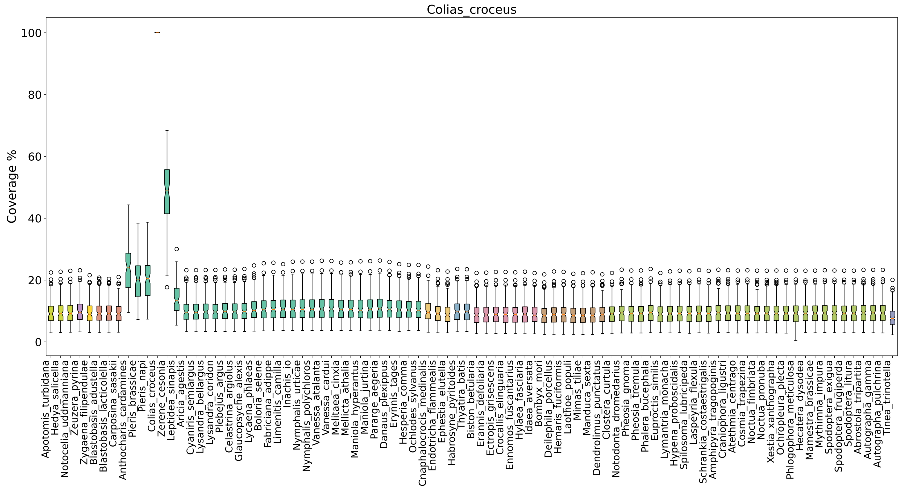
Colias_croceus
Coverage %
100
80
60
40
20
0

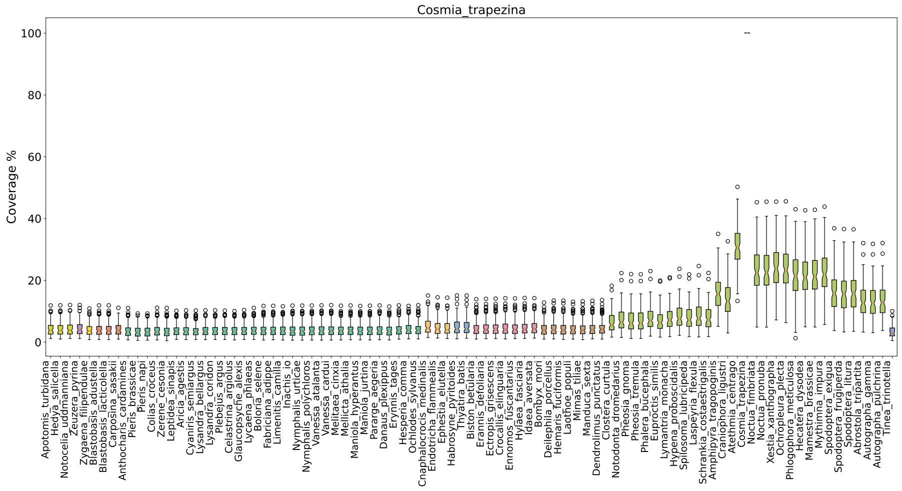
Cosmia_trapezina
Coverage %

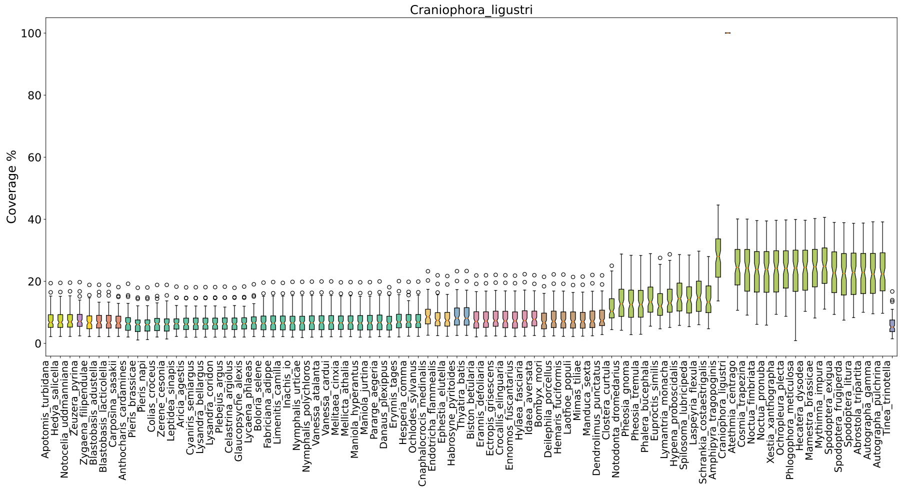
Craniophora_ligustri
Coverage %

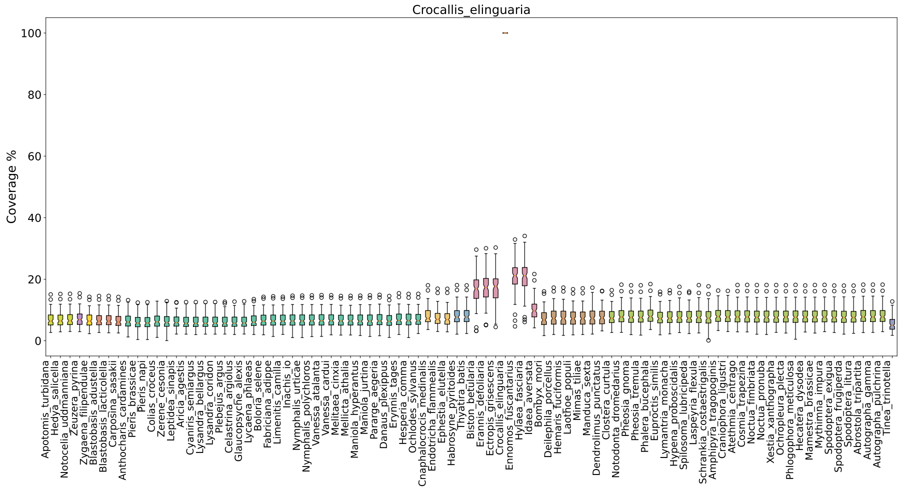
Crocallis_elinguaria
Coverage %

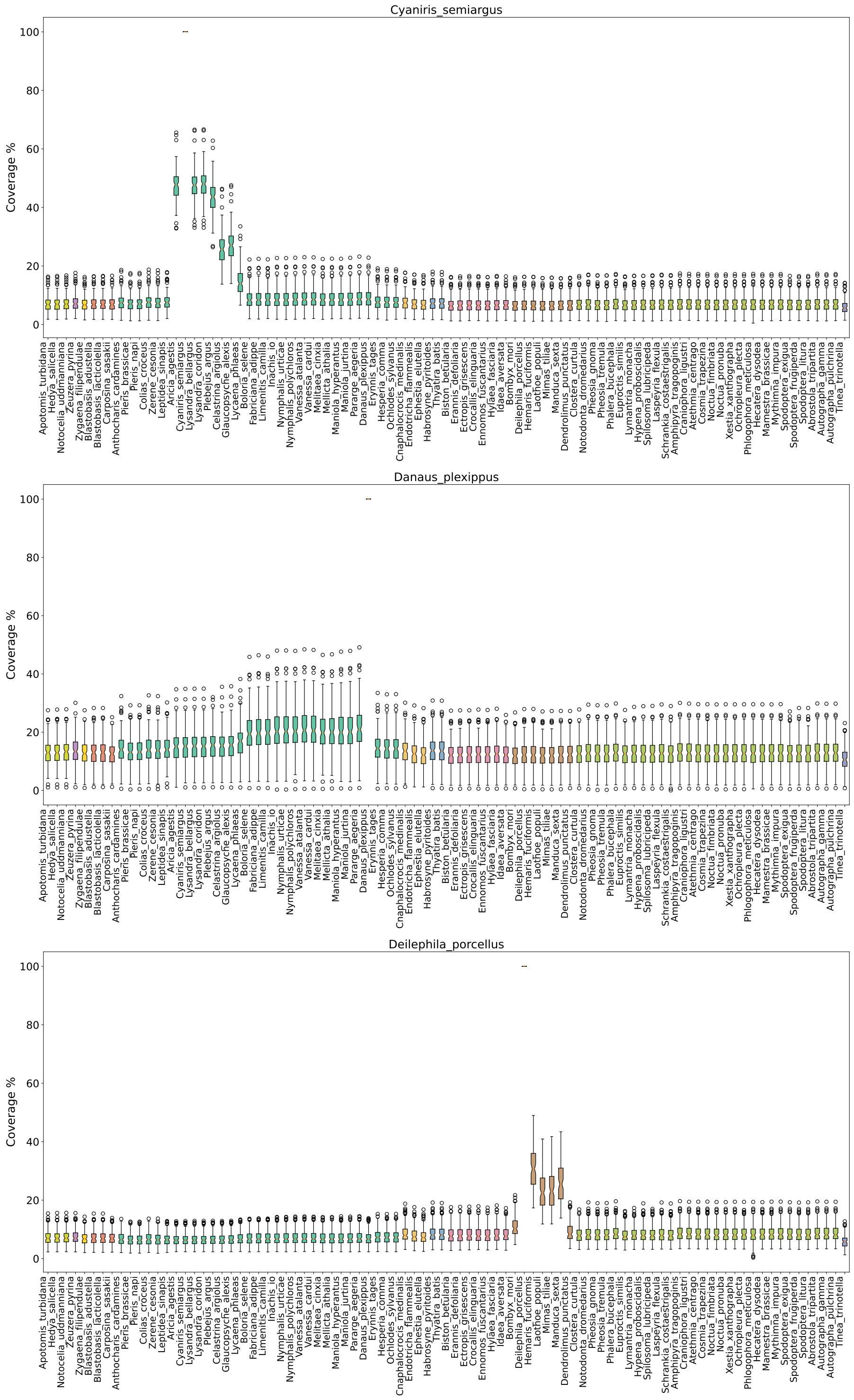
Cyaniris_semiargus
Coverage %
Danaus_plexippus
Coverage %
Deilephila_porcellus
Coverage %

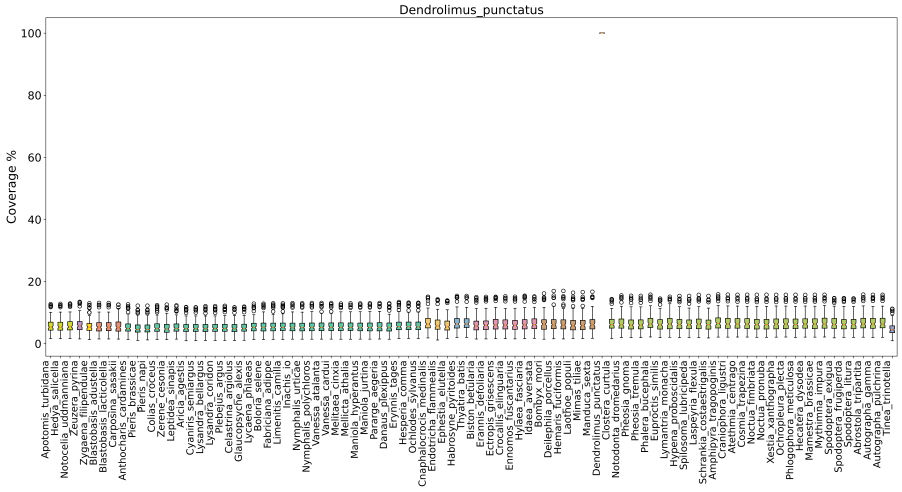
Dendrolimus_punctatus
Coverage %
100
80
60
40
20
0

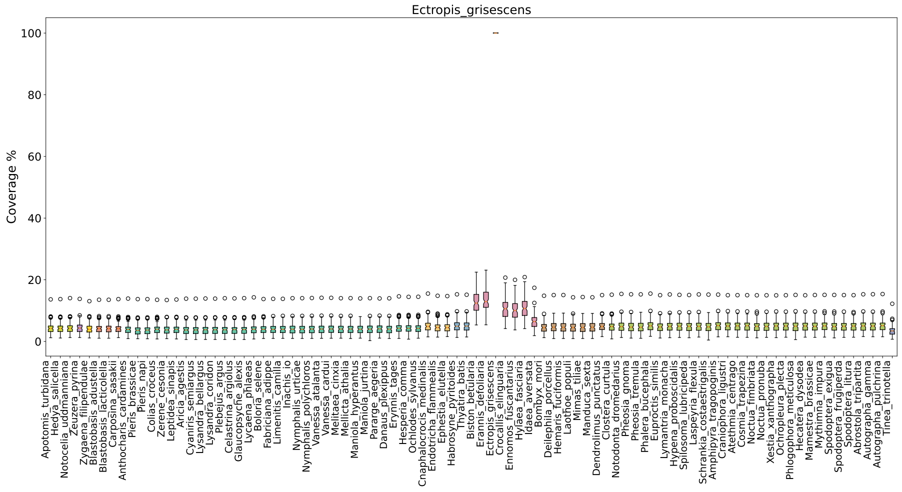
Ectropis_grisescens
Coverage %
100
80
60
40
20
0

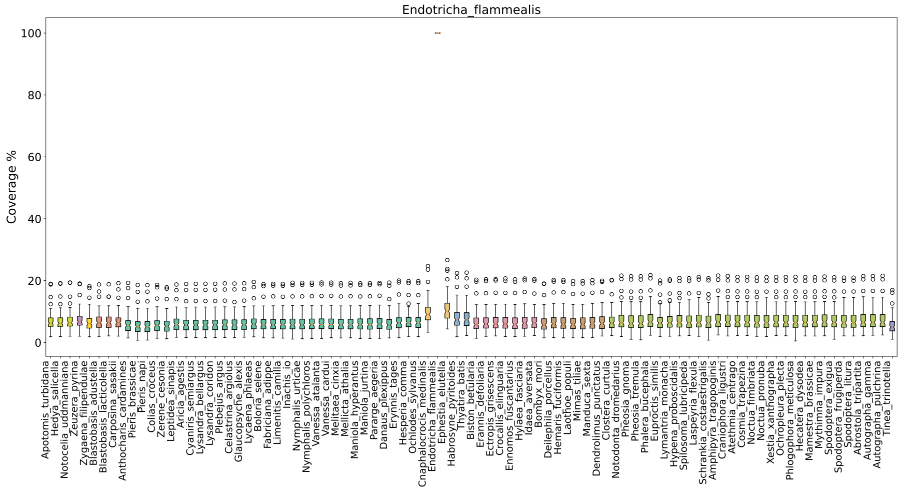
Endotricha_flammealis
Coverage %
100
80
60
40
20
0

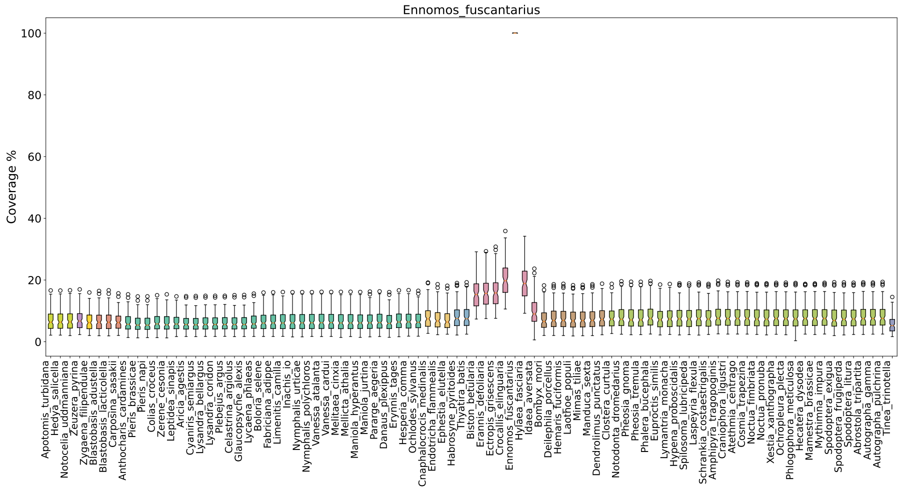
Ennomos_fuscantarius
Coverage %
0
20
40
60
80
100
Apotomis_turbidana
Hedya_salicella
Notocelia_uddmanniana
Zeuzera_pyrina
Zygaena_filipendulae
Blastobasis_adustella
Blastobasis_lacticolella
Carposina_sasakii
Anthocharis_cardamines
Pieris_brassicae
Pieris_napi
Colias_croceus
Zerene_cesonia
Leptidea_sinapis
Aricia_agestis
Cyaniris_semiargus
Lysandra_bellargus
Lysandra_coridon
Plebejus_argus
Celastrina_argiolus
Glaucopsyche_alexis
Lycaena_phlaeas
Boloria_selene
Fabriciana_adippe
Limenitis_camilla
Inachis_io
Nymphalis_urticae
Nymphalis_polychloros
Vanessa_atalanta
Vanessa_cardui
Melitaea_cinxia
Mellicta_athalia
Maniola_hyperantus
Maniola_jurtina
Pararge_aegeria
Danaus_plexippus
Erynnis_tages
Hesperia_comma
Ochlodes_sylvanus
Cnaphalocrocis_medinalis
Endotricha_flammealis
Ephestia_elutella
Habrosyne_pyritoides
Thyatira_batis
Biston_betularia
Erannis_defoliaria
Ectropis_grisescens
Crocallis_elinguaria
Ennomos_fuscantarius
Hylaea_fasciaria
Idaea_aversata
Bombyx_mori
Deilephila_porcellus
Hemaris_fuciformis
Laothoe_populi
Mimas_tiliae
Manduca_sexta
Dendrolimus_punctatus
Clostera_curtula
Notodonta_dromedarius
Pheosia_gnoma
Pheosia_tremula
Phalera_bucephala
Euproctis_similis
Lymantria_monacha
Hypena_proboscidalis
Spilosoma_lubricipeda
Laspeyria_flexula
Schrankia_costaestrigalis
Amphipyra_tragopoginis
Craniophora_ligustri
Atethmia_centrago
Cosmia_trapezina
Noctua_fimbriata
Noctua_pronuba
Xestia_xanthographa
Ochropleura_plecta
Phlogophora_meticulosa
Hecatera_dysodea
Mamestra_brassicae
Mythimna_impura
Spodoptera_exigua
Spodoptera_frugiperda
Spodoptera_litura
Abrostola_tripartita
Autographa_gamma
Autographa_pulchrina
Tinea_trinotella

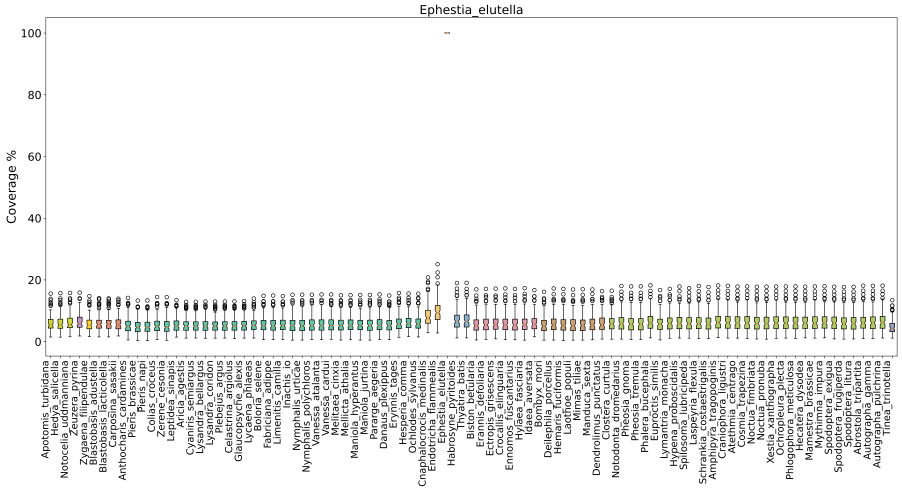
Ephestia_elutella
Coverage %
0
20
40
60
80
100
Apotomis_turbidana
Hedya_salicella
Notocelia_uddmanniana
Zeuzera_pyrina
Zygaena_filipendulae
Blastobasis_adustella
Blastobasis_lacticolella
Carposina_sasakii
Anthocharis_cardamines
Pieris_brassicae
Pieris_napi
Colias_croceus
Zerene_cesonia
Leptidea_sinapis
Aricia_agestis
Cyaniris_semiargus
Lysandra_bellargus
Lysandra_coridon
Plebejus_argus
Celastrina_argiolus
Glaucopsyche_alexis
Lycaena_phlaeas
Boloria_selene
Fabriciana_adippe
Limenitis_camilla
Inachis_io
Nymphalis_urticae
Nymphalis_polychloros
Vanessa_atalanta
Vanessa_cardui
Melitaea_cinxia
Mellicta_athalia
Maniola_hyperantus
Maniola_jurtina
Pararge_aegeria
Danaus_plexippus
Erynnis_tages
Hesperia_comma
Ochlodes_sylvanus
Cnaphalocrocis_medinalis
Endotricha_flammealis
Ephestia_elutella
Habrosyne_pyritoides
Thyatira_batis
Biston_betularia
Erannis_defoliaria
Ectropis_grisescens
Crocallis_elinguaria
Ennomos_fuscantarius
Hylaea_fasciaria
Idaea_aversata
Bombyx_mori
Deilephila_porcellus
Hemaris_fuciformis
Laothoe_populi
Mimas_tiliae
Manduca_sexta
Dendrolimus_punctatus
Clostera_curtula
Notodonta_dromedarius
Pheosia_gnoma
Pheosia_tremula
Phalera_bucephala
Euproctis_similis
Lymantria_monacha
Hypena_proboscidalis
Spilosoma_lubricipeda
Laspeyria_flexula
Schrankia_costaestrigalis
Amphipyra_tragopoginis
Craniophora_ligustri
Atethmia_centrago
Cosmia_trapezina
Noctua_fimbriata
Noctua_pronuba
Xestia_xanthographa
Ochropleura_plecta
Phlogophora_meticulosa
Hecatera_dysodea
Mamestra_brassicae
Mythimna_impura
Spodoptera_exigua
Spodoptera_frugiperda
Spodoptera_litura
Abrostola_tripartita
Autographa_gamma
Autographa_pulchrina
Tinea_trinotella

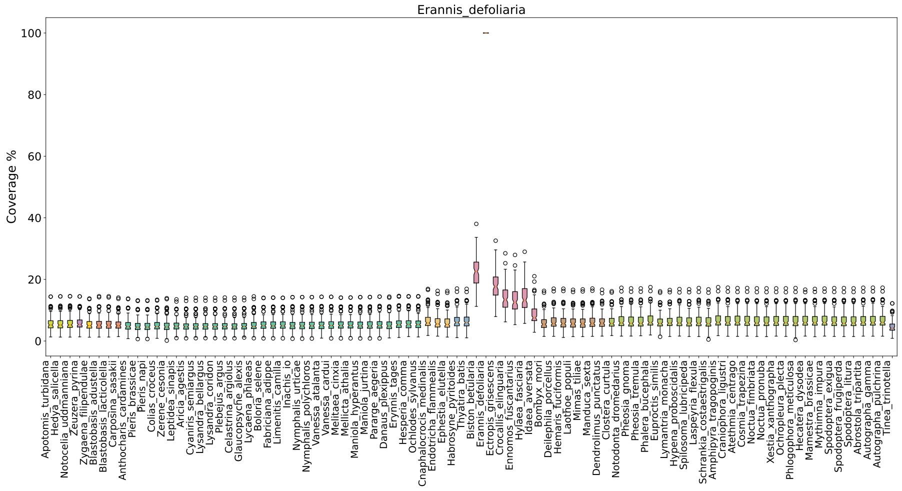
Erannis_defoliaria
Coverage %
0
20
40
60
80
100
Apotomis_turbidana
Hedya_salicella
Notocelia_uddmanniana
Zeuzera_pyrina
Zygaena_filipendulae
Blastobasis_adustella
Blastobasis_lacticolella
Carposina_sasakii
Anthocharis_cardamines
Pieris_brassicae
Pieris_napi
Colias_croceus
Zerene_cesonia
Leptidea_sinapis
Aricia_agestis
Cyaniris_semiargus
Lysandra_bellargus
Lysandra_coridon
Plebejus_argus
Celastrina_argiolus
Glaucopsyche_alexis
Lycaena_phlaeas
Boloria_selene
Fabriciana_adippe
Limenitis_camilla
Inachis_io
Nymphalis_urticae
Nymphalis_polychloros
Vanessa_atalanta
Vanessa_cardui
Melitaea_cinxia
Mellicta_athalia
Maniola_hyperantus
Maniola_jurtina
Pararge_aegeria
Danaus_plexippus
Erynnis_tages
Hesperia_comma
Ochlodes_sylvanus
Cnaphalocrocis_medinalis
Endotricha_flammealis
Ephestia_elutella
Habrosyne_pyritoides
Thyatira_batis
Biston_betularia
Erannis_defoliaria
Ectropis_grisescens
Crocallis_elinguaria
Ennomos_fuscantarius
Hylaea_fasciaria
Idaea_aversata
Bombyx_mori
Deilephila_porcellus
Hemaris_fuciformis
Laothoe_populi
Mimas_tiliae
Manduca_sexta
Dendrolimus_punctatus
Clostera_curtula
Notodonta_dromedarius
Pheosia_gnoma
Pheosia_tremula
Phalera_bucephala
Euproctis_similis
Lymantria_monacha
Hypena_proboscidalis
Spilosoma_lubricipeda
Laspeyria_flexula
Schrankia_costaestrigalis
Amphipyra_tragopoginis
Craniophora_ligustri
Atethmia_centrago
Cosmia_trapezina
Noctua_fimbriata
Noctua_pronuba
Xestia_xanthographa
Ochropleura_plecta
Phlogophora_meticulosa
Hecatera_dysodea
Mamestra_brassicae
Mythimna_impura
Spodoptera_exigua
Spodoptera_frugiperda
Spodoptera_litura
Abrostola_tripartita
Autographa_gamma
Autographa_pulchrina
Tinea_trinotella

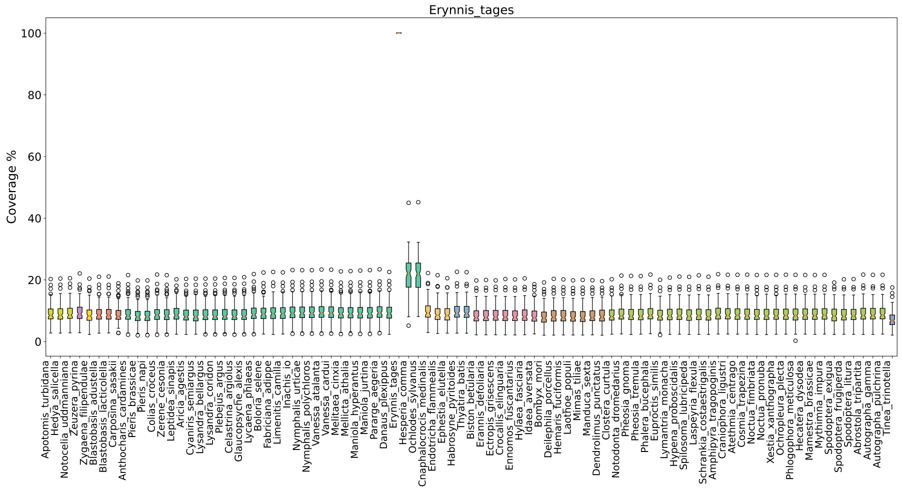
Erynnis_tages
Coverage %
0
20
40
60
80
100

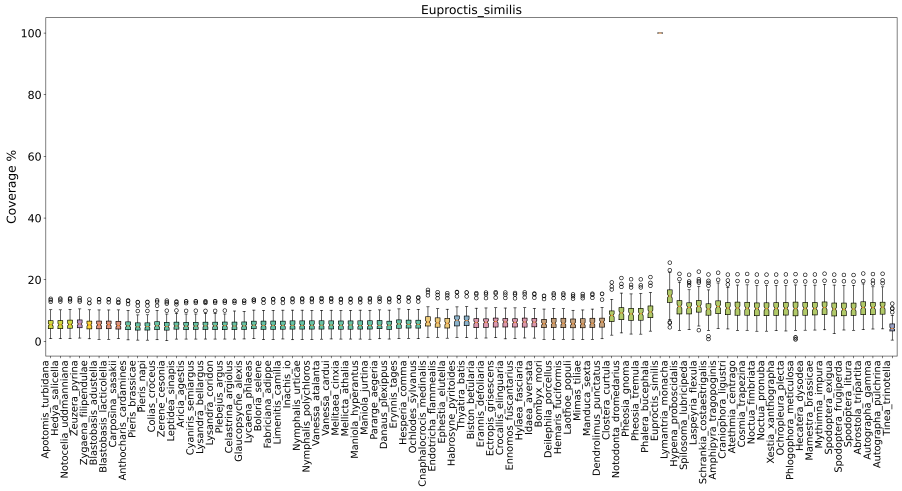
Euproctis_similis
Coverage %
0
20
40
60
80
100

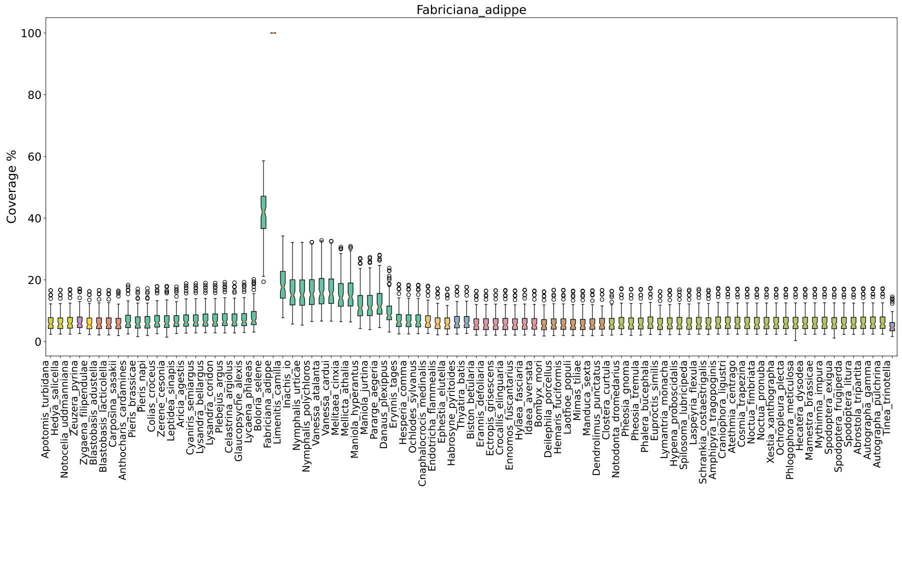
Fabriciana_adippe
Coverage %
0
20
40
60
80
100

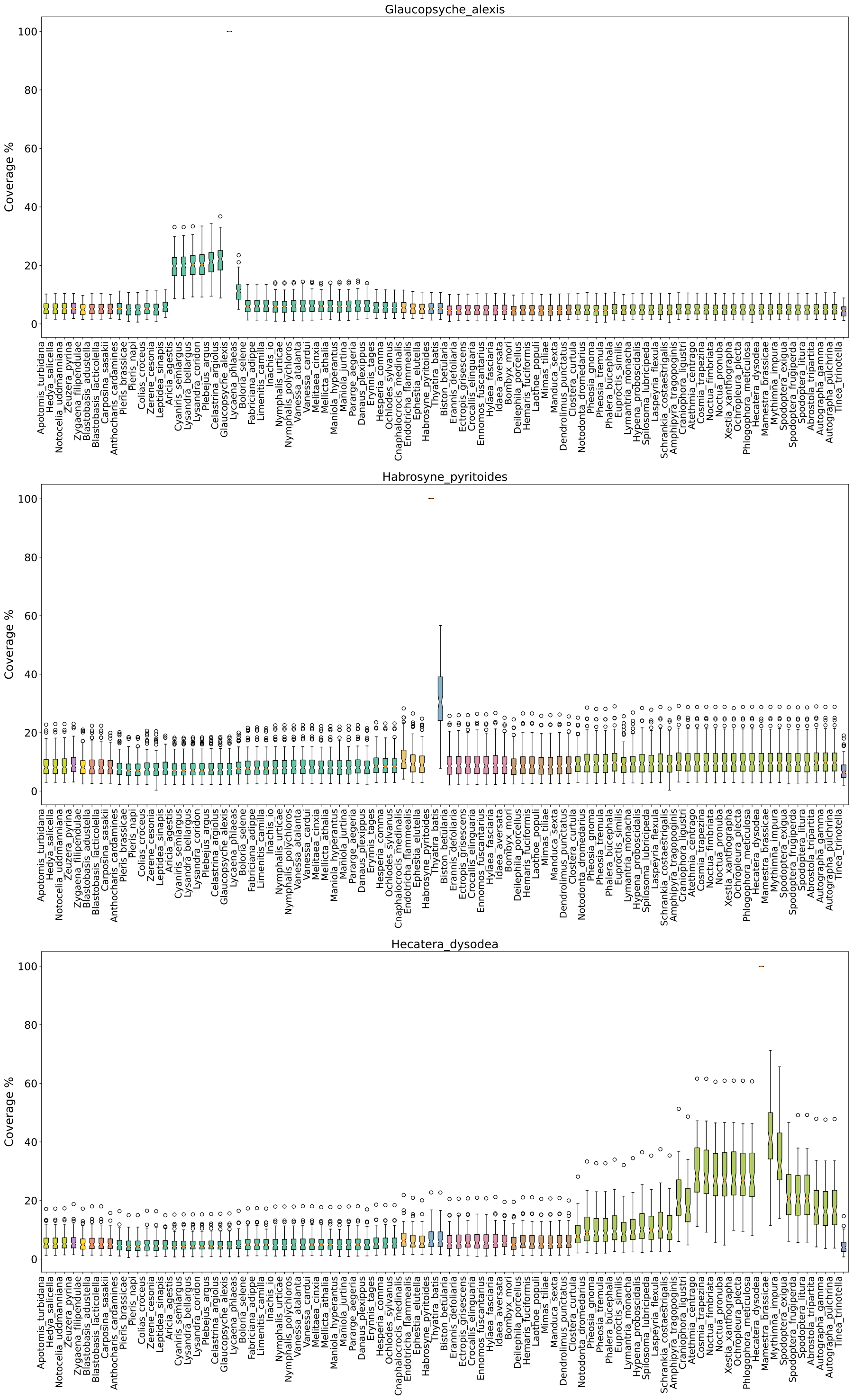
Glaucopsyche_alexis
Coverage %
0
20
40
60
80
100
Habrosyne_pyritoides
Hecatera_dysodea

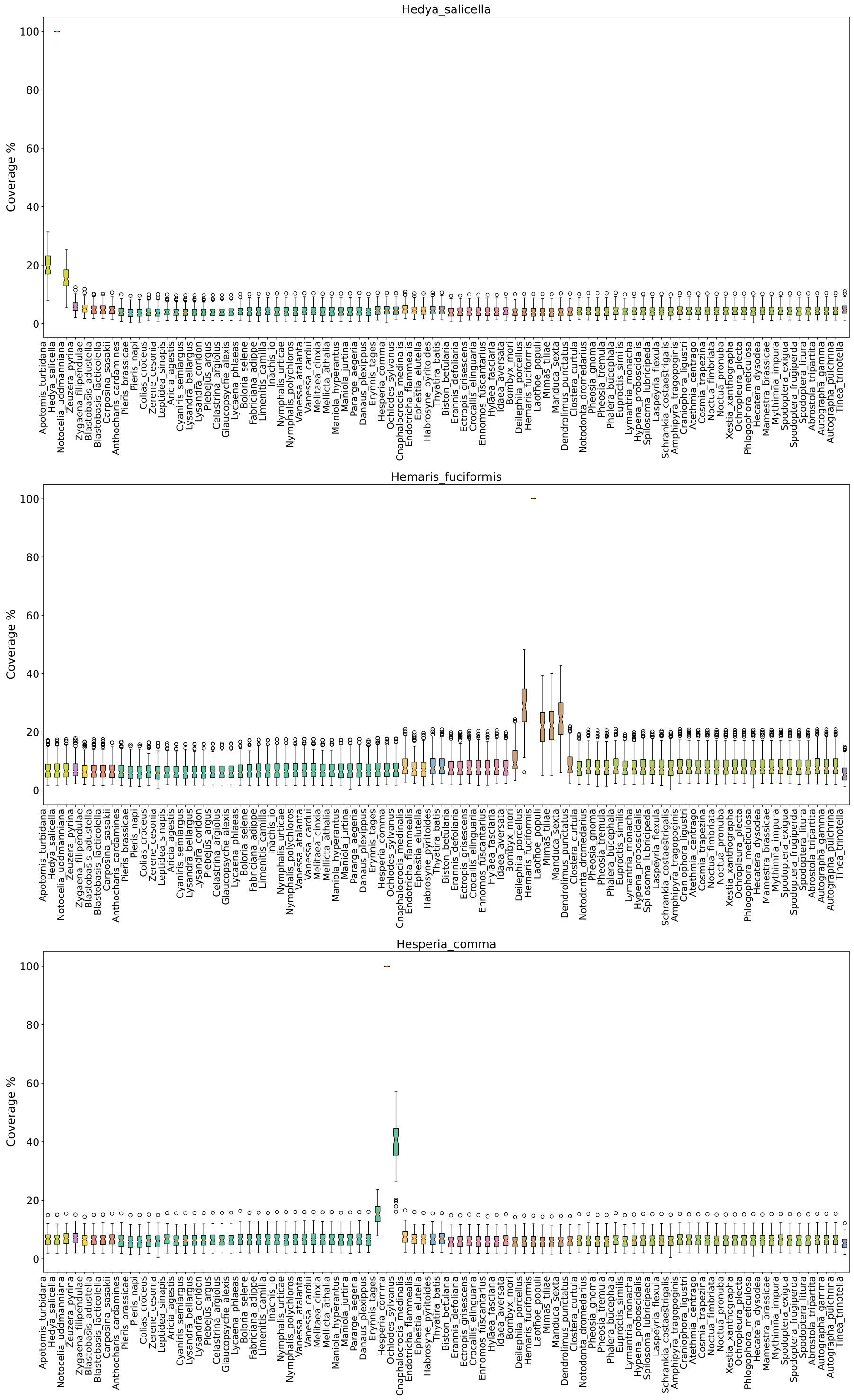
Hedya_salicella
Hemaris_fuciformis
Hesperia_comma
Coverage %
0
20
40
60
80
100
Apotomis_turbidana
Hedya_salicella
Notocelia_uddmanniana
Zeuzera_pyrina
Zygaena_filipendulae
Blastobasis_adustella
Blastobasis_lacticolella
Carposina_sasakii
Anthocharis_cardamines
Pieris_brassicae
Pieris_napi
Colias_croceus
Zerene_cesonia
Leptidea_sinapis
Aricia_agestis
Cyaniris_semiargus
Lysandra_bellargus
Lysandra_coridon
Plebejus_argus
Celastrina_argiolus
Glaucopsyche_alexis
Lycaena_phlaeas
Boloria_selene
Fabriciana_adippe
Limenitis_camilla
Inachis_io
Nymphalis_urticae
Nymphalis_polychloros
Vanessa_atalanta
Vanessa_cardui
Melitaea_cinxia
Mellicta_athalia
Maniola_hyperantus
Maniola_jurtina
Pararge_aegeria
Danaus_plexippus
Erynnis_tages
Hesperia_comma
Ochlodes_sylvanus
Cnaphalocrocis_medinalis
Endotricha_flammealis
Ephestia_elutella
Habrosyne_pyritoides
Thyatira_batis
Biston_betularia
Erannis_defoliaria
Ectropis_grisescens
Crocallis_elinguaria
Ennomos_fuscantarius
Hylaea_fasciaria
Idaea_aversata
Bombyx_mori
Deilephila_porcellus
Hemaris_fuciformis
Laothoe_populi
Mimas_tiliae
Manduca_sexta
Dendrolimus_punctatus
Clostera_curtula
Notodonta_dromedarius
Pheosia_gnoma
Pheosia_tremula
Phalera_bucephala
Euproctis_similis
Lymantria_monacha
Hypena_proboscidalis
Spilosoma_lubricipeda
Laspeyria_flexula
Schrankia_costaestrigalis
Amphipyra_tragopoginis
Craniophora_ligustri
Atethmia_centrago
Cosmia_trapezina
Noctua_fimbriata
Noctua_pronuba
Xestia_xanthographa
Ochropleura_plecta
Phlogophora_meticulosa
Hecatera_dysodea
Mamestra_brassicae
Mythimna_impura
Spodoptera_exigua
Spodoptera_frugiperda
Spodoptera_litura
Abrostola_tripartita
Autographa_gamma
Autographa_pulchrina
Tinea_trinotella

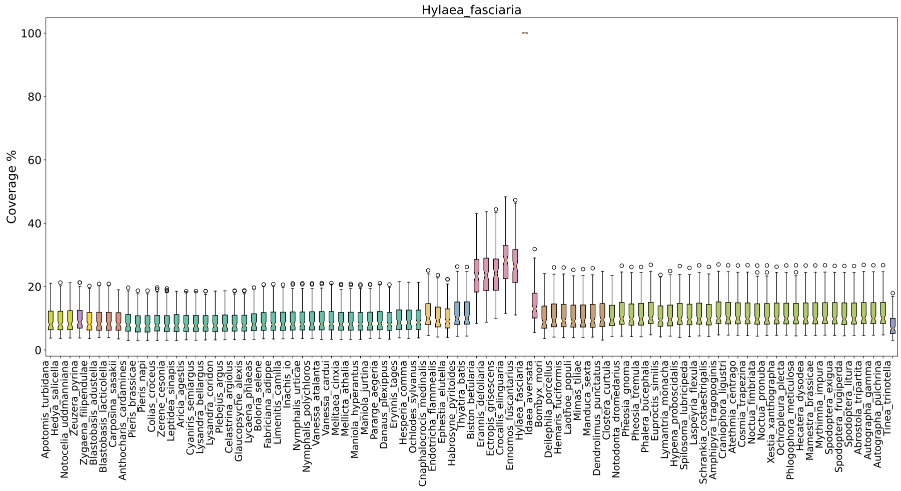
Hylaea_fasciaria
Coverage %
0
20
40
60
80
100
Apotomis_turbidana
Hedya_salicella
Notocelia_uddmanniana
Zeuzera_pyrina
Zygaena_filipendulae
Blastobasis_adustella
Blastobasis_lacticolella
Carposina_sasakii
Anthocharis_cardamines
Pieris_brassicae
Pieris_napi
Colias_croceus
Zerene_cesonia
Leptidea_sinapis
Aricia_agestis
Cyaniris_semiargus
Lysandra_bellargus
Lysandra_coridon
Plebejus_argus
Celastrina_argiolus
Glaucopsyche_alexis
Lycaena_phlaeas
Boloria_selene
Fabriciana_adippe
Limenitis_camilla
Inachis_io
Nymphalis_urticae
Nymphalis_polychloros
Vanessa_atalanta
Vanessa_cardui
Melitaea_cinxia
Mellicta_athalia
Maniola_hyperantus
Maniola_jurtina
Pararge_aegeria
Danaus_plexippus
Erynnis_tages
Hesperia_comma
Ochlodes_sylvanus
Cnaphalocrocis_medinalis
Endotricha_flammealis
Ephestia_elutella
Habrosyne_pyritoides
Thyatira_batis
Biston_betularia
Erannis_defoliaria
Ectropis_grisescens
Crocallis_elinguaria
Ennomos_fuscantarius
Hylaea_fasciaria
Idaea_aversata
Bombyx_mori
Deilephila_porcellus
Hemaris_fuciformis
Laothoe_populi
Mimas_tiliae
Manduca_sexta
Dendrolimus_punctatus
Clostera_curtula
Notodonta_dromedarius
Pheosia_gnoma
Pheosia_tremula
Phalera_bucephala
Euproctis_similis
Lymantria_monacha
Hypena_proboscidalis
Spilosoma_lubricipeda
Laspeyria_flexula
Schrankia_costaestrigalis
Amphipyra_tragopoginis
Craniophora_ligustri
Atethmia_centrago
Cosmia_trapezina
Noctua_fimbriata
Noctua_pronuba
Xestia_xanthographa
Ochropleura_plecta
Phlogophora_meticulosa
Hecatera_dysodea
Mamestra_brassicae
Mythimna_impura
Spodoptera_exigua
Spodoptera_frugiperda
Spodoptera_litura
Abrostola_tripartita
Autographa_gamma
Autographa_pulchrina
Tinea_trinotella
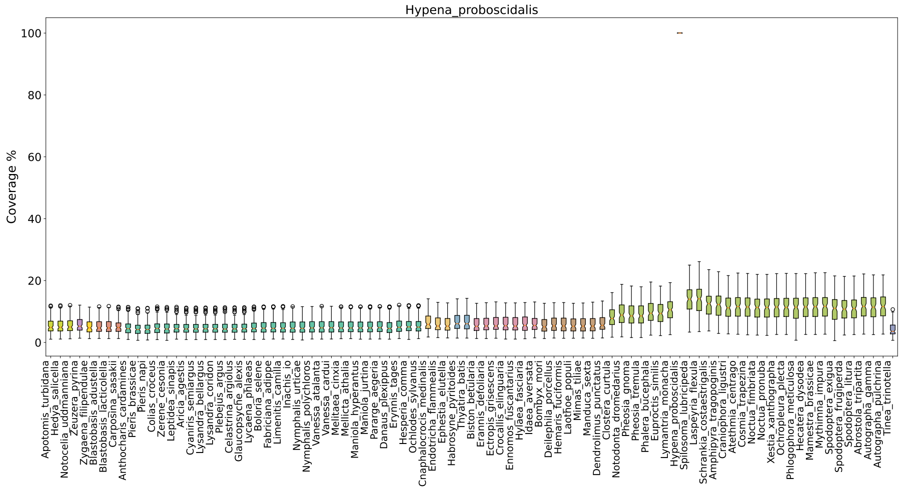
Hypena_proboscidalis
Coverage %
0
20
40
60
80
100
Apotomis_turbidana
Hedya_salicella
Notocelia_uddmanniana
Zeuzera_pyrina
Zygaena_filipendulae
Blastobasis_adustella
Blastobasis_lacticolella
Carposina_sasakii
Anthocharis_cardamines
Pieris_brassicae
Pieris_napi
Colias_croceus
Zerene_cesonia
Leptidea_sinapis
Aricia_agestis
Cyaniris_semiargus
Lysandra_bellargus
Lysandra_coridon
Plebejus_argus
Celastrina_argiolus
Glaucopsyche_alexis
Lycaena_phlaeas
Boloria_selene
Fabriciana_adippe
Limenitis_camilla
Inachis_io
Nymphalis_urticae
Nymphalis_polychloros
Vanessa_atalanta
Vanessa_cardui
Melitaea_cinxia
Mellicta_athalia
Maniola_hyperantus
Maniola_jurtina
Pararge_aegeria
Danaus_plexippus
Erynnis_tages
Hesperia_comma
Ochlodes_sylvanus
Cnaphalocrocis_medinalis
Endotricha_flammealis
Ephestia_elutella
Habrosyne_pyritoides
Thyatira_batis
Biston_betularia
Erannis_defoliaria
Ectropis_grisescens
Crocallis_elinguaria
Ennomos_fuscantarius
Hylaea_fasciaria
Idaea_aversata
Bombyx_mori
Deilephila_porcellus
Hemaris_fuciformis
Laothoe_populi
Mimas_tiliae
Manduca_sexta
Dendrolimus_punctatus
Clostera_curtula
Notodonta_dromedarius
Pheosia_gnoma
Pheosia_tremula
Phalera_bucephala
Euproctis_similis
Lymantria_monacha
Hypena_proboscidalis
Spilosoma_lubricipeda
Laspeyria_flexula
Schrankia_costaestrigalis
Amphipyra_tragopoginis
Craniophora_ligustri
Atethmia_centrago
Cosmia_trapezina
Noctua_fimbriata
Noctua_pronuba
Xestia_xanthographa
Ochropleura_plecta
Phlogophora_meticulosa
Hecatera_dysodea
Mamestra_brassicae
Mythimna_impura
Spodoptera_exigua
Spodoptera_frugiperda
Spodoptera_litura
Abrostola_tripartita
Autographa_gamma
Autographa_pulchrina
Tinea_trinotella
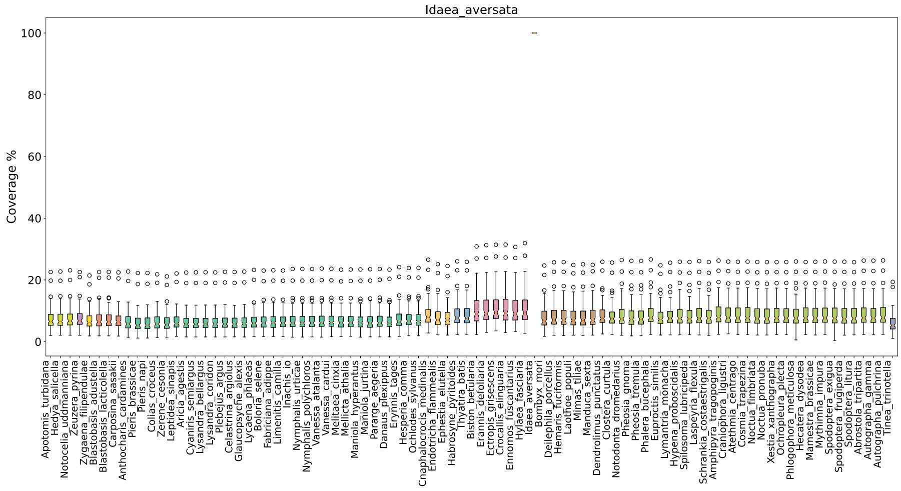
Idaea_aversata
Coverage %
0
20
40
60
80
100
Apotomis_turbidana
Hedya_salicella
Notocelia_uddmanniana
Zeuzera_pyrina
Zygaena_filipendulae
Blastobasis_adustella
Blastobasis_lacticolella
Carposina_sasakii
Anthocharis_cardamines
Pieris_brassicae
Pieris_napi
Colias_croceus
Zerene_cesonia
Leptidea_sinapis
Aricia_agestis
Cyaniris_semiargus
Lysandra_bellargus
Lysandra_coridon
Plebejus_argus
Celastrina_argiolus
Glaucopsyche_alexis
Lycaena_phlaeas
Boloria_selene
Fabriciana_adippe
Limenitis_camilla
Inachis_io
Nymphalis_urticae
Nymphalis_polychloros
Vanessa_atalanta
Vanessa_cardui
Melitaea_cinxia
Mellicta_athalia
Maniola_hyperantus
Maniola_jurtina
Pararge_aegeria
Danaus_plexippus
Erynnis_tages
Hesperia_comma
Ochlodes_sylvanus
Cnaphalocrocis_medinalis
Endotricha_flammealis
Ephestia_elutella
Habrosyne_pyritoides
Thyatira_batis
Biston_betularia
Erannis_defoliaria
Ectropis_grisescens
Crocallis_elinguaria
Ennomos_fuscantarius
Hylaea_fasciaria
Idaea_aversata
Bombyx_mori
Deilephila_porcellus
Hemaris_fuciformis
Laothoe_populi
Mimas_tiliae
Manduca_sexta
Dendrolimus_punctatus
Clostera_curtula
Notodonta_dromedarius
Pheosia_gnoma
Pheosia_tremula
Phalera_bucephala
Euproctis_similis
Lymantria_monacha
Hypena_proboscidalis
Spilosoma_lubricipeda
Laspeyria_flexula
Schrankia_costaestrigalis
Amphipyra_tragopoginis
Craniophora_ligustri
Atethmia_centrago
Cosmia_trapezina
Noctua_fimbriata
Noctua_pronuba
Xestia_xanthographa
Ochropleura_plecta
Phlogophora_meticulosa
Hecatera_dysodea
Mamestra_brassicae
Mythimna_impura
Spodoptera_exigua
Spodoptera_frugiperda
Spodoptera_litura
Abrostola_tripartita
Autographa_gamma
Autographa_pulchrina
Tinea_trinotella

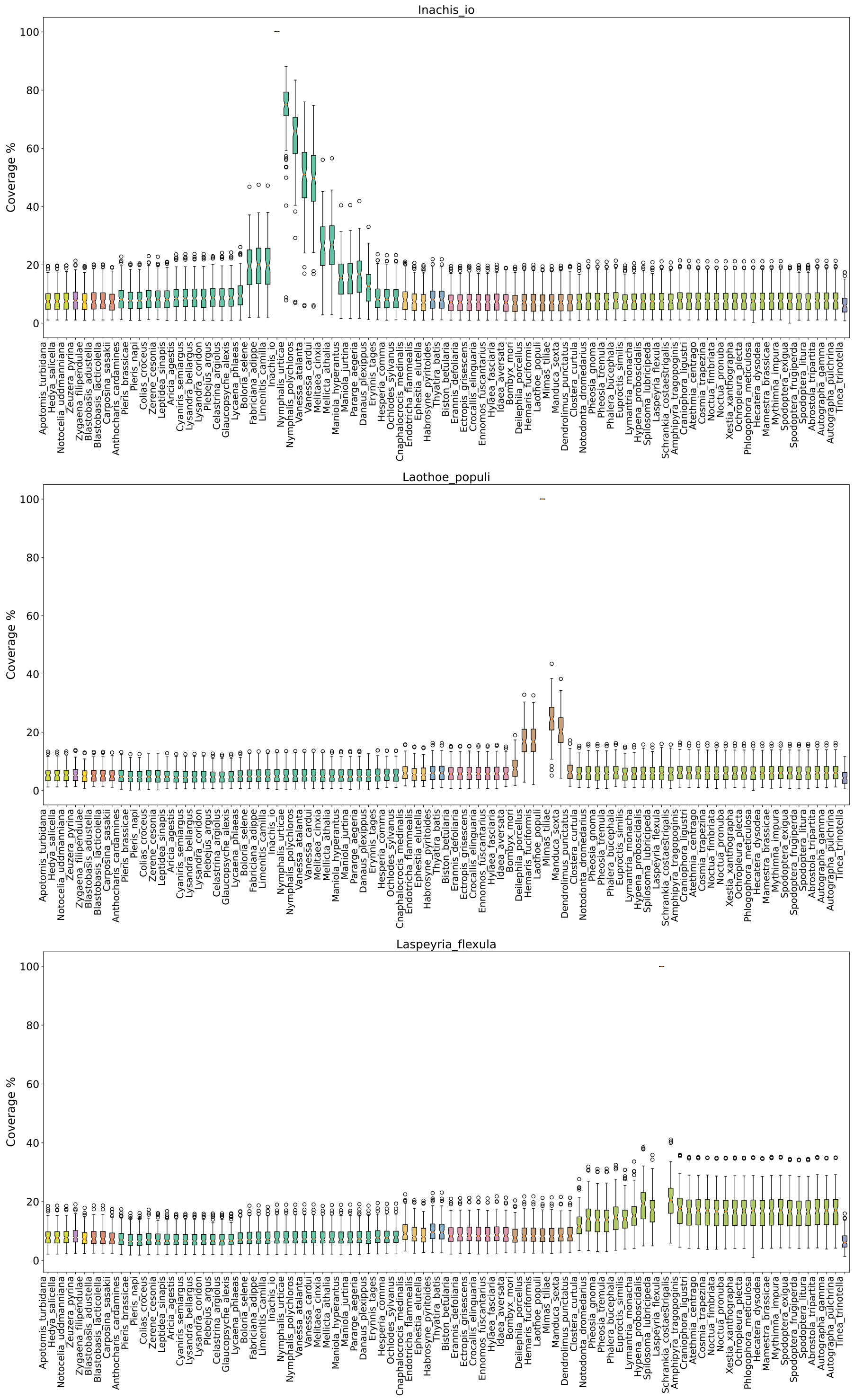
Inachis_io
Coverage %
Laothoe_populi
Coverage %
Laspeyria_flexula
Coverage %

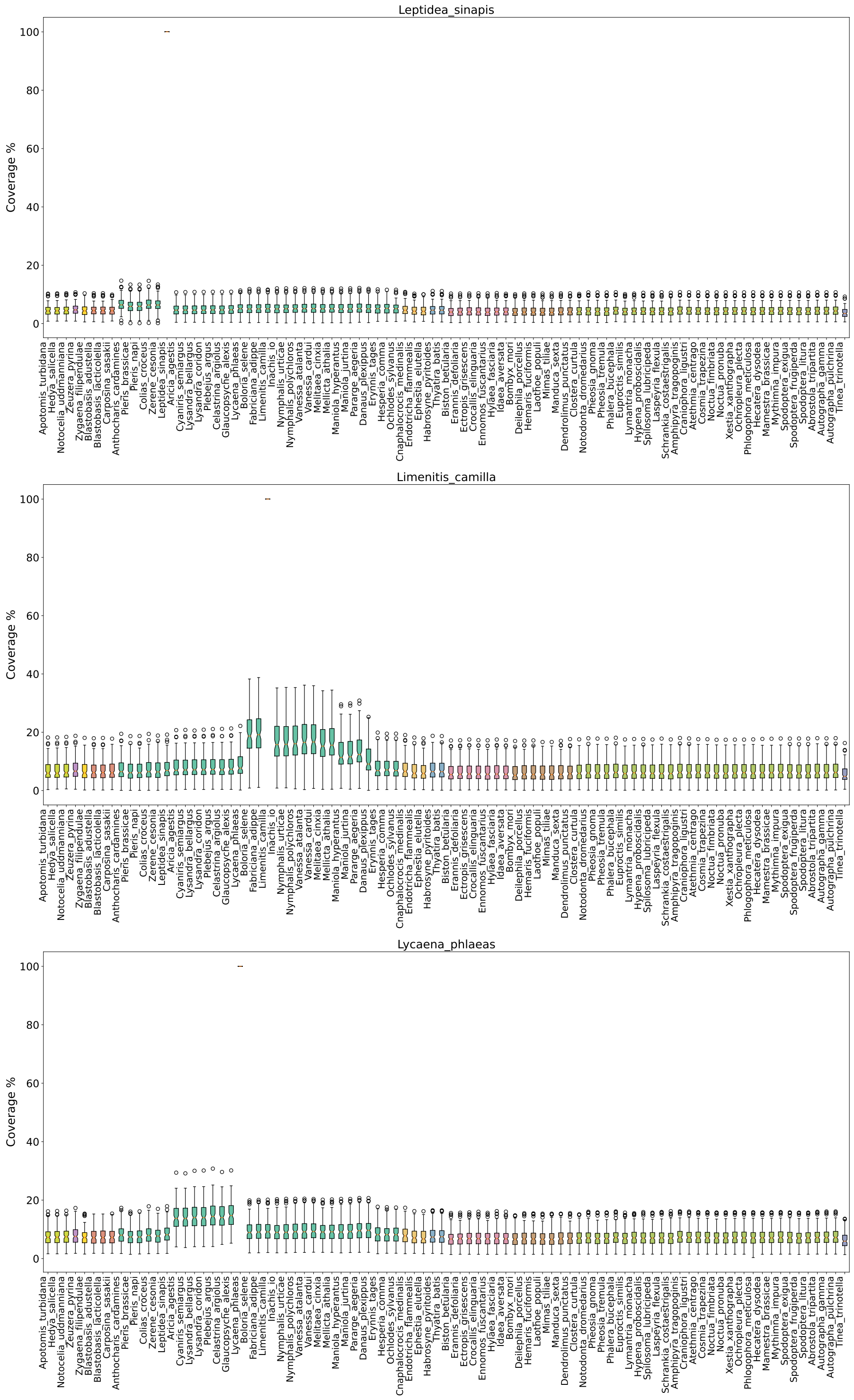
Leptidea_sinapis
Coverage %
0
20
40
60
80
100
Limenitis_camilla
Coverage %
0
20
40
60
80
100
Lycaena_phlaeas
Coverage %
0
20
40
60
80
100

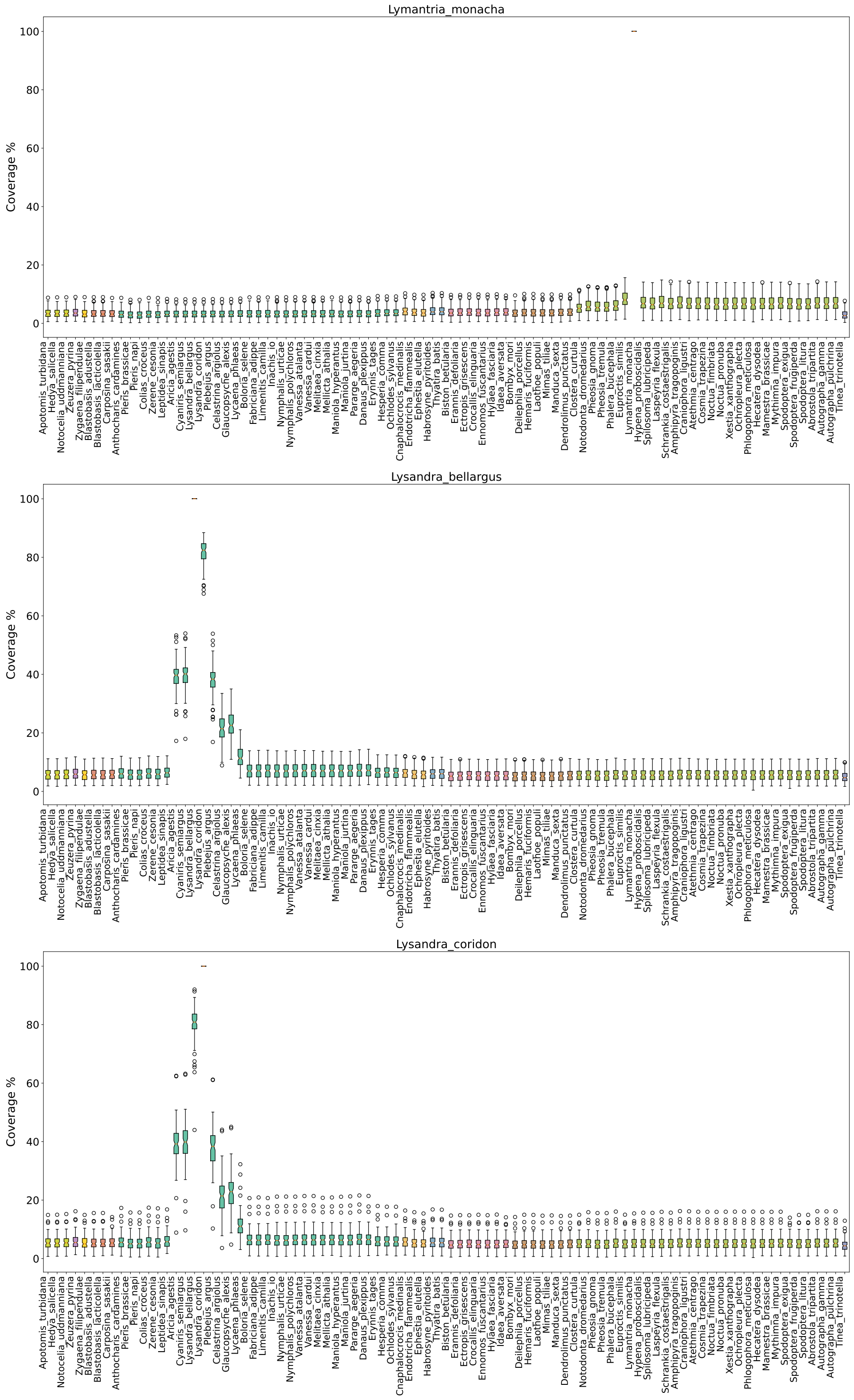
Lymantria_monacha
Lysandra_bellargus
Lysandra_coridon
Coverage %
0
20
40
60
80
100
Apotomis_turbidana
Hedya_salicella
Notocelia_uddmanniana
Zeuzera_pyrina
Zygaena_filipendulae
Blastobasis_adustella
Blastobasis_lacticolella
Carposina_sasakii
Anthocharis_cardamines
Pieris_brassicae
Pieris_napi
Colias_croceus
Zerene_cesonia
Leptidea_sinapis
Aricia_agestis
Cyaniris_semiargus
Lysandra_bellargus
Lysandra_coridon
Plebejus_argus
Celastrina_argiolus
Glaucopsyche_alexis
Lycaena_phlaeas
Boloria_selene
Fabriciana_adippe
Limenitis_camilla
Inachis_io
Nymphalis_urticae
Nymphalis_polychloros
Vanessa_atalanta
Vanessa_cardui
Melitaea_cinxia
Mellicta_athalia
Maniola_hyperantus
Maniola_jurtina
Pararge_aegeria
Danaus_plexippus
Erynnis_tages
Hesperia_comma
Ochlodes_sylvanus
Cnaphalocrocis_medinalis
Endotricha_flammealis
Ephestia_elutella
Habrosyne_pyritoides
Thyatira_batis
Biston_betularia
Erannis_defoliaria
Ectropis_grisescens
Crocallis_elinguaria
Ennomos_fuscantarius
Hylaea_fasciaria
Idaea_aversata
Bombyx_mori
Deilephila_porcellus
Hemaris_fuciformis
Laothoe_populi
Mimas_tiliae
Manduca_sexta
Dendrolimus_punctatus
Clostera_curtula
Notodonta_dromedarius
Pheosia_gnoma
Pheosia_tremula
Phalera_bucephala
Euproctis_similis
Lymantria_monacha
Hypena_proboscidalis
Spilosoma_lubricipeda
Laspeyria_flexula
Schrankia_costaestrigalis
Amphipyra_tragopoginis
Craniophora_ligustri
Atethmia_centrago
Cosmia_trapezina
Noctua_fimbriata
Noctua_pronuba
Xestia_xanthographa
Ochropleura_plecta
Phlogophora_meticulosa
Hecatera_dysodea
Mamestra_brassicae
Mythimna_impura
Spodoptera_exigua
Spodoptera_frugiperda
Spodoptera_litura
Abrostola_tripartita
Autographa_gamma
Autographa_pulchrina
Tinea_trinotella

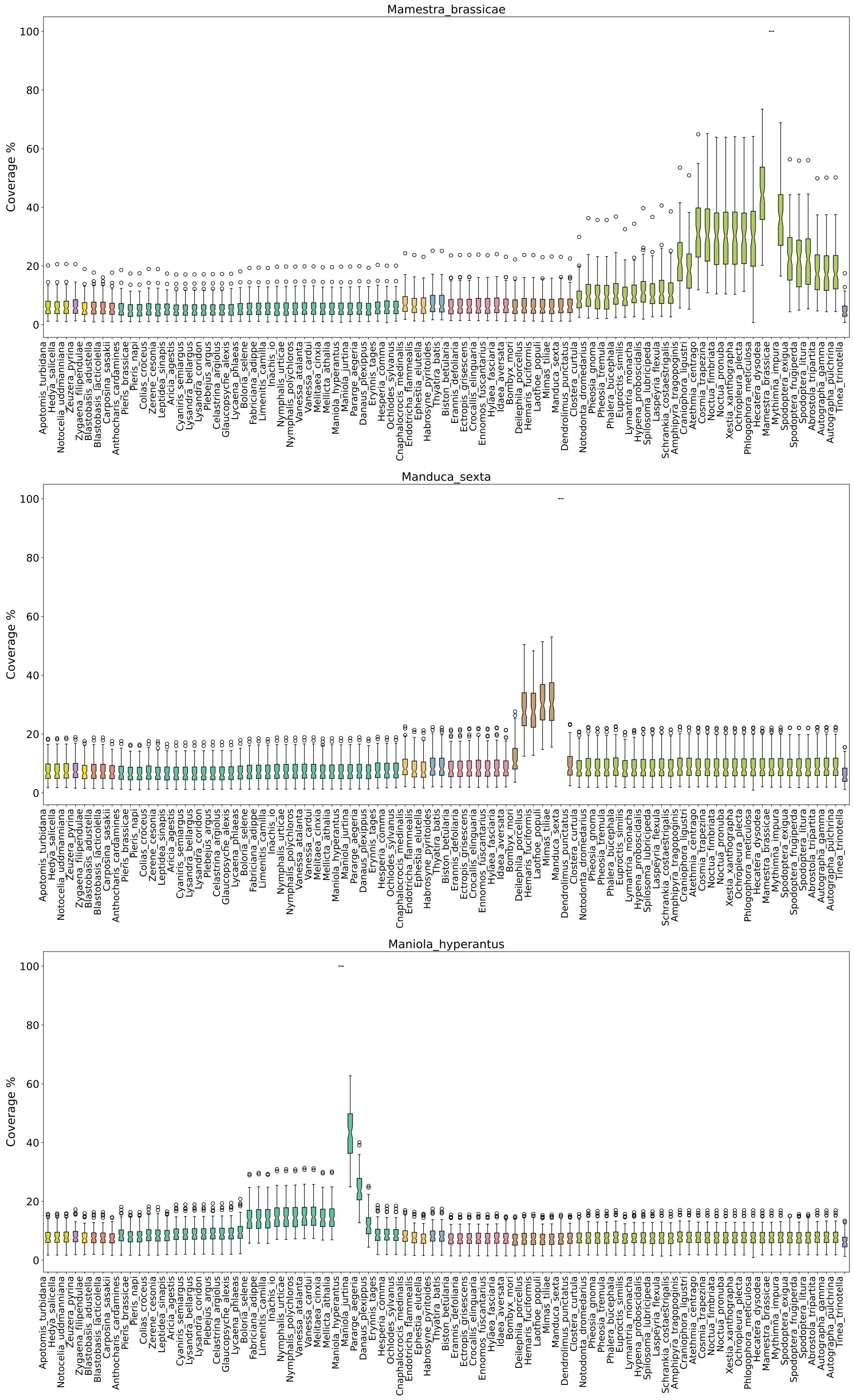
Mamestra_brassicae
Coverage %
Manduca_sexta
Coverage %
Maniola_hyperantus
Coverage %

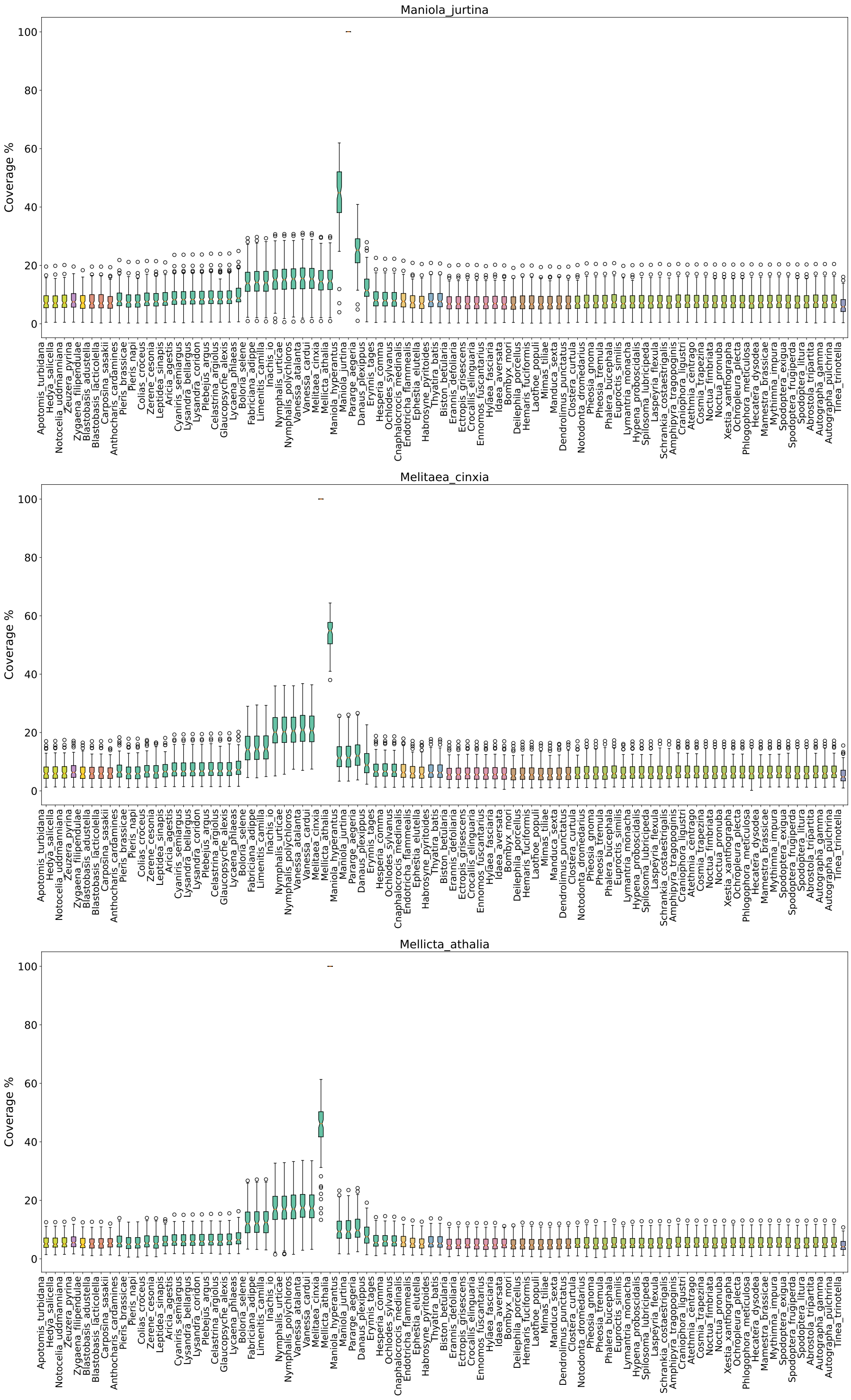
Maniola_jurtina
Coverage %
0
20
40
60
80
100
Melitaea_cinxia
Coverage %
0
20
40
60
80
100
Mellicta_athalia
Coverage %
0
20
40
60
80
100

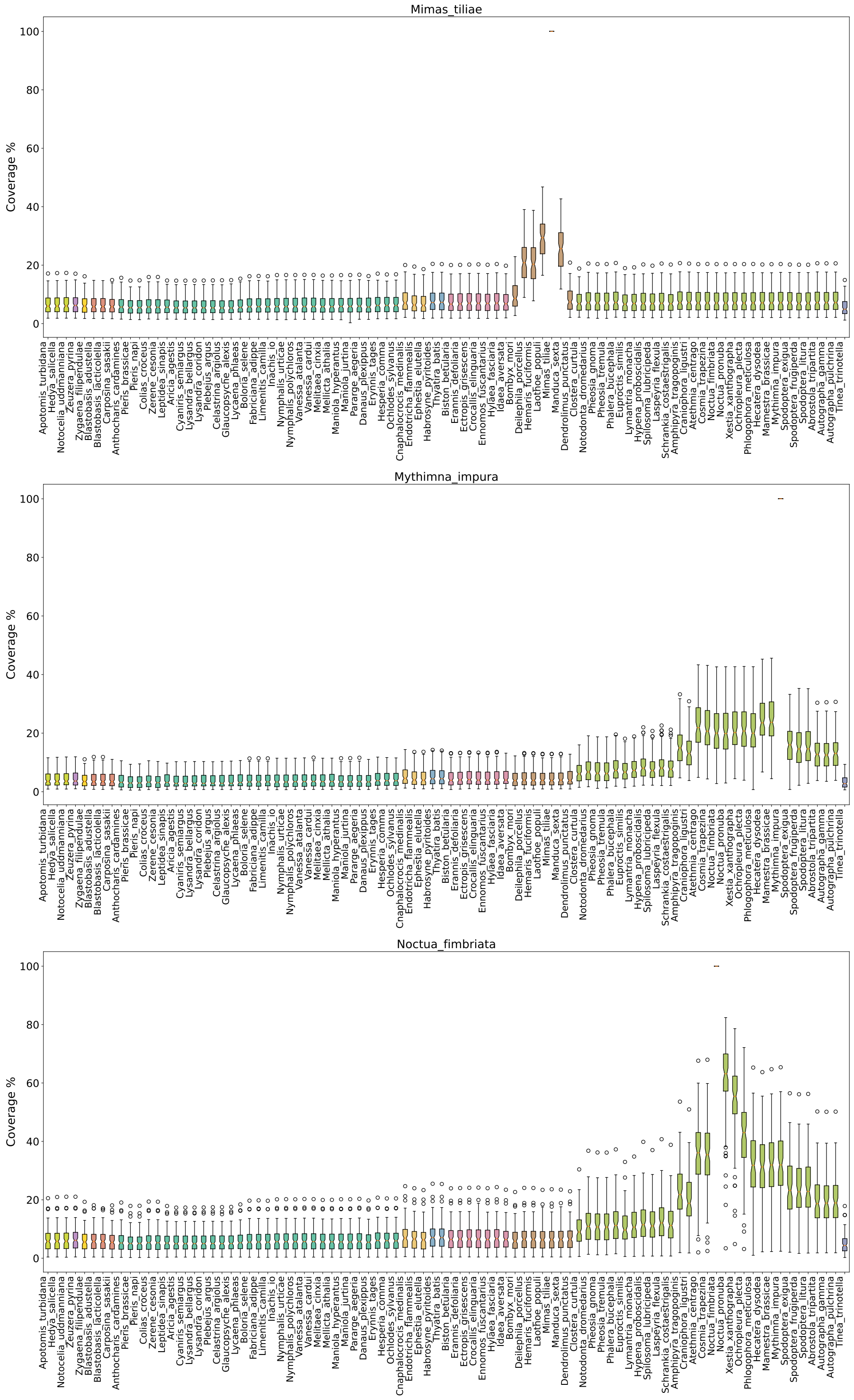
Mimas_tiliae
Coverage %
0
20
40
60
80
100
Mythimna_impura
Coverage %
0
20
40
60
80
100
Noctua_fimbriata
Coverage %
0
20
40
60
80
100

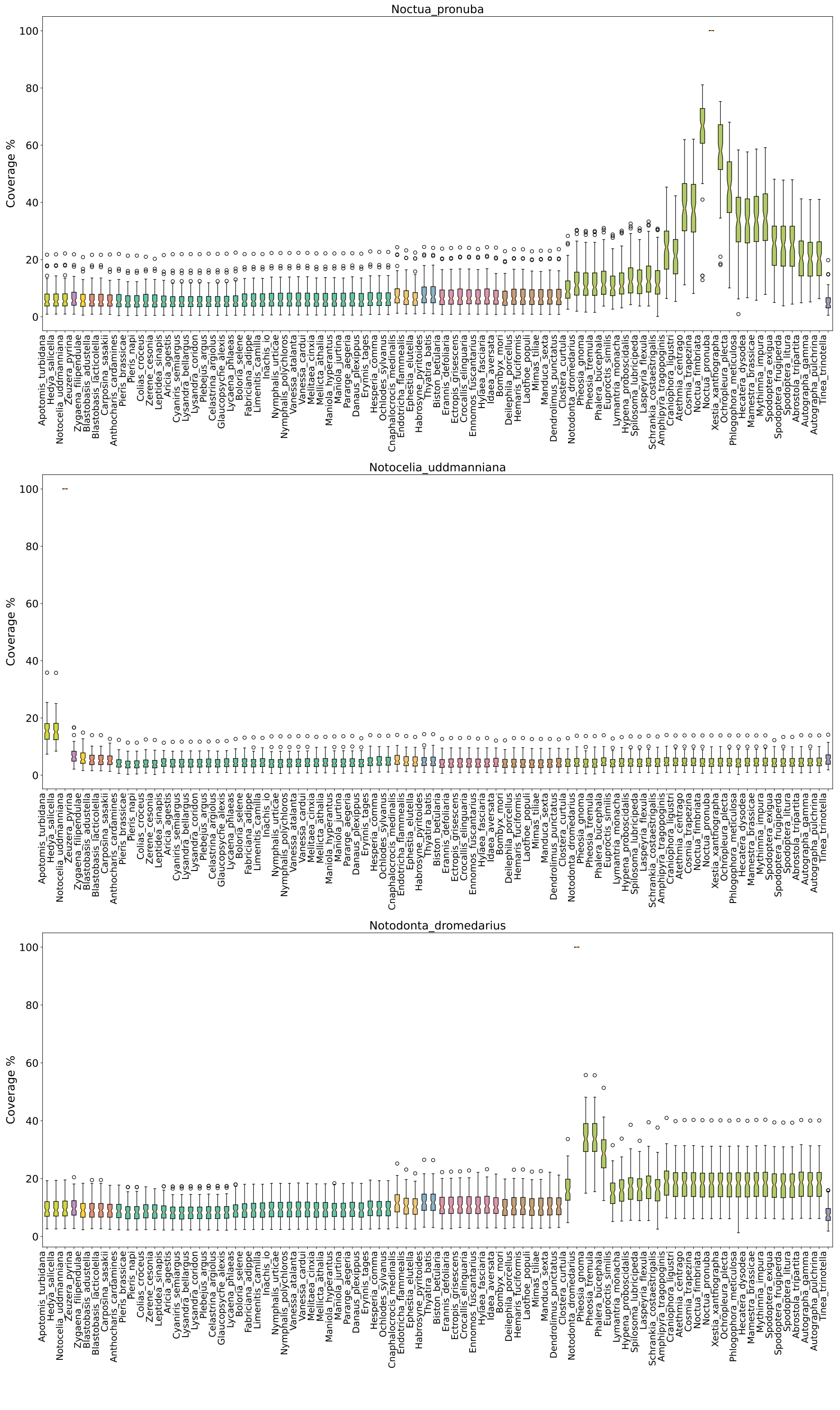

Noctua_pronuba
Coverage %
Notocelia_uddmanniana
Coverage %
Notodonta_dromedarius
Coverage %

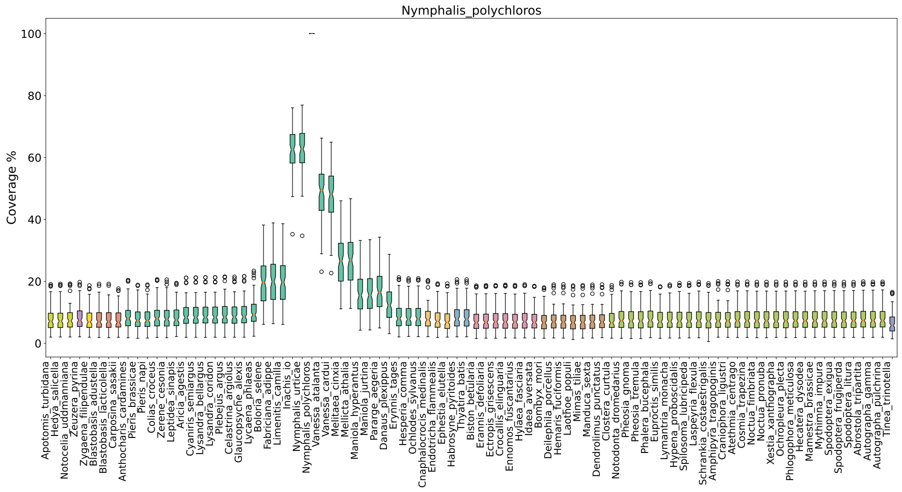
Nymphalis_polychloros
Coverage %
0
20
40
60
80
100

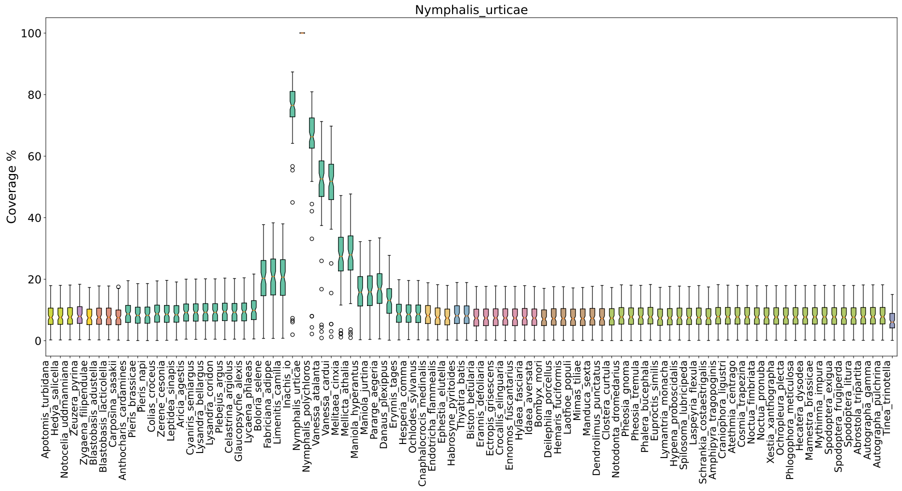
Nymphalis_urticae
Coverage %
0
20
40
60
80
100

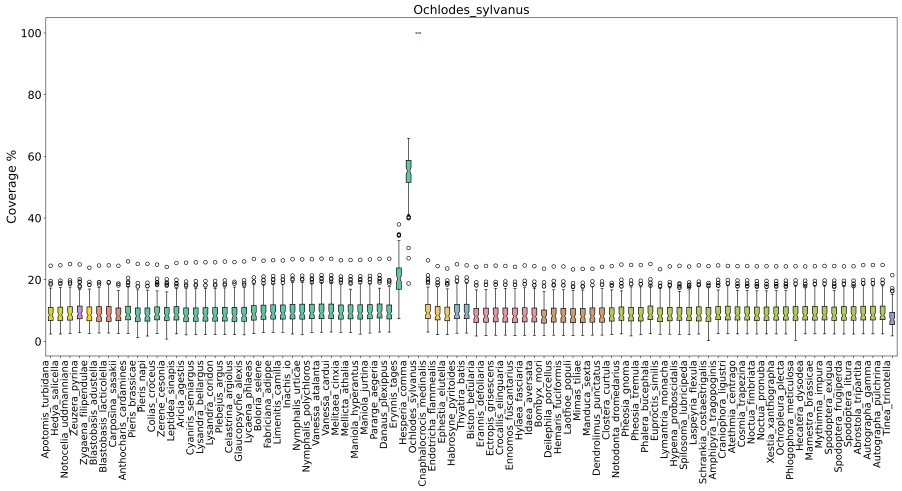
Ochlodes_sylvanus
Coverage %
0
20
40
60
80
100

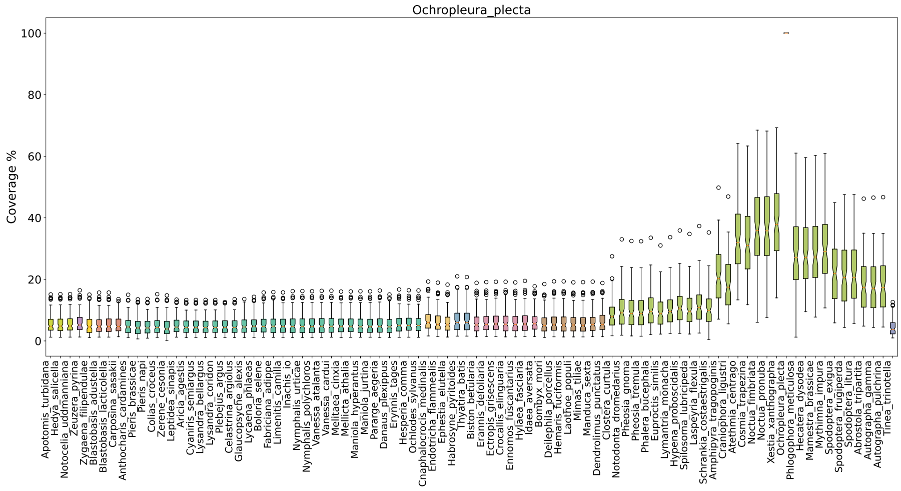
Ochropleura_plecta
Coverage %
0
20
40
60
80
100
Apotomis_turbidana
Hedya_salicella
Notocelia_uddmanniana
Zeuzera_pyrina
Zygaena_filipendulae
Blastobasis_adustella
Blastobasis_lacticolella
Carposina_sasakii
Anthocharis_cardamines
Pieris_brassicae
Pieris_napi
Colias_croceus
Zerene_cesonia
Leptidea_sinapis
Aricia_agestis
Cyaniris_semiargus
Lysandra_bellargus
Lysandra_coridon
Plebejus_argus
Celastrina_argiolus
Glaucopsyche_alexis
Lycaena_phlaeas
Boloria_selene
Fabriciana_adippe
Limenitis_camilla
Inachis_io
Nymphalis_urticae
Nymphalis_polychloros
Vanessa_atalanta
Vanessa_cardui
Melitaea_cinxia
Mellicta_athalia
Maniola_hyperantus
Maniola_jurtina
Pararge_aegeria
Danaus_plexippus
Erynnis_tages
Hesperia_comma
Ochlodes_sylvanus
Cnaphalocrocis_medinalis
Endotricha_flammealis
Ephestia_elutella
Habrosyne_pyritoides
Thyatira_batis
Biston_betularia
Erannis_defoliaria
Ectropis_grisescens
Crocallis_elinguaria
Ennomos_fuscantarius
Hylaea_fasciaria
Idaea_aversata
Bombyx_mori
Deilephila_porcellus
Hemaris_fuciformis
Laothoe_populi
Mimas_tiliae
Manduca_sexta
Dendrolimus_punctatus
Clostera_curtula
Notodonta_dromedarius
Pheosia_gnoma
Pheosia_tremula
Phalera_bucephala
Euproctis_similis
Lymantria_monacha
Hypena_proboscidalis
Spilosoma_lubricipeda
Laspeyria_flexula
Schrankia_costaestrigalis
Amphipyra_tragopoginis
Craniophora_ligustri
Atethmia_centrago
Cosmia_trapezina
Noctua_fimbriata
Noctua_pronuba
Xestia_xanthographa
Ochropleura_plecta
Phlogophora_meticulosa
Hecatera_dysodea
Mamestra_brassicae
Mythimna_impura
Spodoptera_exigua
Spodoptera_frugiperda
Spodoptera_litura
Abrostola_tripartita
Autographa_gamma
Autographa_pulchrina
Tinea_trinotella

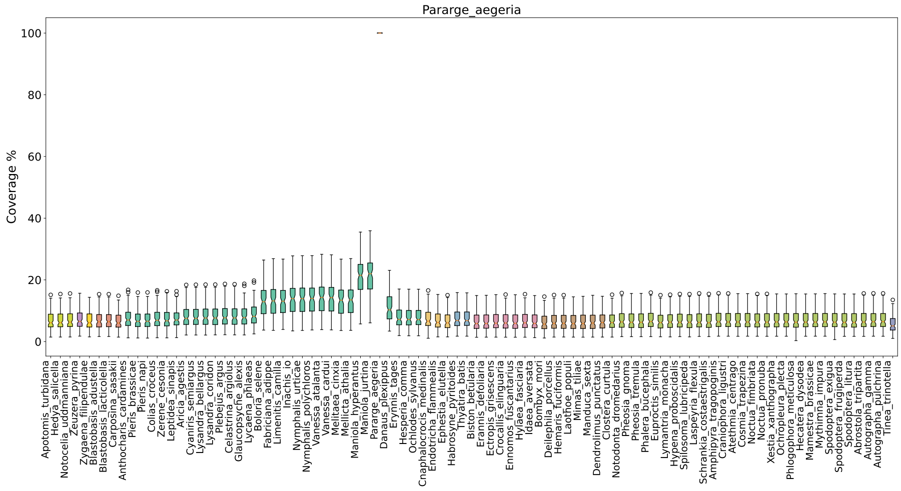
Pararge_aegeria
Coverage %
0
20
40
60
80
100
Apotomis_turbidana
Hedya_salicella
Notocelia_uddmanniana
Zeuzera_pyrina
Zygaena_filipendulae
Blastobasis_adustella
Blastobasis_lacticolella
Carposina_sasakii
Anthocharis_cardamines
Pieris_brassicae
Pieris_napi
Colias_croceus
Zerene_cesonia
Leptidea_sinapis
Aricia_agestis
Cyaniris_semiargus
Lysandra_bellargus
Lysandra_coridon
Plebejus_argus
Celastrina_argiolus
Glaucopsyche_alexis
Lycaena_phlaeas
Boloria_selene
Fabriciana_adippe
Limenitis_camilla
Inachis_io
Nymphalis_urticae
Nymphalis_polychloros
Vanessa_atalanta
Vanessa_cardui
Melitaea_cinxia
Mellicta_athalia
Maniola_hyperantus
Maniola_jurtina
Pararge_aegeria
Danaus_plexippus
Erynnis_tages
Hesperia_comma
Ochlodes_sylvanus
Cnaphalocrocis_medinalis
Endotricha_flammealis
Ephestia_elutella
Habrosyne_pyritoides
Thyatira_batis
Biston_betularia
Erannis_defoliaria
Ectropis_grisescens
Crocallis_elinguaria
Ennomos_fuscantarius
Hylaea_fasciaria
Idaea_aversata
Bombyx_mori
Deilephila_porcellus
Hemaris_fuciformis
Laothoe_populi
Mimas_tiliae
Manduca_sexta
Dendrolimus_punctatus
Clostera_curtula
Notodonta_dromedarius
Pheosia_gnoma
Pheosia_tremula
Phalera_bucephala
Euproctis_similis
Lymantria_monacha
Hypena_proboscidalis
Spilosoma_lubricipeda
Laspeyria_flexula
Schrankia_costaestrigalis
Amphipyra_tragopoginis
Craniophora_ligustri
Atethmia_centrago
Cosmia_trapezina
Noctua_fimbriata
Noctua_pronuba
Xestia_xanthographa
Ochropleura_plecta
Phlogophora_meticulosa
Hecatera_dysodea
Mamestra_brassicae
Mythimna_impura
Spodoptera_exigua
Spodoptera_frugiperda
Spodoptera_litura
Abrostola_tripartita
Autographa_gamma
Autographa_pulchrina
Tinea_trinotella

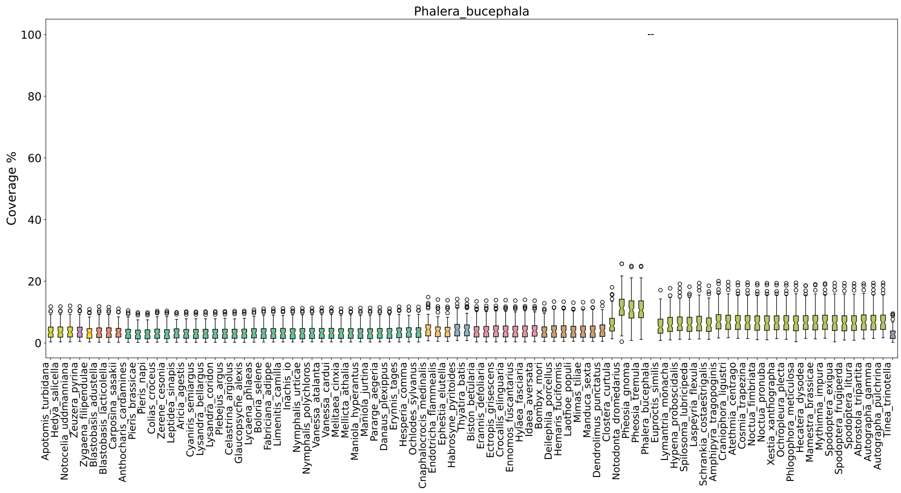
Phalera_bucephala
Coverage %
0
20
40
60
80
100
Apotomis_turbidana
Hedya_salicella
Notocelia_uddmanniana
Zeuzera_pyrina
Zygaena_filipendulae
Blastobasis_adustella
Blastobasis_lacticolella
Carposina_sasakii
Anthocharis_cardamines
Pieris_brassicae
Pieris_napi
Colias_croceus
Zerene_cesonia
Leptidea_sinapis
Aricia_agestis
Cyaniris_semiargus
Lysandra_bellargus
Lysandra_coridon
Plebejus_argus
Celastrina_argiolus
Glaucopsyche_alexis
Lycaena_phlaeas
Boloria_selene
Fabriciana_adippe
Limenitis_camilla
Inachis_io
Nymphalis_urticae
Nymphalis_polychloros
Vanessa_atalanta
Vanessa_cardui
Melitaea_cinxia
Mellicta_athalia
Maniola_hyperantus
Maniola_jurtina
Pararge_aegeria
Danaus_plexippus
Erynnis_tages
Hesperia_comma
Ochlodes_sylvanus
Cnaphalocrocis_medinalis
Endotricha_flammealis
Ephestia_elutella
Habrosyne_pyritoides
Thyatira_batis
Biston_betularia
Erannis_defoliaria
Ectropis_grisescens
Crocallis_elinguaria
Ennomos_fuscantarius
Hylaea_fasciaria
Idaea_aversata
Bombyx_mori
Deilephila_porcellus
Hemaris_fuciformis
Laothoe_populi
Mimas_tiliae
Manduca_sexta
Dendrolimus_punctatus
Clostera_curtula
Notodonta_dromedarius
Pheosia_gnoma
Pheosia_tremula
Phalera_bucephala
Euproctis_similis
Lymantria_monacha
Hypena_proboscidalis
Spilosoma_lubricipeda
Laspeyria_flexula
Schrankia_costaestrigalis
Amphipyra_tragopoginis
Craniophora_ligustri
Atethmia_centrago
Cosmia_trapezina
Noctua_fimbriata
Noctua_pronuba
Xestia_xanthographa
Ochropleura_plecta
Phlogophora_meticulosa
Hecatera_dysodea
Mamestra_brassicae
Mythimna_impura
Spodoptera_exigua
Spodoptera_frugiperda
Spodoptera_litura
Abrostola_tripartita
Autographa_gamma
Autographa_pulchrina
Tinea_trinotella

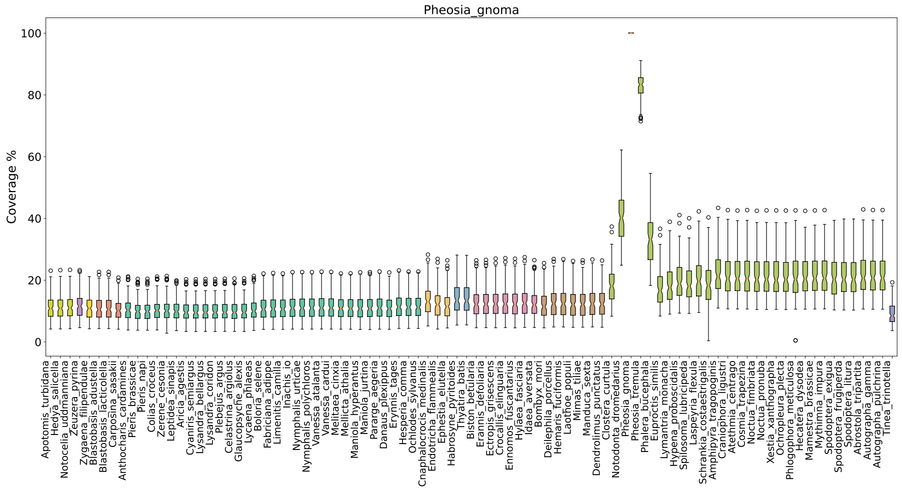
Pheosia_gnoma
Coverage %

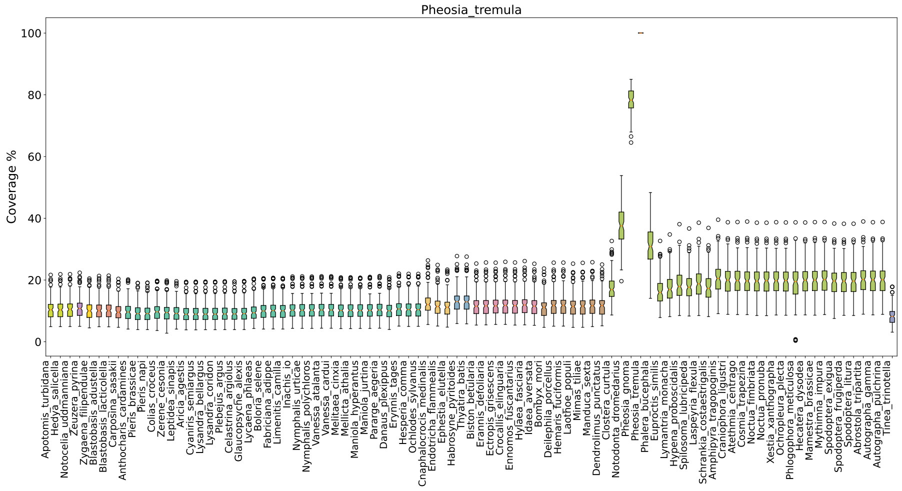
Pheosia_tremula
Coverage %

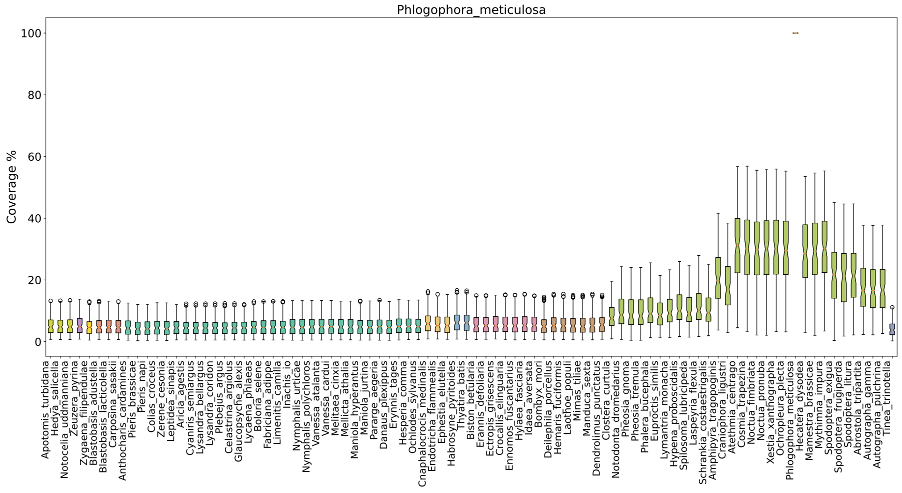
Phlogophora_meticulosa
Coverage %

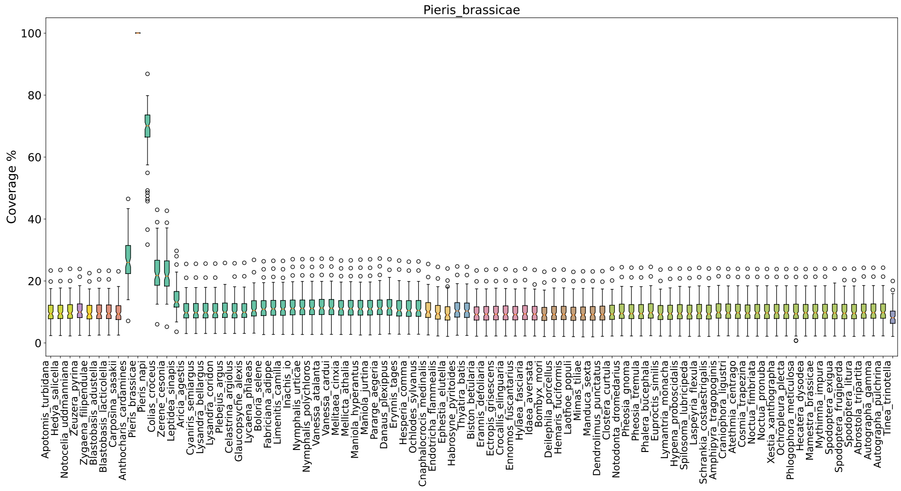
Pieris_brassicae
Coverage %
0
20
40
60
80
100
Apotomis_turbidana
Hedya_salicella
Notocelia_uddmanniana
Zeuzera_pyrina
Zygaena_filipendulae
Blastobasis_adustella
Blastobasis_lacticolella
Carposina_sasakii
Anthocharis_cardamines
Pieris_brassicae
Pieris_napi
Colias_croceus
Zerene_cesonia
Leptidea_sinapis
Aricia_agestis
Cyaniris_semiargus
Lysandra_bellargus
Lysandra_coridon
Plebejus_argus
Celastrina_argiolus
Glaucopsyche_alexis
Lycaena_phlaeas
Boloria_selene
Fabriciana_adippe
Limenitis_camilla
Inachis_io
Nymphalis_urticae
Nymphalis_polychloros
Vanessa_atalanta
Vanessa_cardui
Melitaea_cinxia
Mellicta_athalia
Maniola_hyperantus
Maniola_jurtina
Pararge_aegeria
Danaus_plexippus
Erynnis_tages
Hesperia_comma
Ochlodes_sylvanus
Cnaphalocrocis_medinalis
Endotricha_flammealis
Ephestia_elutella
Habrosyne_pyritoides
Thyatira_batis
Biston_betularia
Erannis_defoliaria
Ectropis_grisescens
Crocallis_elinguaria
Ennomos_fuscantarius
Hylaea_fasciaria
Idaea_aversata
Bombyx_mori
Deilephila_porcellus
Hemaris_fuciformis
Laothoe_populi
Mimas_tiliae
Manduca_sexta
Dendrolimus_punctatus
Clostera_curtula
Notodonta_dromedarius
Pheosia_gnoma
Pheosia_tremula
Phalera_bucephala
Euproctis_similis
Lymantria_monacha
Hypena_proboscidalis
Spilosoma_lubricipeda
Laspeyria_flexula
Schrankia_costaestrigalis
Amphipyra_tragopoginis
Craniophora_ligustri
Atethmia_centrago
Cosmia_trapezina
Noctua_fimbriata
Noctua_pronuba
Xestia_xanthographa
Ochropleura_plecta
Phlogophora_meticulosa
Hecatera_dysodea
Mamestra_brassicae
Mythimna_impura
Spodoptera_exigua
Spodoptera_frugiperda
Spodoptera_litura
Abrostola_tripartita
Autographa_gamma
Autographa_pulchrina
Tinea_trinotella

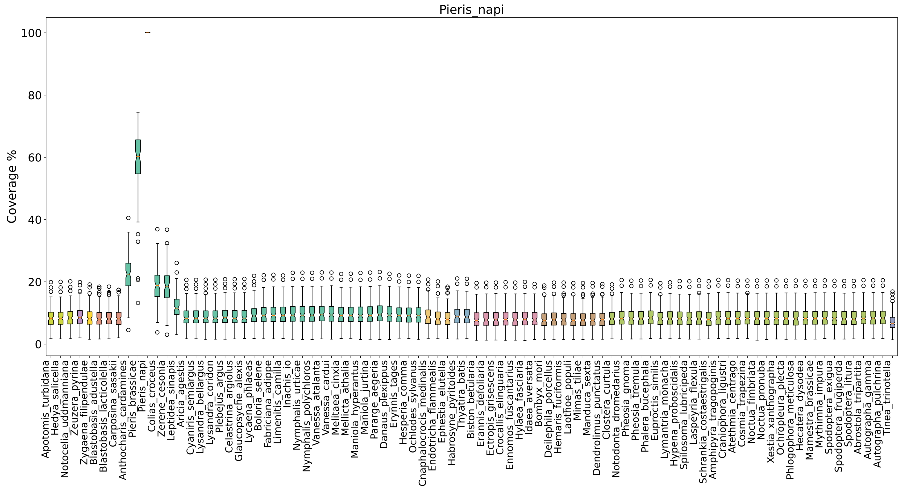
Pieris_napi
Coverage %
0
20
40
60
80
100
Apotomis_turbidana
Hedya_salicella
Notocelia_uddmanniana
Zeuzera_pyrina
Zygaena_filipendulae
Blastobasis_adustella
Blastobasis_lacticolella
Carposina_sasakii
Anthocharis_cardamines
Pieris_brassicae
Pieris_napi
Colias_croceus
Zerene_cesonia
Leptidea_sinapis
Aricia_agestis
Cyaniris_semiargus
Lysandra_bellargus
Lysandra_coridon
Plebejus_argus
Celastrina_argiolus
Glaucopsyche_alexis
Lycaena_phlaeas
Boloria_selene
Fabriciana_adippe
Limenitis_camilla
Inachis_io
Nymphalis_urticae
Nymphalis_polychloros
Vanessa_atalanta
Vanessa_cardui
Melitaea_cinxia
Mellicta_athalia
Maniola_hyperantus
Maniola_jurtina
Pararge_aegeria
Danaus_plexippus
Erynnis_tages
Hesperia_comma
Ochlodes_sylvanus
Cnaphalocrocis_medinalis
Endotricha_flammealis
Ephestia_elutella
Habrosyne_pyritoides
Thyatira_batis
Biston_betularia
Erannis_defoliaria
Ectropis_grisescens
Crocallis_elinguaria
Ennomos_fuscantarius
Hylaea_fasciaria
Idaea_aversata
Bombyx_mori
Deilephila_porcellus
Hemaris_fuciformis
Laothoe_populi
Mimas_tiliae
Manduca_sexta
Dendrolimus_punctatus
Clostera_curtula
Notodonta_dromedarius
Pheosia_gnoma
Pheosia_tremula
Phalera_bucephala
Euproctis_similis
Lymantria_monacha
Hypena_proboscidalis
Spilosoma_lubricipeda
Laspeyria_flexula
Schrankia_costaestrigalis
Amphipyra_tragopoginis
Craniophora_ligustri
Atethmia_centrago
Cosmia_trapezina
Noctua_fimbriata
Noctua_pronuba
Xestia_xanthographa
Ochropleura_plecta
Phlogophora_meticulosa
Hecatera_dysodea
Mamestra_brassicae
Mythimna_impura
Spodoptera_exigua
Spodoptera_frugiperda
Spodoptera_litura
Abrostola_tripartita
Autographa_gamma
Autographa_pulchrina
Tinea_trinotella

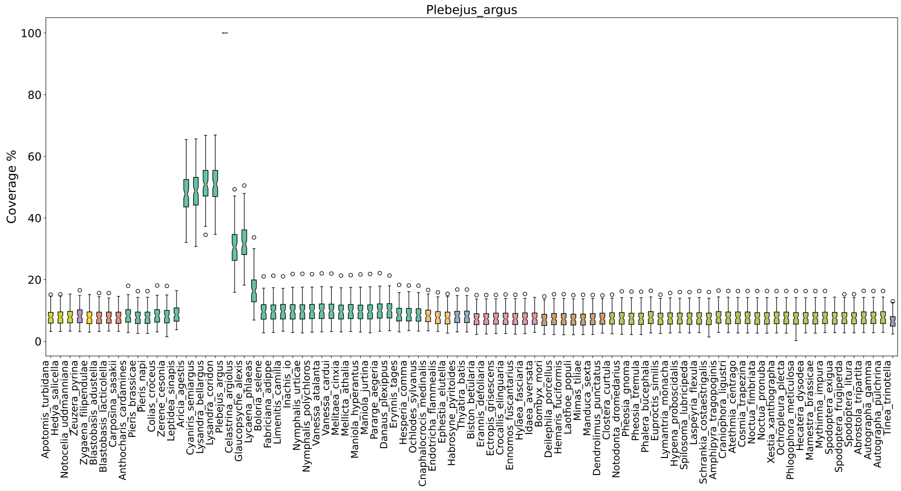
Plebejus_argus
Coverage %
0
20
40
60
80
100
Apotomis_turbidana
Hedya_salicella
Notocelia_uddmanniana
Zeuzera_pyrina
Zygaena_filipendulae
Blastobasis_adustella
Blastobasis_lacticolella
Carposina_sasakii
Anthocharis_cardamines
Pieris_brassicae
Pieris_napi
Colias_croceus
Zerene_cesonia
Leptidea_sinapis
Aricia_agestis
Cyaniris_semiargus
Lysandra_bellargus
Lysandra_coridon
Plebejus_argus
Celastrina_argiolus
Glaucopsyche_alexis
Lycaena_phlaeas
Boloria_selene
Fabriciana_adippe
Limenitis_camilla
Inachis_io
Nymphalis_urticae
Nymphalis_polychloros
Vanessa_atalanta
Vanessa_cardui
Melitaea_cinxia
Mellicta_athalia
Maniola_hyperantus
Maniola_jurtina
Pararge_aegeria
Danaus_plexippus
Erynnis_tages
Hesperia_comma
Ochlodes_sylvanus
Cnaphalocrocis_medinalis
Endotricha_flammealis
Ephestia_elutella
Habrosyne_pyritoides
Thyatira_batis
Biston_betularia
Erannis_defoliaria
Ectropis_grisescens
Crocallis_elinguaria
Ennomos_fuscantarius
Hylaea_fasciaria
Idaea_aversata
Bombyx_mori
Deilephila_porcellus
Hemaris_fuciformis
Laothoe_populi
Mimas_tiliae
Manduca_sexta
Dendrolimus_punctatus
Clostera_curtula
Notodonta_dromedarius
Pheosia_gnoma
Pheosia_tremula
Phalera_bucephala
Euproctis_similis
Lymantria_monacha
Hypena_proboscidalis
Spilosoma_lubricipeda
Laspeyria_flexula
Schrankia_costaestrigalis
Amphipyra_tragopoginis
Craniophora_ligustri
Atethmia_centrago
Cosmia_trapezina
Noctua_fimbriata
Noctua_pronuba
Xestia_xanthographa
Ochropleura_plecta
Phlogophora_meticulosa
Hecatera_dysodea
Mamestra_brassicae
Mythimna_impura
Spodoptera_exigua
Spodoptera_frugiperda
Spodoptera_litura
Abrostola_tripartita
Autographa_gamma
Autographa_pulchrina
Tinea_trinotella

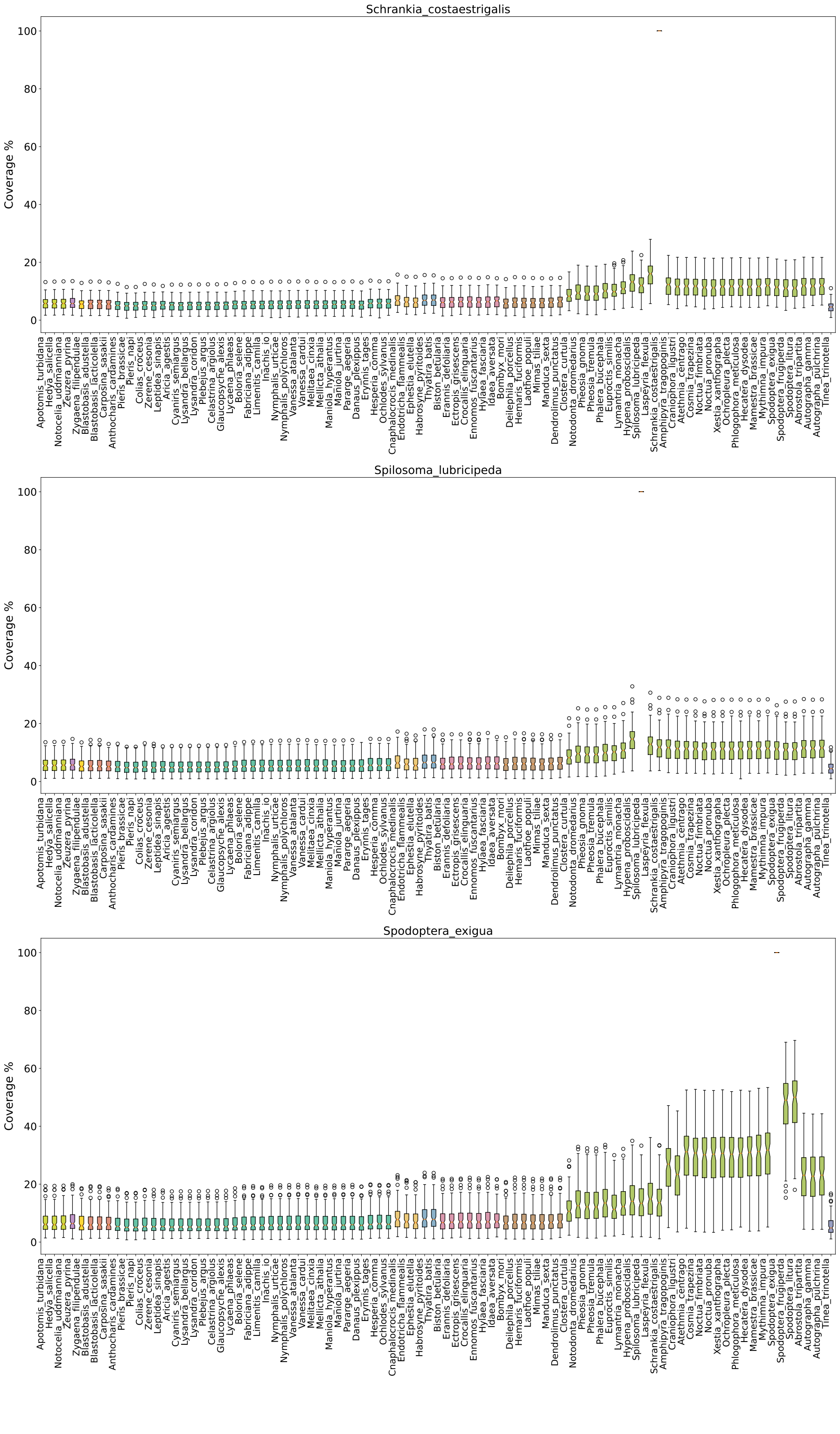
Schrankia_costaestrigalis
Spilosoma_lubricipeda
Spodoptera_exigua
Coverage %
0
20
40
60
80
100

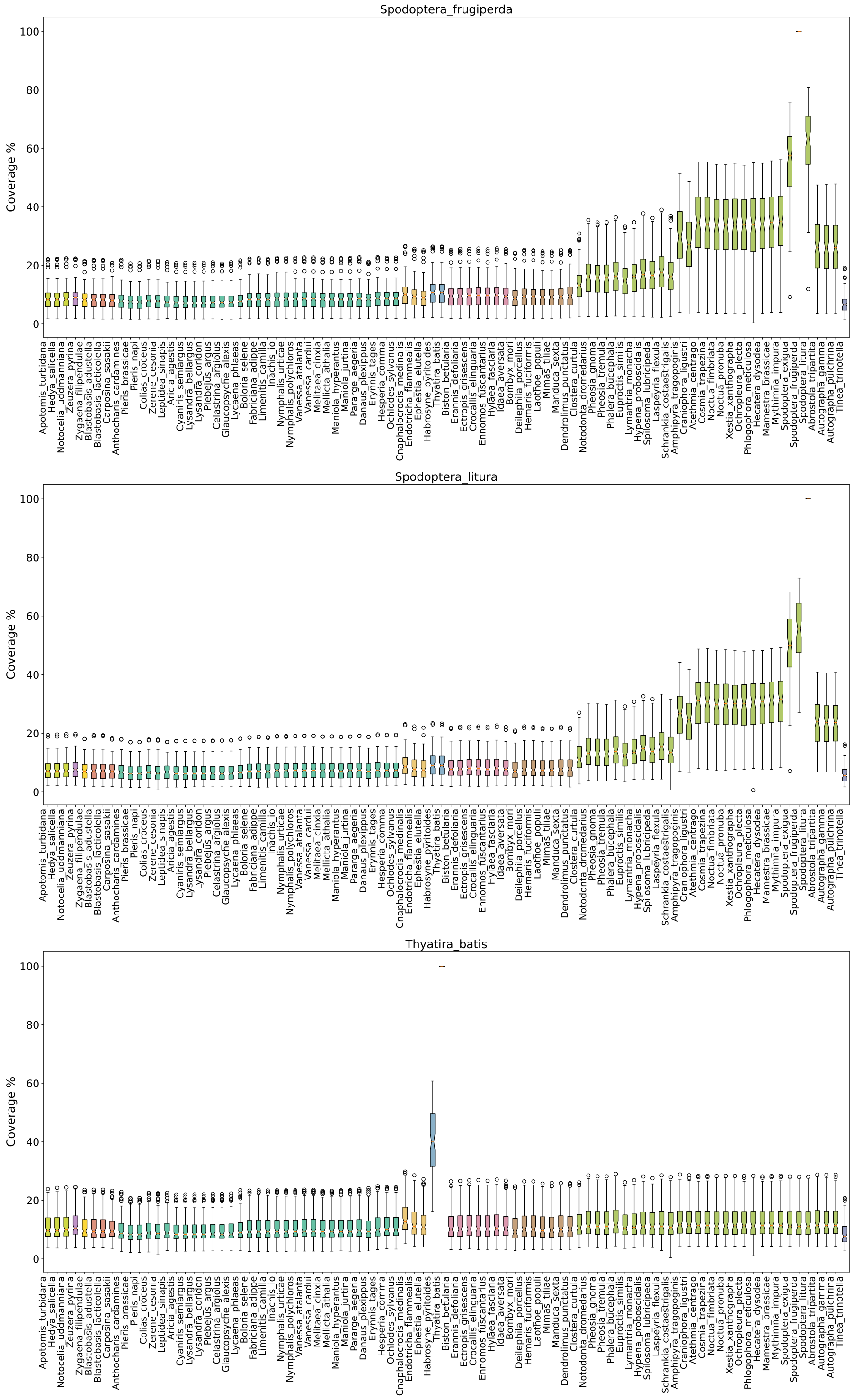
Spodoptera_frugiperda
Coverage %
Spodoptera_litura
Coverage %
Thyatira_batis
Coverage %

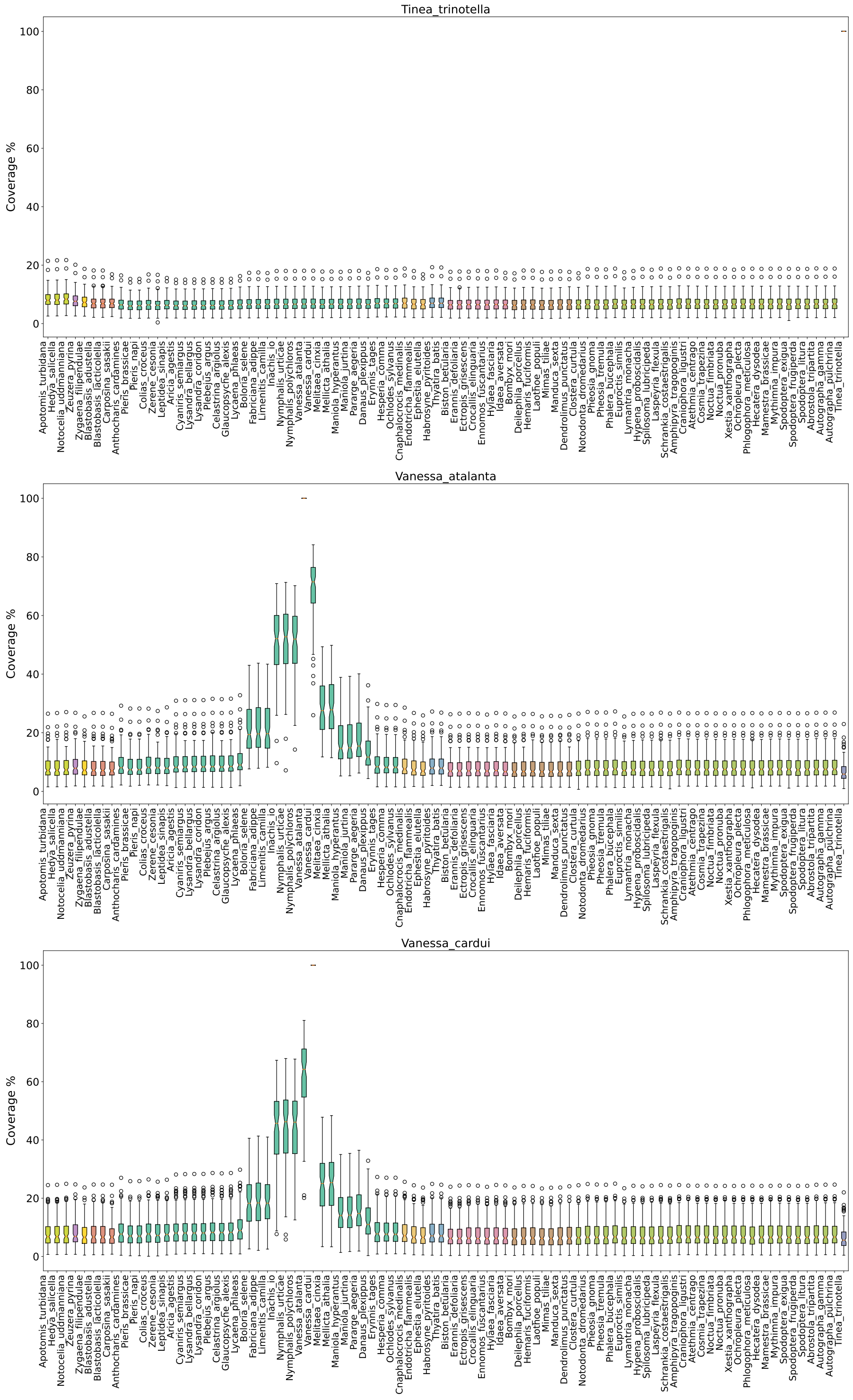
Tinea_trinotella
Coverage %
0 20 40 60 80 100
Apotomis_turbidana Hedya_salicella Notocelia_uddmanniana Zeuzera_pyrina Zygaena_filipendulae Blastobasis_adustella Blastobasis_lacticolella Carposina_sasakii Anthocharis_cardamines Pieris_brassicae Pieris_napi Colias_croceus Zerene_cesonia Leptidea_sinapis Aricia_agestis Cyaniris_semiargus Lysandra_bellargus Lysandra_coridon Plebejus_argus Celastrina_argiolus Glaucopsyche_alexis Lycaena_phlaeas Boloria_selene Fabriciana_adippe Limenitis_camilla Inachis_io Nymphalis_urticae Nymphalis_polychloros Vanessa_atalanta Vanessa_cardui Melitaea_cinxia Mellicta_athalia Maniola_hyperantus Maniola_jurtina Pararge_aegeria Danaus_plexippus Erynnis_tages Hesperia_comma Ochlodes_sylvanus Cnaphalocrocis_medinalis Endotricha_flammealis Ephestia_elutella Habrosyne_pyritoides Thyatira_batis Biston_betularia Erannis_defoliaria Ectropis_grisescens Crocallis_elinguaria Ennomos_fuscantarius Hylaea_fasciaria Idaea_aversata Bombyx_mori Deilephila_porcellus Hemaris_fuciformis Laothoe_populi Mimas_tiliae Manduca_sexta Dendrolimus_punctatus Clostera_curtula Notodonta_dromedarius Pheosia_gnoma Pheosia_tremula Phalera_bucephala Euproctis_similis Lymantria_monacha Hypena_proboscidalis Spilosoma_lubricipeda Laspeyria_flexula Schrankia_costaestrigalis Amphipyra_tragopoginis Craniophora_ligustri Atethmia_centrago Cosmia_trapezina Noctua_fimbriata Noctua_pronuba Xestia_xanthographa Ochropleura_plecta Phlogophora_meticulosa Hecatera_dysodea Mamestra_brassicae Mythimna_impura Spodoptera_exigua Spodoptera_frugiperda Spodoptera_litura Abrostola_tripartita Autographa_gamma Autographa_pulchrina Tinea_trinotella
Vanessa_atalanta
Coverage %
0 20 40 60 80 100
Apotomis_turbidana Hedya_salicella Notocelia_uddmanniana Zeuzera_pyrina Zygaena_filipendulae Blastobasis_adustella Blastobasis_lacticolella Carposina_sasakii Anthocharis_cardamines Pieris_brassicae Pieris_napi Colias_croceus Zerene_cesonia Leptidea_sinapis Aricia_agestis Cyaniris_semiargus Lysandra_bellargus Lysandra_coridon Plebejus_argus Celastrina_argiolus Glaucopsyche_alexis Lycaena_phlaeas Boloria_selene Fabriciana_adippe Limenitis_camilla Inachis_io Nymphalis_urticae Nymphalis_polychloros Vanessa_atalanta Vanessa_cardui Melitaea_cinxia Mellicta_athalia Maniola_hyperantus Maniola_jurtina Pararge_aegeria Danaus_plexippus Erynnis_tages Hesperia_comma Ochlodes_sylvanus Cnaphalocrocis_medinalis Endotricha_flammealis Ephestia_elutella Habrosyne_pyritoides Thyatira_batis Biston_betularia Erannis_defoliaria Ectropis_grisescens Crocallis_elinguaria Ennomos_fuscantarius Hylaea_fasciaria Idaea_aversata Bombyx_mori Deilephila_porcellus Hemaris_fuciformis Laothoe_populi Mimas_tiliae Manduca_sexta Dendrolimus_punctatus Clostera_curtula Notodonta_dromedarius Pheosia_gnoma Pheosia_tremula Phalera_bucephala Euproctis_similis Lymantria_monacha Hypena_proboscidalis Spilosoma_lubricipeda Laspeyria_flexula Schrankia_costaestrigalis Amphipyra_tragopoginis Craniophora_ligustri Atethmia_centrago Cosmia_trapezina Noctua_fimbriata Noctua_pronuba Xestia_xanthographa Ochropleura_plecta Phlogophora_meticulosa Hecatera_dysodea Mamestra_brassicae Mythimna_impura Spodoptera_exigua Spodoptera_frugiperda Spodoptera_litura Abrostola_tripartita Autographa_gamma Autographa_pulchrina Tinea_trinotella
Vanessa_cardui
Coverage %
0 20 40 60 80 100
Apotomis_turbidana Hedya_salicella Notocelia_uddmanniana Zeuzera_pyrina Zygaena_filipendulae Blastobasis_adustella Blastobasis_lacticolella Carposina_sasakii Anthocharis_cardamines Pieris_brassicae Pieris_napi Colias_croceus Zerene_cesonia Leptidea_sinapis Aricia_agestis Cyaniris_semiargus Lysandra_bellargus Lysandra_coridon Plebejus_argus Celastrina_argiolus Glaucopsyche_alexis Lycaena_phlaeas Boloria_selene Fabriciana_adippe Limenitis_camilla Inachis_io Nymphalis_urticae Nymphalis_polychloros Vanessa_atalanta Vanessa_cardui Melitaea_cinxia Mellicta_athalia Maniola_hyperantus Maniola_jurtina Pararge_aegeria Danaus_plexippus Erynnis_tages Hesperia_comma Ochlodes_sylvanus Cnaphalocrocis_medinalis Endotricha_flammealis Ephestia_elutella Habrosyne_pyritoides Thyatira_batis Biston_betularia Erannis_defoliaria Ectropis_grisescens Crocallis_elinguaria Ennomos_fuscantarius Hylaea_fasciaria Idaea_aversata Bombyx_mori Deilephila_porcellus Hemaris_fuciformis Laothoe_populi Mimas_tiliae Manduca_sexta Dendrolimus_punctatus Clostera_curtula Notodonta_dromedarius Pheosia_gnoma Pheosia_tremula Phalera_bucephala Euproctis_similis Lymantria_monacha Hypena_proboscidalis Spilosoma_lubricipeda Laspeyria_flexula Schrankia_costaestrigalis Amphipyra_tragopoginis Craniophora_ligustri Atethmia_centrago Cosmia_trapezina Noctua_fimbriata Noctua_pronuba Xestia_xanthographa Ochropleura_plecta Phlogophora_meticulosa Hecatera_dysodea Mamestra_brassicae Mythimna_impura Spodoptera_exigua Spodoptera_frugiperda Spodoptera_litura Abrostola_tripartita Autographa_gamma Autographa_pulchrina Tinea_trinotella

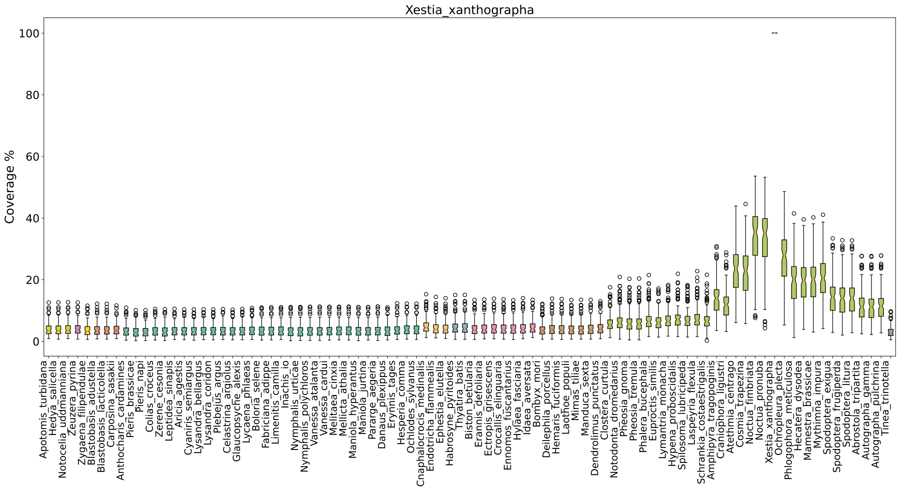
Xestia_xanthographa
Coverage %
0
20
40
60
80
100
Apotomis_turbidana
Hedya_salicella
Notocelia_uddmanniana
Zeuzera_pyrina
Zygaena_filipendulae
Blastobasis_adustella
Blastobasis_lacticolella
Carposina_sasakii
Anthocharis_cardamines
Pieris_brassicae
Pieris_napi
Colias_croceus
Zerene_cesonia
Leptidea_sinapis
Aricia_agestis
Cyaniris_semiargus
Lysandra_bellargus
Lysandra_coridon
Plebejus_argus
Celastrina_argiolus
Glaucopsyche_alexis
Lycaena_phlaeas
Boloria_selene
Fabriciana_adippe
Limenitis_camilla
Inachis_io
Nymphalis_urticae
Nymphalis_polychloros
Vanessa_atalanta
Vanessa_cardui
Melitaea_cinxia
Mellicta_athalia
Maniola_hyperantus
Maniola_jurtina
Pararge_aegeria
Danaus_plexippus
Erynnis_tages
Hesperia_comma
Ochlodes_sylvanus
Cnaphalocrocis_medinalis
Endotricha_flammealis
Ephestia_elutella
Habrosyne_pyritoides
Thyatira_batis
Biston_betularia
Erannis_defoliaria
Ectropis_grisescens
Crocallis_elinguaria
Ennomos_fuscantarius
Hylaea_fasciaria
Idaea_aversata
Bombyx_mori
Deilephila_porcellus
Hemaris_fuciformis
Laothoe_populi
Mimas_tiliae
Manduca_sexta
Dendrolimus_punctatus
Clostera_curtula
Notodonta_dromedarius
Pheosia_gnoma
Pheosia_tremula
Phalera_bucephala
Euproctis_similis
Lymantria_monacha
Hypena_proboscidalis
Spilosoma_lubricipeda
Laspeyria_flexula
Schrankia_costaestrigalis
Amphipyra_tragopoginis
Craniophora_ligustri
Atethmia_centrago
Cosmia_trapezina
Noctua_fimbriata
Noctua_pronuba
Xestia_xanthographa
Ochropleura_plecta
Phlogophora_meticulosa
Hecatera_dysodea
Mamestra_brassicae
Mythimna_impura
Spodoptera_exigua
Spodoptera_frugiperda
Spodoptera_litura
Abrostola_tripartita
Autographa_gamma
Autographa_pulchrina
Tinea_trinotella

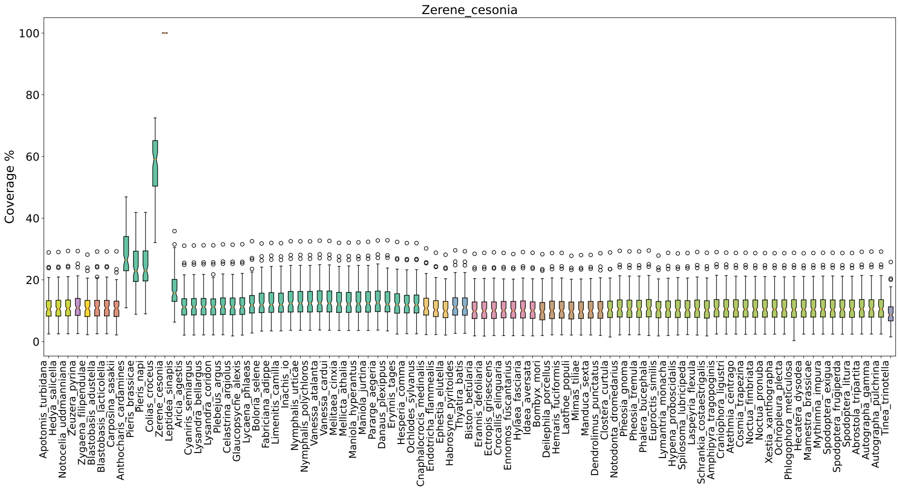
Zerene_cesonia
Coverage %
0
20
40
60
80
100
Apotomis_turbidana
Hedya_salicella
Notocelia_uddmanniana
Zeuzera_pyrina
Zygaena_filipendulae
Blastobasis_adustella
Blastobasis_lacticolella
Carposina_sasakii
Anthocharis_cardamines
Pieris_brassicae
Pieris_napi
Colias_croceus
Zerene_cesonia
Leptidea_sinapis
Aricia_agestis
Cyaniris_semiargus
Lysandra_bellargus
Lysandra_coridon
Plebejus_argus
Celastrina_argiolus
Glaucopsyche_alexis
Lycaena_phlaeas
Boloria_selene
Fabriciana_adippe
Limenitis_camilla
Inachis_io
Nymphalis_urticae
Nymphalis_polychloros
Vanessa_atalanta
Vanessa_cardui
Melitaea_cinxia
Mellicta_athalia
Maniola_hyperantus
Maniola_jurtina
Pararge_aegeria
Danaus_plexippus
Erynnis_tages
Hesperia_comma
Ochlodes_sylvanus
Cnaphalocrocis_medinalis
Endotricha_flammealis
Ephestia_elutella
Habrosyne_pyritoides
Thyatira_batis
Biston_betularia
Erannis_defoliaria
Ectropis_grisescens
Crocallis_elinguaria
Ennomos_fuscantarius
Hylaea_fasciaria
Idaea_aversata
Bombyx_mori
Deilephila_porcellus
Hemaris_fuciformis
Laothoe_populi
Mimas_tiliae
Manduca_sexta
Dendrolimus_punctatus
Clostera_curtula
Notodonta_dromedarius
Pheosia_gnoma
Pheosia_tremula
Phalera_bucephala
Euproctis_similis
Lymantria_monacha
Hypena_proboscidalis
Spilosoma_lubricipeda
Laspeyria_flexula
Schrankia_costaestrigalis
Amphipyra_tragopoginis
Craniophora_ligustri
Atethmia_centrago
Cosmia_trapezina
Noctua_fimbriata
Noctua_pronuba
Xestia_xanthographa
Ochropleura_plecta
Phlogophora_meticulosa
Hecatera_dysodea
Mamestra_brassicae
Mythimna_impura
Spodoptera_exigua
Spodoptera_frugiperda
Spodoptera_litura
Abrostola_tripartita
Autographa_gamma
Autographa_pulchrina
Tinea_trinotella

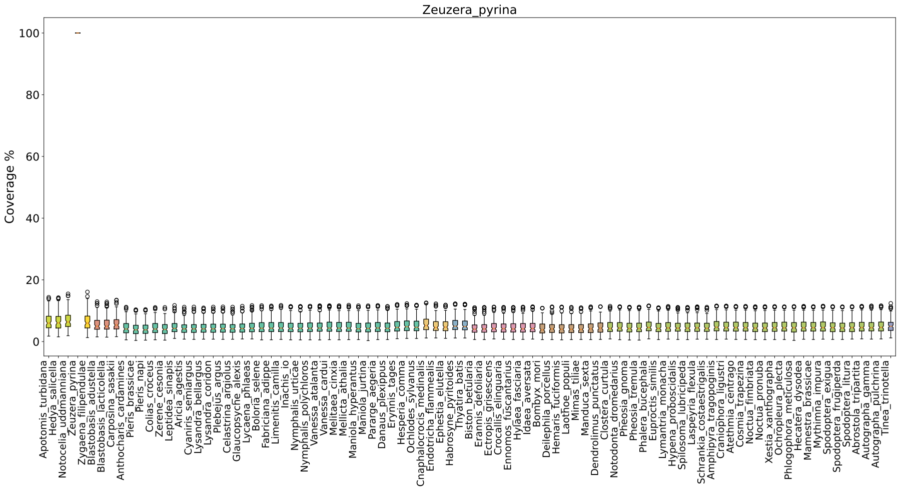
Zeuzera_pyrina
Coverage %
0
20
40
60
80
100
Apotomis_turbidana
Hedya_salicella
Notocelia_uddmanniana
Zeuzera_pyrina
Zygaena_filipendulae
Blastobasis_adustella
Blastobasis_lacticolella
Carposina_sasakii
Anthocharis_cardamines
Pieris_brassicae
Pieris_napi
Colias_croceus
Zerene_cesonia
Leptidea_sinapis
Aricia_agestis
Cyaniris_semiargus
Lysandra_bellargus
Lysandra_coridon
Plebejus_argus
Celastrina_argiolus
Glaucopsyche_alexis
Lycaena_phlaeas
Boloria_selene
Fabriciana_adippe
Limenitis_camilla
Inachis_io
Nymphalis_urticae
Nymphalis_polychloros
Vanessa_atalanta
Vanessa_cardui
Melitaea_cinxia
Mellicta_athalia
Maniola_hyperantus
Maniola_jurtina
Pararge_aegeria
Danaus_plexippus
Erynnis_tages
Hesperia_comma
Ochlodes_sylvanus
Cnaphalocrocis_medinalis
Endotricha_flammealis
Ephestia_elutella
Habrosyne_pyritoides
Thyatira_batis
Biston_betularia
Erannis_defoliaria
Ectropis_grisescens
Crocallis_elinguaria
Ennomos_fuscantarius
Hylaea_fasciaria
Idaea_aversata
Bombyx_mori
Deilephila_porcellus
Hemaris_fuciformis
Laothoe_populi
Mimas_tiliae
Manduca_sexta
Dendrolimus_punctatus
Clostera_curtula
Notodonta_dromedarius
Pheosia_gnoma
Pheosia_tremula
Phalera_bucephala
Euproctis_similis
Lymantria_monacha
Hypena_proboscidalis
Spilosoma_lubricipeda
Laspeyria_flexula
Schrankia_costaestrigalis
Amphipyra_tragopoginis
Craniophora_ligustri
Atethmia_centrago
Cosmia_trapezina
Noctua_fimbriata
Noctua_pronuba
Xestia_xanthographa
Ochropleura_plecta
Phlogophora_meticulosa
Hecatera_dysodea
Mamestra_brassicae
Mythimna_impura
Spodoptera_exigua
Spodoptera_frugiperda
Spodoptera_litura
Abrostola_tripartita
Autographa_gamma
Autographa_pulchrina
Tinea_trinotella

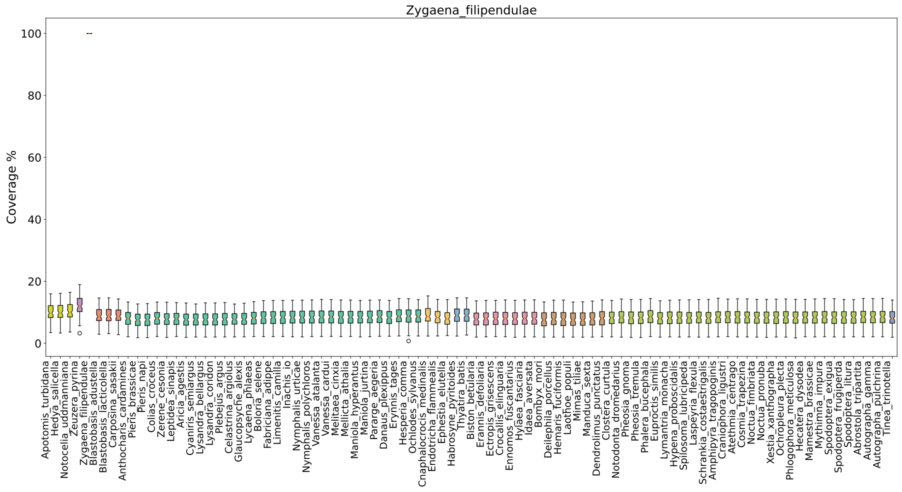


Figure S2: **Coverage of the cactus alignment**. Coverage is here expressed as the proportion of sites in species $A$ (i.e. query) that aligns to $n$ or more sites in species $B$ (i.e. target). Species are ordered on the x-axis according to the phylogenetic tree and are colored based on the superfamily they belong to. Coverage on the y-axis ranges from 0% to 100%.
