## Supplemental Figures 3, 4, 5 for "Lepidoptera genomics based on 88 chromosomal reference sequences informs population genetic parameters for conservation"

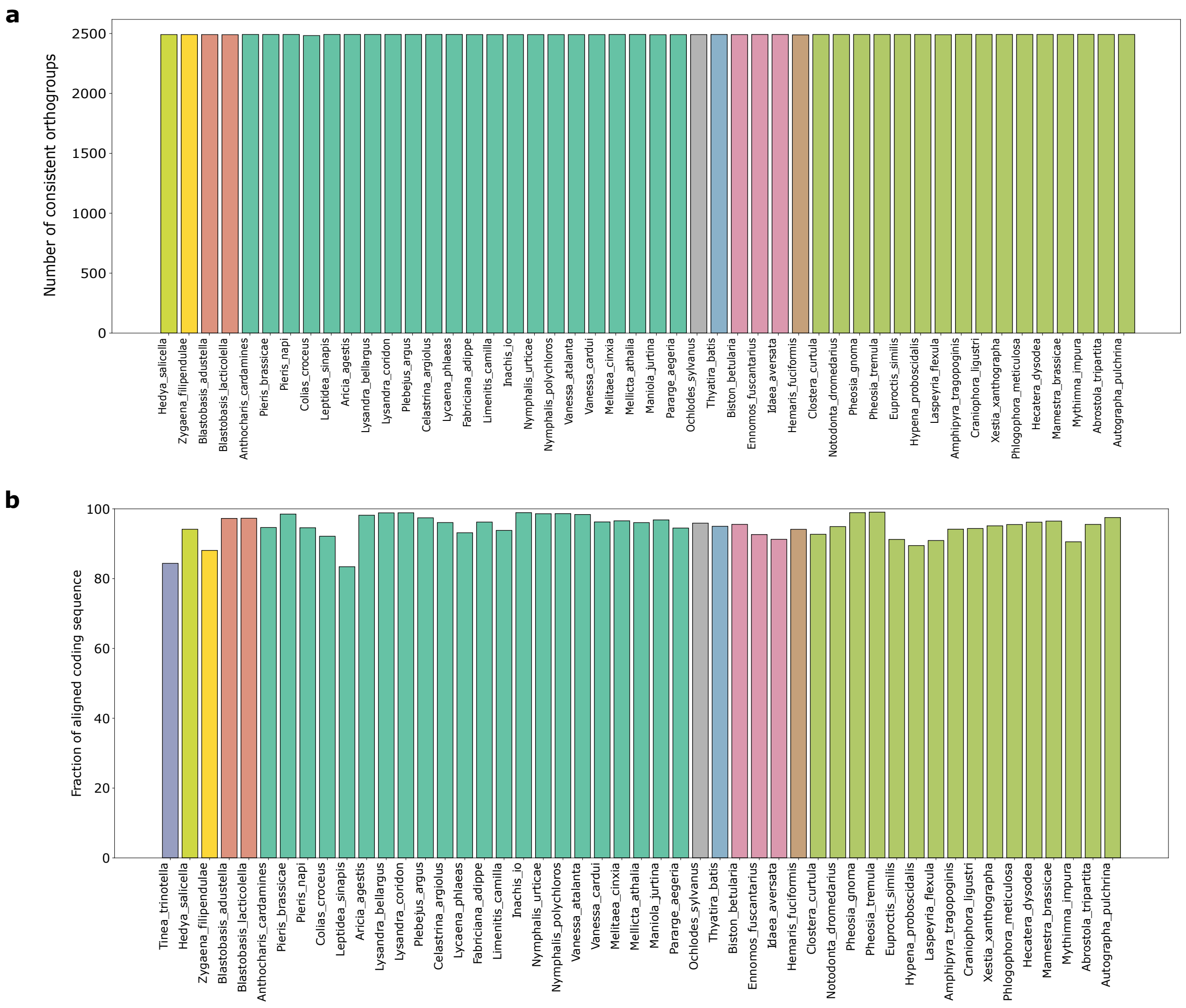


Figure S3: **Mapping consistency and alignment coverage**. **a.** Mapping consistency of single copy orthogroups. The y-axis shows the total number of correctly mapped (or consistent) single copy orthogroups as identified in OrthoFinder ($n$ = 2 501). **b.** Alignment coverage of protein-coding sequences. The y-axis shows the percentage of the protein coding sequence that is covered, in each species, by at least one other genome. In both bar plots, species are ordered on the x-axis according to the phylogenetic tree and are colored based on the superfamily they belong to.

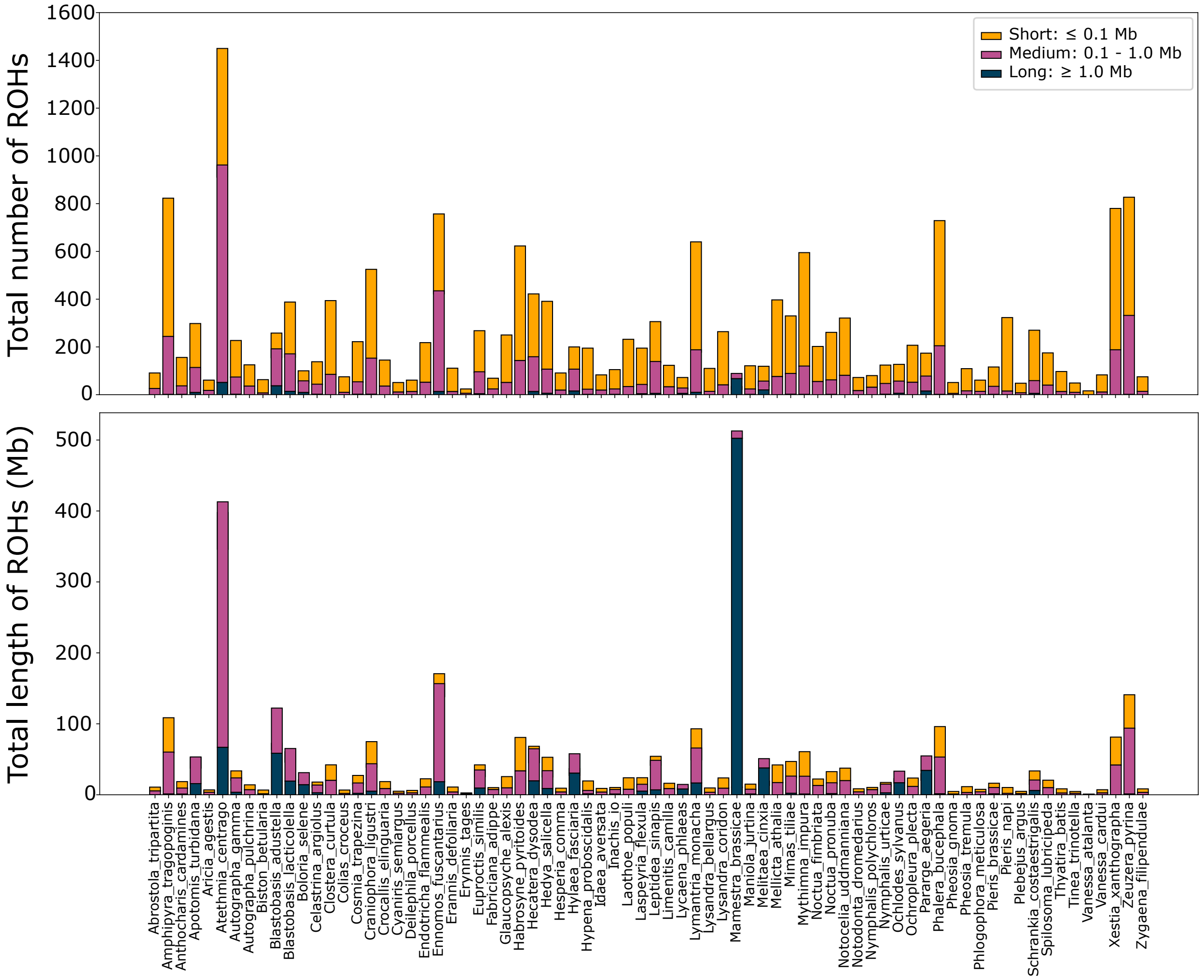


Figure S4: **Total number of runs of homozygosity (ROHs) and proportion of the genome covered by ROHs**. **a.** Number of ROHs belonging to the size classes small (<100 Kbp) medium (0.1 to 1 Mbp) and large (>1 Mbp) in each individual. **b.** Total size of the genome that is covered by a particular ROH size class in each individual.

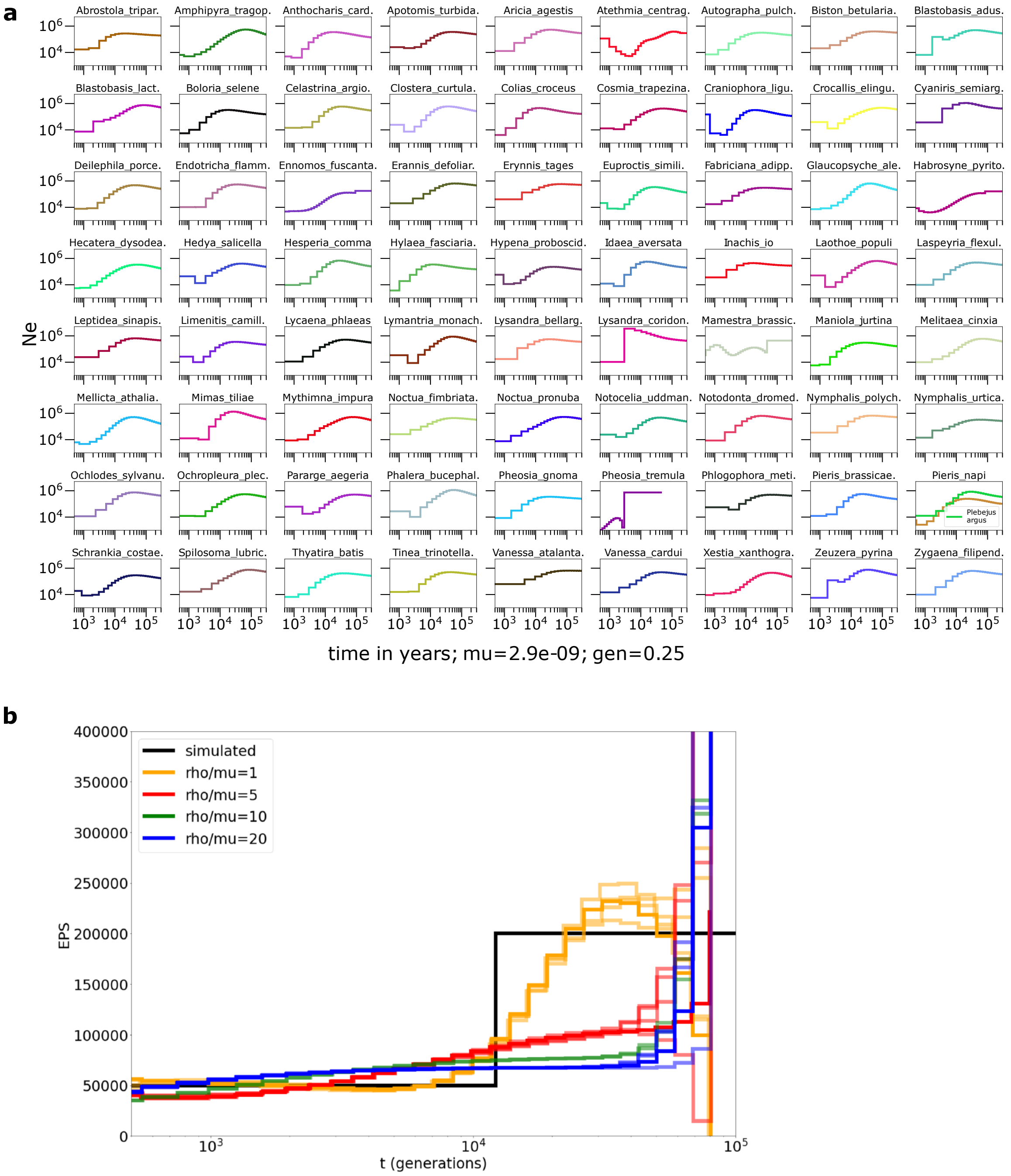


Figure S5: **PSMC estimates of the changes in effective population size over time for 74 species of Lepidoptera**. **a.** Changes in effective population size over time for each species, separately. **b.** Inference on a simulated population that undergoes a severe and prolonged bottleneck, with varying ratios of recombination to mutation. For all simulations, long term EPS N = 2e + 05, mu = 2.9e - 09, sequence length of 300Mb. Five replicates for each ratio are shown.
